# Engineering site-selective incorporation of fluorine into natural product analogs

**DOI:** 10.1101/2021.08.09.455754

**Authors:** S. Sirirungruang, O. Ad, T. M. Privalsky, S. Ramesh, J. L. Sax, H. Dong, E. E. K. Baidoo, B. Amer, C. Khosla, M. C. Y. Chang

## Abstract

While bioactive compounds are commonly derived both by human design as well as from living organisms, man-made and natural products typically display very different structural characteristics. As such, a longstanding goal in the discovery of new molecular function is to develop approaches to incorporate the advantageous elements of both groups of molecules, thereby expanding the molecular space accessible for this purpose. In this work, we report the engineering a fluorine-selective enzyme that can complement mutated acyltransferase (AT) domains of a modular polyketide synthase, which are the main determinants of the identity and location of substituents on polyketides, to produce different fluorinated regioisomers of the erythromycin precursor *in vitro*. We further show that by engineering cell uptake of fluorinated building blocks, we can control fluorine selectivity *in vivo* to produce selectively fluorinated polyketides using engineered *E. coli*. These results demonstrate that it is possible to introduce fluorine, a key synthetic design element for drug development, selectively into the scaffold of a complex natural product and produce these analogs by microbial fermentation.

## Main text

The blending of biological and chemical structure offers enormous potential for accessing new areas of structural space for discovery of function.^1-4^ While living organisms are particularly adept at constructing complex bioactive organic scaffolds, their use of the periodic table is limited when compared to human designs.^1, 5^ One such element is fluorine, which is found in only a handful of the >10^5^ known natural products, yet is highly prevalent as a functional design element amongst synthetic compounds.^6-8^ Continuing advances in fluorination methodology have enabled access to a growing range of fluorinated structures but its introduction into natural product scaffolds remains challenging.^9-13^ Here, we report a method to site-selectively incorporate fluorine *in vivo* into complex structures to produce regioselectively fluorinated polyketide natural products. By using engineered microbes, it is possible to produce elaborate fluorinated compounds by fermentation, offering the potential for expanding the identification and development of bioactive fluorinated small molecules.

Fluorine plays a pivotal role in molecular design due to its many unique properties, impacting a broad range of applications in agrochemicals to liquid crystals.^14, 15^ Indeed, the number of approved drugs containing fluorine has increased over an order of magnitude to reach 20-30% today.^6, 8^ However, the distinctive properties of fluorine that contribute to widespread usage also create problems in its introduction into target structures.^6^ As such, methods to site-selectively install fluorine into complex molecular architectures, especially under mild conditions remain of great interest. One especially challenging goal is to introduce fluorine selectively into natural products made by microbes, plants, and marine invertebrates. Given their size and complexity, natural products and their variants are particularly difficult to synthesize using purely chemical methods yet have evolved to contain structural features that can selectively target macromolecules ranging from proteins, nucleic acids, and carbohydrates. Given these attributes, natural products also make up a large proportion (33.5%) of approved drugs.^16^ Thus, combining natural and synthetic features in different ways could allow for increasing the accessible molecular space in which to search for new molecular functions.

We seek to expand the synthetic capabilities of living organisms to produce new molecules of interest to human society that combine key functional elements of both natural and synthetic compounds (**Figure 1a**). Using this approach, a multistep synthesis can potentially be telescoped to a single scalable growth process from simple universal building blocks such as glucose.^17-21^ In particular, we aim to develop a platform to introduce fluorine into complex drug-like scaffolds to allow synthesis of new organofluorines that would otherwise be difficult to access through chemical synthesis. We have focused on modular polyketides as they display diverse molecular structures that can be built with relatively predictable biosynthetic logic.^22, 23^ Given the hierarchical domain and module organization of the type I modular polyketide synthases (PKSs) that make these molecules, gene sequence and product structure are directly connected such that changes can be introduced site-selectively into the molecule by targeting mutations to the corresponding domain. We have now shown that a complete PKS system can be engineered to produce site-selectively fluorinated drug precursors *in vitro* and in an *Escherichia coli* production host using a modular platform.

**Fig. 1.**
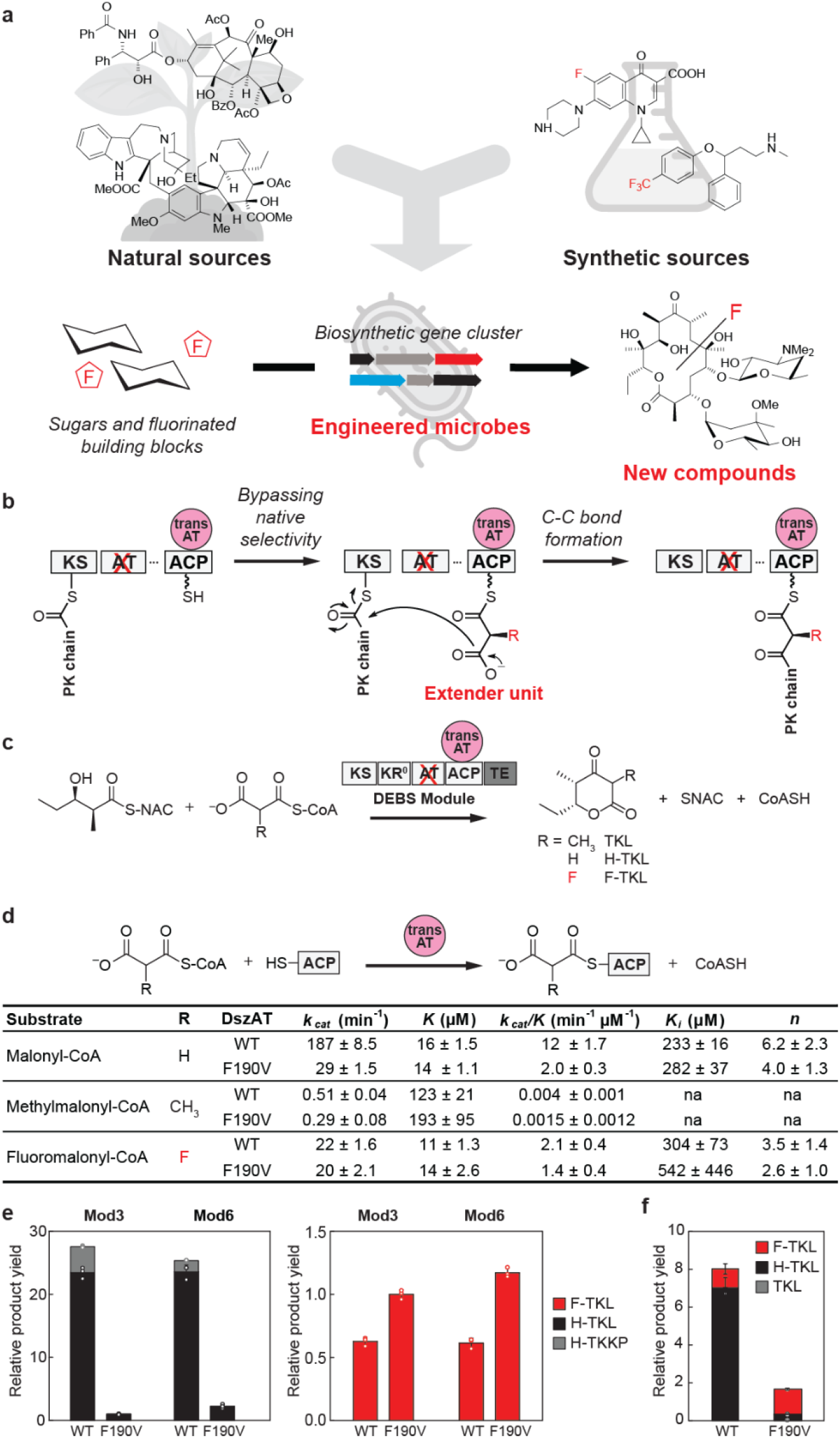
Merging biological and chemical strategies to expand the scope of fluorinated small molecule production. **(a)** By merging the ability of living systems to produce complex molecular architecture with design elements from synthetic chemistry we can expand the accessible scope of molecular space for discovery of new function. **(b)** Insertion of a non-native extender unit can be achieved by inactivating the *cis*-AT within a modular PKS and complementation in with a standalone *trans*-AT. This allows the native selectivity of the *cis*-AT, which is the dominant domain involved in selecting the identity of the extender unit, to be bypassed. **(c)** Formation of triketide lactones (TKLs) from N-acetylcysteamine thioester of (2*S*,3*R*)-2-methyl-3-hydroxypentanic acid (NDK-SNAC) and carboxyacyl-CoA extender unit catalyzed by single modular polyketide synthase construct. **(d)** Steady-state kinetic characterization of ACP transacylation catalyzed by DszAT variants with malonyl-CoA, methylmalonyl-CoA, and fluoromalonyl-CoA substrates. Table contains *k*_cat_, *K, k*_cat_/*K, K*_i_, and *n* calculated by non-linear curve fitting to the Hill equation with substrate inhibition (malonyl-CoA and fluoromalonyl-CoA, *K* = *K*_0.5_, *k*_cat_/*K* = *k*_cat_/*K*_0.5_) or Michaelis-Menten equation (methylmalonyl-CoA, *K* = *K*_M_, *k*_cat_/*K* = *k*_cat_/*K*_M_). Data are mean ± s.e. of three technical replicates. Error in *k*_cat_/*K*_M_ is obtained by propagation from the individual kinetic terms. **(e)** *In vitro* triketide lactone assay comparing wild-type and F190V DszAT under pure malonyl-CoA (left) and fluoromalonyl-CoA (right) extender unit conditions. Normalized representation of the amount of products arising from malonyl-CoA, H-TKL (single extension; black) and H-TTKP (double extension; grey), and from fluoromalonyl-CoA, F-TKL (red), are shown. Data are mean ± s.d. of technical replicates (n = 3). **(f**) *In vitro* triketide lactone assay comparing wild-type and F190V DszAT under mixed carboxyacyl-CoA extender unit condition. Normalized representation of the amount of H-TKL (black, malonyl-CoA product), TKL (grey, methylmalonyl-CoA product), and F-TKL (red, fluoromalonyl-CoA product) are shown. Data are mean ± s.d. of technical replicates (n = 3).

Of the >10,000 known polyketide natural products, most utilize malonyl coenzyme A (CoA; R=H) or methylmalonyl-CoA (R = methyl) as an extender unit. More unusual substituents on extender units are rare in native polyketide structures.^22, 24, 25^ Our goal is to expand the diversity of polyketides using a general platform to site-selectively introduce fluorine into full-length targets. Our previous work showed that we could use enzymes and cells to synthesize the fluoromalonyl-CoA (R = F) extender unit and incorporate it site-selectively into simple polyketide fragments.^26^ In this system, fluoromalonyl-CoA could be targeted to a particular module for incorporation using a complementation strategy,^27, 28^ where a separate *trans*-AT is used to load the fluorinated extender unit onto a PKS module where the *cis*-AT has been inactivated (**Figure 1b**).

Due to the small size of the polyketide fragments previously made,^26^ we were able to utilize a wild-type *trans*-AT from the disorazole pathway (DszAT) that natively charges the PKS with malonyl-CoA extender units.^29^ This method could be applied to simple model constructs of the 6-deoxyerythronolide B (6dEB) synthase (DEBS).^30^ However, to produce fluorinated variants of most complex polyketides, DszAT would need to be engineered to increase its selectivity for fluoromalonyl-CoA while greatly reducing activity on its native substrate malonyl-CoA, which is cellularly abundant.^31^ With this goal in mind, we initiated structure-guided protein engineering studies of DszAT to increase its selectivity towards fluoromalonyl-CoA. This simple strategy allows selective incorporation of fluorine in any module containing an inactivated *cis*-AT domain with the evolution of a single fluorine-selective *trans*-AT

The crystal structure of DszAT shows a phenylalanine residue at position 190 near the active site^32^ (**Extended Data Fig. 1a**). F190 is conserved across all malonyl-CoA selective AT domains from *cis*-AT PKS systems,^33, 34^ suggesting that it could be important for selectivity. L87 was found in the second sphere of the active site and hypothesized to be involved in substrate positioning with respect to the oxyanion hole. If the oxyanion hole is weakened, we reasoned that the effect should be less deleterious with fluorinated substrates as they are relatively activated for acyl transfer based on inductive effects (**Extended Data Fig. 1c**). Based on these observations, a DszAT saturation mutagenesis library was constructed at position F190 and combined with L87V or L87A mutations to reduce steric bulk. *E. coli* cells expressing the DszAT mutants were lysed and screened by triketide lactone (TKL) formation experiments^35^ (**Fig 1c, Extended Data Fig. 2**) by adding a purified DEBS module 3 construct with an inactivated AT domain and fused to thioesterase (TE) domain (Mod3_DEBS_+TE(AT^0^)) (**Extended Data Fig. 3a**). Select mutants that showed increased relative production of fluorinated TKL (F-TKL) were purified and investigated *in vitro* with malonyl-CoA, methylmalonyl-CoA, and fluoromalonyl-CoA substrates (**Extended Data Fig. 3b**).

These preliminary screens indicated that F190V DszAT mutant was more selective towards fluoromalonyl-CoA compared to wild-type enzyme. Steady state kinetic characterization of the purified mutant further shows that transacylation of an ACP partner is maintained with fluoromalonyl-CoA but significantly decreased with malonyl- and methylmalonyl-CoA (**Fig. 1d, Extended Data Fig. 4ab**). Interestingly, unproductive hydrolysis of fluoromalonyl-CoA is also reduced in the mutant (**Extended Data Fig. 4c**). Thus, F190V DszAT attains high fluoromalonyl-CoA selectivity by both suppressing malonyl-CoA turnover and fluoromalonyl-CoA hydrolysis.

With this mutant in hand, we then tested TKL formation *in vitro* with single or mixed carboxyacyl-CoA extender units (R = H, F, Me). When malonyl-CoA was provided as the sole extender unit, the production of desmethyl triketide lactone (H-TKL) by DszAT F190V was found to be reduced by an order of magnitude compared to wild-type (**Fig. 1e**). Moreover, DszAT F190V was able to moderately increase F-TKL production compared to that produced by the wild-type enzyme when fluoromalonyl-CoA was provided. Next, the engineered enzyme was subjected to mixed extender unit conditions, which would be required to produce a complex polyketide product utilizing malonyl-CoA, methylmalonyl-CoA, or both (**Fig. 1f**). When presented with an equimolar mixture of all three extender units, wild-type DszAT produces 7-fold more H-TKL than F-TKL. In comparison, F-TKL production is maintained with DszAT F190V with a 20-fold reduction in H-TKL leading to F-TKL as the dominant product (79 ± 5%). Taken together, these results show that DszAT F190V demonstrates the desired selectivity needed for a fluorine-selective *trans*-AT reagent that can be used in an environment with a complex carboxyacyl-CoA extender unit pool.

To test the ability of DszAT F190V for preparation of complex polyketides, we decided to focus on the DEBS system, which has been fully reconstituted *in vitro*.^36^ The polyketide product of the DEBS pathway, 6dEB, contains 21 carbons and 10 stereocenters and is produced through a series of nearly 30 enzymatic steps from 1 propionyl-CoA starter unit and 6 methylmalonyl-CoA extender units^30^ (**Fig. 2a**). The DEBS PKS itself consists of three proteins (DEBS1-3), each about 300 kDa in monomeric mass and together comprise 6 modules and a loading didomain. Hence the introduction of a single point mutations into one of the DEBS proteins is complicated by their large size as well as homology with the other module encoded on the same protein. Thus, a strategy was developed to selectively inactivate an AT domain by first replacing a section with an antibiotic marker so that the mutated domain can cleanly be reintroduced. Full sequencing is also needed to ensure that sequences are not lost to recombination.

To produce 2-fluoro-2-desmethyl-6dEB, the AT domain of module 6 was inactivated (Mod6 AT^0^ DEBS) and complemented with either wild-type or F190V DszAT. When fluoromalonyl-CoA and *trans*-AT were supplied to the system, a mass peak corresponding to a monofluorinated desmethyl 6dEB analog was observed (**Fig. 2b, Extended Data Fig. 6a**). This product was absent when there was no active site mutation in the AT domain of module 6 of the DEBS system. These results demonstrate the robustness of the complementation strategy to selectively incorporate fluorine using a multi-modular PKS in the presence of a mixed extender unit pool.

**Fig. 2.**
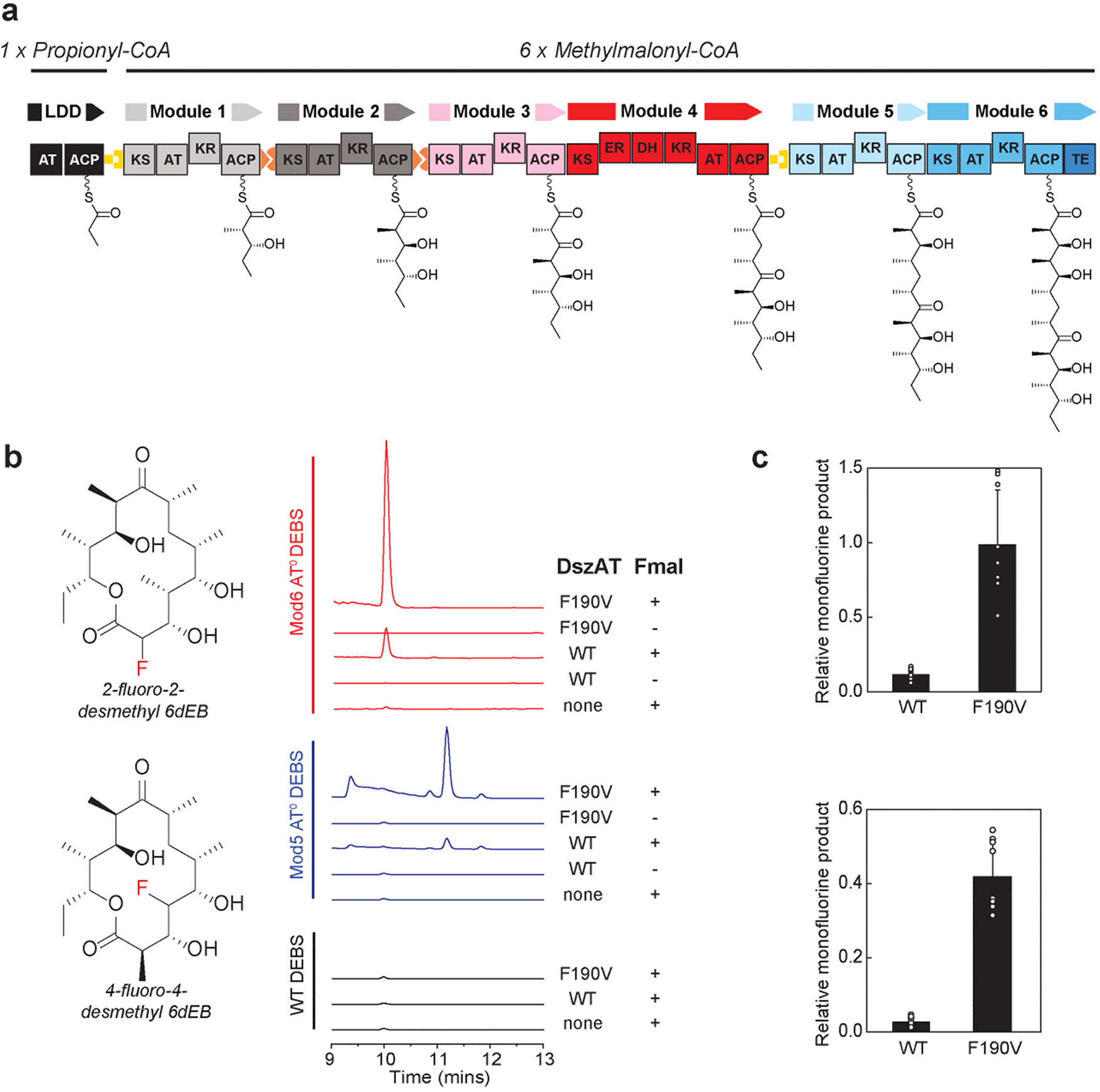
*In vitro* generation of regioselectively-fluorinated 6dEB analog. **(a)** The 6-deoxyerythronolide B synthase (DEBS) produces the aglycone precursor of antibiotic erythromycin and consists of a loading didomain and six modules, assembling one propionyl-CoA and six methylmalonyl-CoA molecules into product consisting of 21 carbons and 10 stereocenters.**(b)** The fluorine substituent was incorporated regioselectively to produce 2-fluoro-2-desmethyl 6dEB and 4-fluoro-4-desmethyl 6dEB analogs by *in vitro* reconstitution of Mod6 AT^0^ DEBS and Mod5 AT^0^ DEBS, respectively. Reactions contained 2 mM methylmalonate and when indicated 10 mM fluoromalonate. Extracted ion chromatograms shown are representative of at least three independent replicates. **(c)** System selectivity of fluoromalonyl-CoA incorporation by Mod6 AT^0^ DEBS (top) and Mod5 AT^0^ DEBS (bottom) by WT and F190V DszAT. Relative monofluorine product was calculated from the ratio between integrated extracted ion counts for monofluorinated desmethyl 6dEB analog to 6dEB of each replicate. Data are mean ± s.d. of nine replicates.

To confirm regiospecific incorporation of fluoromalonyl-CoA, fragmentation analysis was used to study the substitution pattern of the observed fluorine-containing desmethyl 6dEB analog produced by both F190V and WT DszAT (**Extended Data Fig. 7**,**8**). The fragmentation patterns obtained were consistent with the fluoromalonyl-CoA extender unit being incorporated by module 6, suggesting that the observed fluorine-containing desmethyl 6dEB analog was the expected target, 2-fluoro-2-desmethyl 6dEB^37^ (**Extended Data Fig. 6bc**). When comparing the system selectivity, the relative amount of 2-fluoro-2-desmethyl 6dEB analog to 6dEB was found higher in the system containing the engineered DszAT than in the system containing wild-type DszAT (**Fig. 2c, Extended Data Fig. 6d**).

Next, the AT domain of module 5 was inactivated (Mod5 AT^0^ DEBS) with the intention of producing 4-fluoro-4-desmethyl 6dEB to show that normal chain elongation could occur after the incorporation of a fluoromalonyl-CoA extender unit. Indeed, we were able to again detect the mass peak corresponding to a monofluorinated desmethyl 6dEB when Mod5 AT^0^ DEBS was complemented with WT or F190V DszAT (**Fig. 2b, Extended Data Fig. 9a**). Similar to the production of 2-fluoro-2-desmethyl 6dEB, the production of this compound was dependent on fluoromalonyl-CoA, *trans*-AT, and the AT domain active site mutation. Notably, the retention time of this compound differed from the product observed with Mod6 AT^0^ DEBS, suggesting that a different regioisomer was made. Fragmentation analysis is consistent with the monofluorinated desmethyl 6dEB analog arising from fluoromalonyl-CoA incorporation by module 5, producing 4-fluoro-4-desmethyl 6dEB (**Extended Data Fig 9bc**). Again, an increase in system selectivity towards the fluorine-containing analog was observed when F190V DszAT was used (**Fig. 2c, Extended Data Fig. 9d**).

With site-selective fluorine incorporation achieved *in vitro*, we sought to develop an *in vivo* cellular production system for fluorine-containing polyketide molecules. While *in vitro* experiments allowed us highly controlled environments for enzymatic production, they can be challenging to scale up for larger scale processes compared to fermentation in engineered microbes. However, the use of an unusual element such as fluorine yields new challenges in the precise control of the intracellular concentration and flux of system components as well as issues in toxicity and cross reactivity.

Previous work showed that feeding fluoromalonate to the culture medium of *E. coli* cells expressing DEBS_Mod6_ leads to detectable production of F-TKL.^26^ We thus hypothesized that increasing the availability of fluoromalonyl-CoA in the cell, as well as using a complementation strategy, would allow us to increase the yield of F-TKL and produce fluorine-containing polyketides of increasing complexity in living cells. Toward the first goal of increasing the intracellular pool of fluoromalonyl-CoA, both a malonate transporter and a malonyl-CoA synthetase (*Rhodopseudomonas palustris* MatB) were included (**Fig. 3a**). We then constructed a four-gene, three-plasmid system for *in vivo* biosynthesis of F-TKL from fluoromalonate as the fluorine source, containing a malonate transporter, MatB, the *trans*-AT, and either Mod3_DEBS_+TE(AT^0^) or Mod6_DEBS_+TE(AT^0^) proteins^38^ (**Fig. 3b**).

**Fig. 3.**
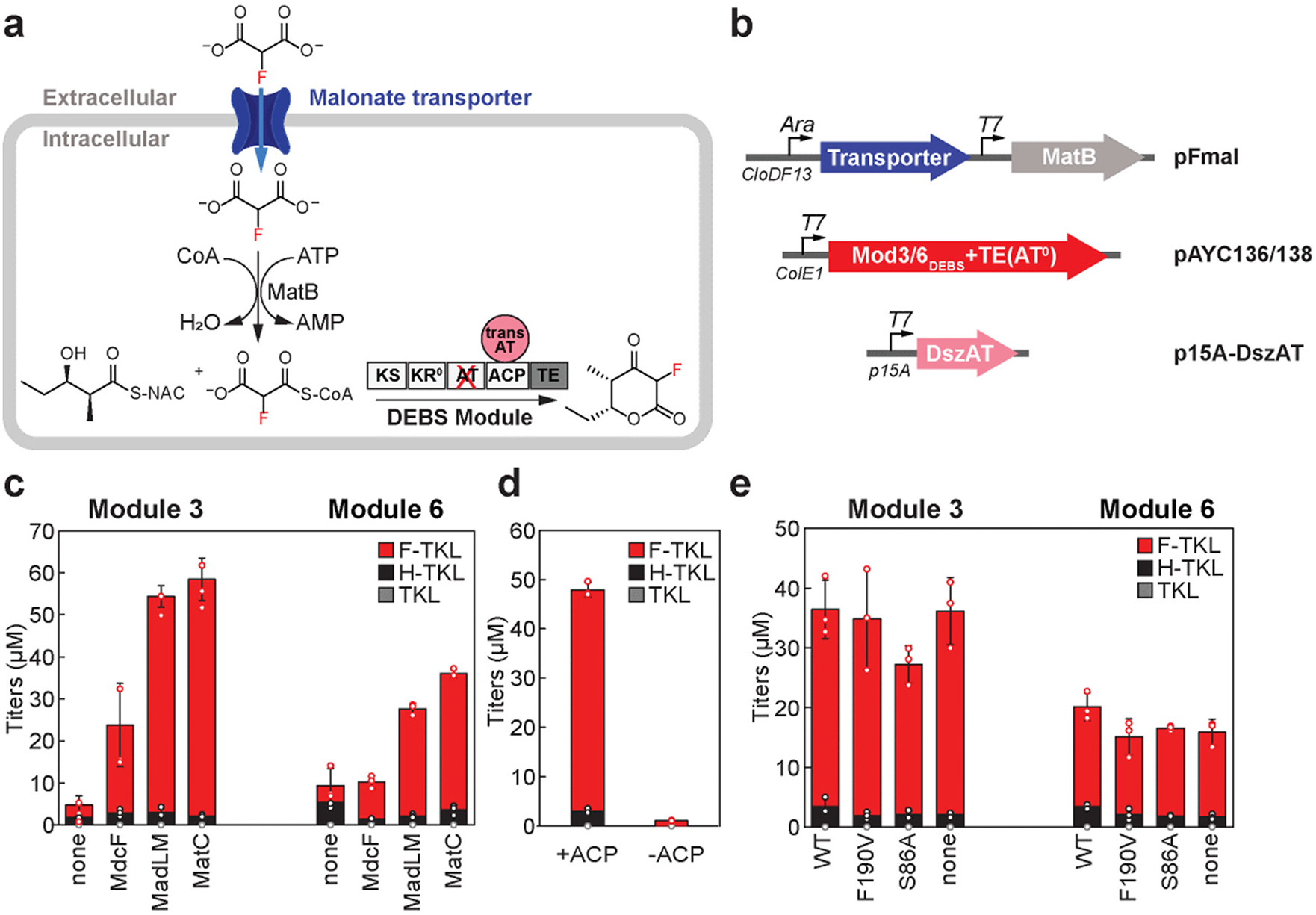
Cellular production of fluorinated polyketide model compounds by engineered *E. coli.* **(a)** Complementation strategy for fluorinated polyketide fragments production in *E. coli*. In addition to the PKS and the *trans*-AT, the cellular production system requires a specialized transport system to import fluoromalonate into the intracellular space and activate to the fluoromalonyl-CoA extender unit. Cells natively maintain a high level of ATP, which provides the necessary ATP regeneration. **(b)** Engineered pathway for fluorinated triketide lactone (F-TKL) production in *E. coli*. The system includes three plasmids with compatible origins and antibiotic markers, which together encode for four pathway proteins. pFmal (cloDF13 origin, Sp^R^) series encode for a malonate transporter and malonyl-CoA synthetase MatB. pAYC136 and 138 (ColE1 origin, Cb^R^) encode for Mod3_DEBS_+TE(AT^0^) and Mod6_DEBS_+TE(AT^0^), respectively. p15A-DszAT (p15A origin, Cm^R^) encodes for the DszAT variant. **(c)** Production titers of TKLs showing the contribution of malonate transporters (*Klebsiella pneumoniae* MdcF, *Pseudomonas fluorescens* MadLM, and *Streptomyces coelicolor* MatC) when NDK-SNAC and fluoromalonate were provided to production cultures. Data are mean ± s.d. of three biological replicates. **(d)** Production titers of TKLs by *E. coli* expressing pathway with or without an active ACP domain showing the requirement for the domain in F-TKL production. Data are mean ± s.d. of three biological replicates. **(e)** Production titers of TKLs by *E. coli* expressing pathway with an active (WT and F190V) or an inactive (S86A and no) *trans-*AT Data are mean ± s.d. of three biological replicates.

Comparison of the three malonate transporters MdcF, MadLM, and MatC showed that all significantly increase the yield of F-TKL produced by both Mod3_DEBS_+TE(AT^0^) and Mod6_DEBS_+TE(AT^0^) (**Fig. 3c**). While MdcF only demonstrated a moderate effect, expression of either MadLM or MatB increased F-TKL titer up to 60 μM with MadLM showing less dependence on growth phase.^39^ To test that F-TKL formation was dependent on proper acylation of the ACP, the ACP domain was inactivated in the module 3 construct, Mod3_DEBS_+TE(AT^0^ ACP^0^). The mutation in the ACP led to a virtual abolishment of F-TKL production, suggesting that production proceeds through the expected covalently tethered intermediate^40^ that is competent for multiple chain extension reactions (**Fig 3d**). Interestingly, production appears to be independent of a functional DszAT, suggesting that an endogenous host protein may also be competent at complementation (**Fig 3e**). Since *E. coli* does contain a malonyl-CoA-ACP transacylase (FabD) in the fatty acid synthase pathway, this enzyme could serve as the endogenous *trans*-AT. Despite its low sequence similarity to DszAT and DEBS AT domains, FabD does contain similar structural features (**Extended Data Fig. 10ab**) and is competent to acylate the DEBS module 6 ACP domain as shown both by steady-state kinetic analysis (**Extended Data Fig. 10c**) and TKL formation (**Extended Data Fig. 10d**). These results show that *in vivo* fluorinated polyketide yield and selectivity is likely governed by the high intracellular concentration of fluoromalonyl-CoA in the engineered host (**Extended Data Fig. 11**).

Based on these findings we adapted our engineered system to incorporate *E. coli* FabD as the source of *trans*-AT activity. We tested the ability of native *E. coli* metabolism to incorporate fluoromalonyl-CoA into complex polyketide products using a high-yielding *E. coli* 6dEB production model system^41^ (**Fig. 4a**). We then sought to produce regioselectively fluorinated 6dEB analogs *in vivo* with both Mod5 AT^0^ and Mod6 AT^0^ DEBS. In the absence of fluoromalonate, the system predominantly produces a single species of desmethyl-6dEB with fragmentation patterns consistent with regioselective production of 4-desmethyl-6dEB and 2-desmethyl-6dEB^37^ as expected from the Mod5 AT^0^ and Mod6 AT^0^ DEBS constructs, respectively (**Extended Data Fig. 12**). This analysis supports that the observed desmethyl-6dEB species were a result of the module with inactivated AT domain incorporating malonyl-CoA instead of methylmalonyl-CoA. Upon expression of MatB and MadLM and feeding of fluoromalonate to produce fluoromalonyl-CoA, we found that both Mod5 AT^0^ and Mod6 AT^0^ DEBS each produced a distinct monofluorinated desmethyl 6dEB analog (**Fig. 4bc**). Fragmentation, NMR analysis, and comparison to products produced *in vitro* supported the assignment to the expected regioisomers for each construct (Mod5 AT^0^ DEBS, 4-fluoro-4-desmethyl-6dEB; Mod6 AT^0^ DEBS, 2-fluoro-2-desmethyl-6dEB; **Extended Data Fig. 13**). Taken together, these results show that regiospecific introduction of fluorine substitution can be achieved in complex and full-length polyketide natural products in engineered living cells.

**Fig. 4.**
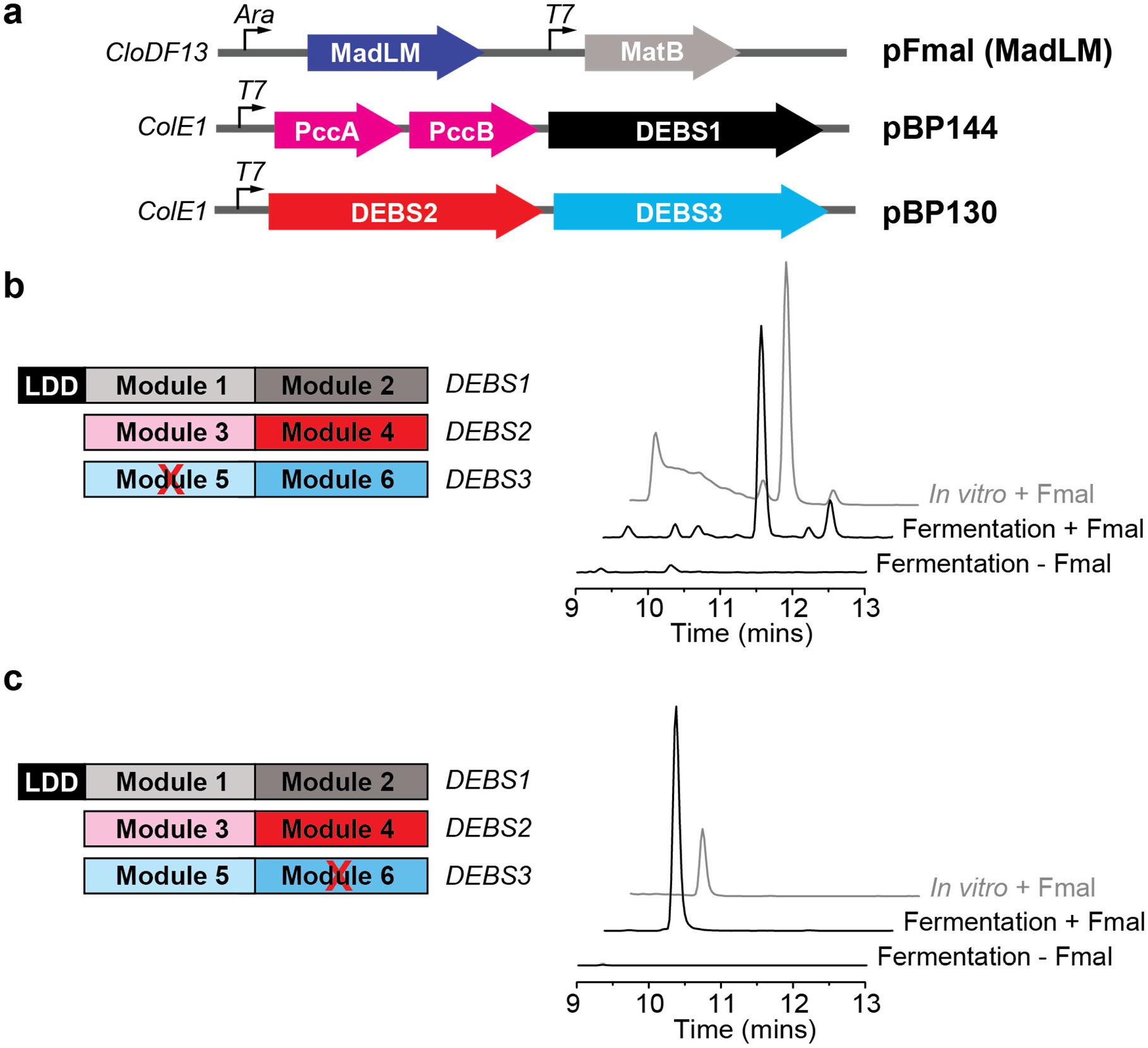
Regiospecific incorporation of fluorine into complex polyketides by *E. coli* production host. **(a)** Engineered pathway for production of fluorinated 6dEB analogs in *E. coli*. The system includes three plasmids with compatible origins and antibiotic markers, which together encode for six pathway proteins. pFmal(MadLM) plasmid (cloDF13 origin, Sp^R^) encodes for malonate transporter MadLM, and malonyl-CoA synthetase MatB. pBP130 (ColE1 origin, Cb^R^) and pBP144 (ColE1 origin, Km^R^) encode for DEBS1, DEBS2, and DEBS3 proteins. pBP144 also encodes for a propionyl-CoA carboxylase (PccAB). **(b)** Extracted ion chromatograms showing the generation of the 4-fluoro-4-desmethyl 6dEB analog (*m/z* 373.2→275.1) in culture medium of *E. coli* expressing Mod5 AT^0^ DEBS. Chromatograms shown are representative of at least three biological replicates. **(c)** Extracted ion chromatograms showing the generation of 2-fluoro-2-desmethyl 6dEB analog (*m/z* 373.2→275.1) in culture medium of *E. coli* expressing Mod6 AT^0^ DEBS. Chromatograms shown are representative of at least three biological replicates.

In summary, we have developed a method to introduce fluorine site-selectively into polyketide natural products that are immediate drug precursors through engineered biosynthesis in living cells. Applications of this platform open opportunities for discovery of new bioactive compounds by expanding the accessible structural space for organofluorines.

## Methods

### Commercial materials

Commercial materials used and their sources are provided in Supplementary Information.

### Bacterial strains

*E. coli* DH10B-T1^R^ was used for DNA construction. *E. coli* BL21(DE3)-T1^R^ , BAP1^41^ , and BAP1-T1^R 42^ were used for heterologous protein expression. BAP1 and BAP1-T1^R^ were used for expression of DEBS modules and DEBS ACP domains as they require modification with phosphopantethiene. BL21(DE3)-T1^R^ was used for all other proteins.

### Lysate assay for screening DszAT mutant library

DszAT lysate is prepared by inoculating 50 mL of TB (containing 50 µg/mL Carbenicillin) with freshly transformed *E. coli* harboring pFW3 and its derivatives (OD_600_ = 0.05). The cultures were grown to OD_600_ of 0.6-0.8 at 37°C, cold-shocked on ice for 10-20 min, subsequently induced with 400 µM IPTG, and grown overnight at 16°C. 10 mL aliquots of each culture were pelleted by centrifugation at 12,000 × g for 5 minutes at 4°C. The cell pellets were then resuspended in 290 mM sodium phosphate pH 7.5 at 1 mg/mL concentration. The cells were lysed with 0.1 mm bacterial glass disruption beads using Mini-BeadBeater-24 (Biospec products, Bartlesville, OK) in 2 × 45 s exposures at maximum speed. The resulting homogenate was spun down at 20,000 × g at 4°C for 20 minutes and the supernatant was used as the source of DszAT variants in the assay. Lysate from a strain containing empty pET51b+ vector (Invitrogen, Waltham MA) was analyzed as negative control for background endogenous transacylation by *E. coli* BL21(DE3)-T1^R^ .

The assay mixture contained equal volume of cell lysate and regeneration system (100 µL total). Final assay mixtures contained 400 mM sodium phosphate, pH 7.5, phosphoenolpyruvate (50 mM), TCEP (5 mM), magnesium chloride (10 mM), ATP (2.5 mM), pyruvate kinase/lactate dehydrogenase (15 U/mL), adenylate kinase (10 U/mL), methylmalonyl-CoA epimerase (5 µM), CoA (1 mM), MatB (20 µM), methyl- or fluoromalonate (5 mM). The mixture was incubated at 37ºC for 30-45 min and initiated by the addition of NDK-SNAC (5 mM) and DEBS protein (10 µM). The reaction was incubated for 24 h at 37°C. 50 µL aliquots were removed, quenched with 2.5 µL of 70% (*v/v*) perchloric acid, and centrifuged at 18,000 × g for 10 min to pellet the precipitated protein. The supernatant was removed and flash frozen.

Before analysis, frozen samples were centrifuged at 18,000 × g for 5 min at to remove salts. The supernatant was removed and analyzed by LC-QQQ using MRM in negative ionization mode on an EclipsePlus C-18 RRHD column (1.8 µm, 2.1 × 50 mm, r.t, Agilent) or a Poroshell 120 SB-Aq column (2.7 µm, 2.1 × 50 mm, r.t, Agilent) using a linear gradient from 0 to 40% acetonitrile over 4 min with 0.1% formic acid as the aqueous mobile phase after an initial hold at 0% acetonitrile for 12 s at 0.6 mL/min. The transitions are as follows (parent m/z→ product m/z, fragmentation voltage, collision energy): H-TKL: 155 → 97, 100, 5; F-TKL: 173 → 59, 135, 20; H-TTKL: 197 → 95, 135, 5; TKL: 169 → 111, 135, 10. Production of TKLs and tetraketide pyrones (TTKPs) were normalized to the corresponding amounts observed with WT DszAT samples analyzed in conjunction.

### *In vitro* assay for screening selected DszAT mutants

All assay mixtures (30 µL) contained 5 µM methylmalonyl-CoA epimerase (Epi), 20 µM MatB, 2.5 mM ATP, 15 U/mL PK, 10 U/mL adenylate kinase, and 50 mM phosphoenol pyruvate (PEP) in 400 mM sodium phosphate, 10 mM magnesium chloride, 5 mM TCEP, pH 7.5. Reactions contained 1 mM carboxyacyl-CoA (fluoromalonyl-CoA, malonyl-CoA, or methylmalonyl-CoA), 2.5 mM NDK-SNAC, 10 µM Mod3_DEBS_+TE(AT^0^), and 10 µM DszAT variants. Epi, MatB, PK, myokinase, PEP, ATP, and carboxyacyl-CoA substrate were pre-incubated at 37 °C for 30 min. Reactions were then initiated with NDK-SNAC, Mod3_DEBS_+TE(AT^0^), and *trans*-AT, and incubated for 16 h at 37°C. 50 µL aliquots were removed, quenched with 2.5 µL of 70% (*v/v*) perchloric acid, and centrifuged at 18,000 × g for 10 min. The supernatant was removed and stored at -80 °C until analysis. Reactions were analyzed as in lysate assay.

### Steady-state kinetic characterization of hydrolysis activity

The DszAT assay was adapted from a coenzyme A (CoA) release assay previously described in the literature.^43-45^ Assays were performed in 96-well microplate (black, polystyrene, µ-clear, f-bottom, chimney well, med binding; Greiner Bio-One) by monitoring free CoA with DTNB at 412 nm using a Synergy Mx microplate reader (Biotek). Reactions were monitored for 4 min using the minimum interval setting between measurements.

Assay components were prepared in three solutions in 50 mM sodium phosphate, 1 mM EDTA, 10% (*v/v*) glycerol, pH 7.5: Solution 1, 2.4 mM DTNB; Solution 2, 4× fluoromalonyl-CoA; Solution 3, 20 µM *trans*-AT enzyme and 0.1 mg/mL BSA. Solution 1 (25 µL) and Solution 2 (25 µL) were added to the wells and pre-incubated at room temperature for 15-20 min. The reactions were then initiated with Solution 3 (50 µL). The final concentrations of fluoromalonyl-CoA were 200, 150, 100, 50, 25, 12.5, 6.25, 3.13, 1.56, 0.78, 0.39, and 0.20 μM. Absorbance values were converted to concentrations using the ε_412 nm_ = 14,150 M^-1^ cm^-1^ and a pathlength of 0.3 cm. Initial velocities (*v*_0_) were defined as the linear portion of the change of product concentration over time curve. The data were fit by non-linear curve fitting to the standard Michaelis-Menten equation.

### Steady-state kinetic characterization of transacylation activity

The transacylase assay was adapted from a previously described method.^27, 46, 47^ Assays were performed in 96-well microplate (half area, µCLEAR, black polystyrene, medium binding, non-sterile, Greiner Bio-One) with NADH fluorescence (λ_Ex_ = 340 nm, λ_Em_ = 450 nm) monitored using a Synergy Mx microplate reader (Biotek; top optical position at 8 mm read height, gain = 100). Reactions were monitored for 5 mins at 30 °C using minimum interval setting between measurements.

Assay components were prepared in three solutions in 50 mM sodium phosphate, 1 mM EDTA, 10% (*v/v*) glycerol, pH 7.5: Solution 1, 2.4 mU/µL α-ketoglutarate dehydrogenase, 1.6 mM NAD^+^ , 1.6 mM TPP, and 8 mM α-ketoglutarate; Solution 2, 4× carboxyacyl-CoA (malonyl-CoA, methylmalonyl-CoA, or fluoromalonyl-CoA); Solution 3, 150 µM ACP_DEBSMod6_, 2× *trans*-AT , and 0.1 mg/mL BSA. Solution 1 (12.5 µL) and Solution 2 (12.5 µL) were added to the wells and pre-incubated at room temperature for 15-20 min. The reactions were then initiated with Solution 3 (25 µL).

The final concentrations of malonyl-CoA, methylmalonyl-CoA, and fluoromalonyl-CoA were 200, 150, 100, 50, 25, 12.5, 6.25, and 3.13 μM. The final concentrations of DszAT (WT and F190V) were 50 nM for malonyl-CoA and fluoromalonyl-CoA and 10 µM for methylmalonyl-CoA. The final concentrations of FabD were 30 nM for malonyl-CoA, 50 nM for fluoromalonyl-CoA, and 100 nM for methylmalonyl-CoA reactions. Controls with no *trans*-AT added were run in parallel for all reactions. Fluorescence was converted to concentration using an NADH standard curve collected externally in the same assay matrix with carboxyacyl-CoA substrate, ACP domain, and DszAT omitted. Initial velocities (*v*_0_) were defined as the linear portion of the change of product concentration over time curve. Transacylation with methylmalonyl-CoA was fit by non-linear curve fitting to the standard Michaelis-Menten equation. Transacylation with malonyl-CoA and fluoromalonyl-CoA were fit by non-linear curve fitting to the Hill equation with substrate inhibition.

### Assay for in vitro triketide lactone production

All assays contained 5 µM methylmalonyl-CoA epimerase (Epi), 20 µM MatB, 2.5 mM ATP, 15 U/mL PK, 10 U/mL myokinase, and 50 mM phosphoenol pyruvate in 400 mM sodium phosphate, 10 mM magnesium chloride, 5 mM TCEP, pH 7.5.

To compare WT and F190V DszAT under pure and mixed carboxyacyl-CoA extender unit conditions, the assay mixture contained 1 mM fluoromalonyl-CoA, malonyl-CoA, and/or methylmalonyl-CoA, 10 mM NDK-SNAC, 10 µM Mod3_DEBS_+TE(AT^0^) and DszAT variants. Epi, MatB, PK, myokinase, PEP, ATP, and carboxyacyl-CoA substrate were pre-incubated at 37 °C for 30 min. Reactions were then initiated with NDK-SNAC, Mod3_DEBS_+TE(AT^0^), and *trans*-AT and incubated for 1 h at 37 °C.

For FabD activity assay, assay mixture contained 1 mM of malonate substrate (fluoromalonate, malonate, or methylmalonate), 1 mM of coenzyme A, 1 mM NDK-SNAC, 10 µM Mod3_DEBS_+TE(AT^0^), and 30 µM of *trans*-AT enzyme. Epi, MatB, PK, myokinase, PEP, ATP, and acyl-CoA substrate were pre-incubated at 37 °C for 30 min. The reactions were then initiated with NDK-SNAC, Mod3_DEBS_+TE(AT^0^), and *trans*-AT and incubated overnight at 37 °C.

To quench the reaction, 25 µL of reaction mixture was mixed with 25 µL of 10% (*v/v*) perchloric acid and centrifuged for 10 min at 4 °C at 20,817 × *g*. The supernatant was removed and stored at -80 °C until analysis.

### Quantification of triketide lactone products by LC-QQQ

Frozen samples were centrifuged for 10 min at 4 °C at 20,817 × *g* to remove salts. The supernatant was removed and analyzed by LC-QQQ using MRM in positive ionization mode on an Agilent 6460 QQQ MS. Samples were analyzed on two tandem Ascentis Express RP-amide HPLC columns (2.7 µm, 2.1 mm × 10 cm, 25 °C; Supelco) on an Agilent 1290 UPLC using an isocratic solvent system of 10% acetonitrile in 0.1% (*v/v*) formic acid as the aqueous mobile phase for 20 min after an initial ramp of 0 to 10% acetonitrile over 0.5 min at flow rate of 0.4 mL/min. Products were monitored with transitions as follows (parent ion *m/z* → product ion *m/z*, fragmentor, collision energy): TKL (171 → 153, 90,6); H-TKL (157 → 139, 90, 4); F-TKL (175 → 157, 90, 5); H-TTKP (199 → 111, 100, 9). The identity and the quantity of each compound was verified against synthetic standards using external standard curves. Synthetic standards were obtained as previously described.^26^

### Assay for *in vitro* 6dEB analog production

All assays (Mod6 AT^0^ DEBS: 30 µL, Mod3 AT^0^ DEBS and Mod5 AT^0^ DEBS: 50 μL) contained 2 µM LDD_DEBS_, 2 µM Mod1_DEBS_, 2 µM Mod2_DEBS_, 2 µM DEBS2 variant (WT or DEBS2(Mod3 AT^0^)), and 2 µM DEBS3 variant (WT, DEBS3(Mod5 AT^0^), or DEBS3(Mod6 AT^0^)), 2 µM MatB, 4 µM Epi, 2 µM PrpE, 1 mM CoA, 4 mM ATP, 2 mM methylmalonate, 0.5 mM NADPH, 0.5 mM propionate in 250 mM sodium phosphate, 10 mM magnesium chloride, 5 mM TCEP, pH 7.5. When used, 10 mM fluoromalonate, 2 mM malonate, and 6 µM DszAT variant (WT or F190V) were added. Reaction mixtures were incubated at room temperature for 3 h before quenching and extracting twice with 4 × reaction volume of ethyl acetate The combined organic layers were dried using a Speed-Vac SC110 (Savant).

### Identification of 6dEB analogs by LC-QTOF and LC-QQQ

The reaction residue was resuspended in methanol (Mod6 AT^0^ DEBS, 30 µL; Mod3 AT^0^ DEBS and Mod5 AT^0^ DEBS, 25 μL). and centrifuged for 10 min at 4 °C at 20,817 × *g*. Products were analyzed by LC-QTOF in positive ionization mode on at Agilent 6530B QTOF MS. Samples were analyzed on Ascentis Express RP-amide HPLC columns (2.7 µm, 2.1 mm × 10 cm, 25 °C; Supelco) on an Agilent 1290 UPLC using a linear gradient from 5 to 65% acetonitrile over 16.75 min with 0.1% (*v/v*) formic acid as the aqueous mobile phase after an initial increase of 0 to 5% acetonitrile over 30 s at flow rate of 0.25 mL/min. Masses observed correspond to [M+H^+^ -H_2_O]^+^ of the target molecules as follows (molecular formula, expected mass, observed mass, Δppm(*exp*-*obs*)): 6dEB (C_21_H_37_O_5_^+^ , 369.2636, 369.2639, 0.08 ppm), desmethyl 6dEB (C_20_H_35_O_5_^+^ , 355.2479, 355.2476, 0.8 ppm), and fluorodesmethyl 6dEB (C_20_H_34_FO_5_^+^ , 373.2385, 373.2384, 0.3 ppm). Fragmentation analysis was performed using the targeted MS/MS mode with fragmentation voltage of 185 and collision energy of 10. From the fragmentation patterns, unique transitions were developed for detection with increased sensitivity with LC-QQQ using MRM in positive ionization mode on an Agilent 6460 QQQ MS with the same LC method. Products were monitored with transitions as follows (parent *m/z* → product *m/z*, fragmentor, collision energy): 6dEB (369.2→239.1, 50, 1), 2-desmethyl 6dEB and 4-desmethyl 6dEB (355→239.1, 50, 1), 2-fluoro-2-desmethyl 6dEB, and 4-fluoro-4-desmethyl 6dEB (373.2→275.1, 105, 0).

### *In vivo* production of triketide lactones in *E. coli*

LB (50 mL) containing appropriate antibiotics (carbenicillin and chloramphenicol, 50 μg/mL; spectinomycin, 100 μg/mL) in a 250 mL baffled flask was inoculated with 1 mL of an overnight LB culture of *E. coli* BAP1 or BAP1-T1^R^ transformed with relevant plasmids. Cultures were grown at 37 °C with shaking at 200 rpm to OD_600_ = 0.4-0.6. Then, cultures were cooled on ice for 20 min and protein expression was induced with 1 mM IPTG and 0.2% (*w/v*) arabinose. Induced cultures were incubated at 16 °C with shaking for 16-24 h. Cells (40 mL) were harvested by centrifugation at 1000 × g for 15 min at 4 °C after recording the final OD_600_. Cells were washed once with 40 mL of M9 medium and resuspended in M9 medium to OD_600_ ∼ 100. The high-density cell suspension (50 µL) was added to a 1.7 mL microcentrifuge tube along with 5 mM fluoromalonate (1 M stock in water) and 1 mM NDK-SNAC (250 mM stock in 25% DMSO). Cell suspensions were incubated with the two substrates at 16 °C with shaking at 200 rpm for 20-24 h. Then, the cell suspension was centrifuged at 20,817 × g for 1 min at 4 °C. The supernatant (35 µL) was removed and mixed with 10% (*v/v*) perchloric acid (35 µL) and centrifuged at 20,817 × g for 10 min at 4 °C. The supernatant was removed and stored at -80 °C until analysis.

### *In vivo* production of 6dEB analogs in *E. coli*

6dEB analogs were produced in *E. coli* BAP1 or BAP1-T1^R^ cultures grown in growth medium previously optimized for 6dEB production ^48^ . The 6dEB production medium (50 mL) containing appropriate antibiotics in a 250 mL baffled flask was inoculated with 1 mL of overnight culture of *E. coli* transformed with relevant plasmids in the same medium. Cultures were grown to OD_600_ = 0.4-0.6 at 37 °C with shaking at 200 rpm. Then, cultures were cooled on ice for 20-40 min and protein expression was induced with 1 mM IPTG. Cultures were incubated at 22 °C with shaking at 250 rpm for 16-24 h. Cells were then collected by centrifugation at 10,000 × *g* for 5 min at 4 °C and resuspended in 1.5 mL of fresh media containing 1mM IPTG, 0.2% (*w*/*v*) arabinose, 5 mM fluoromalonate, 20 mM propionate, and appropriate antibiotics. High-density cell suspensions were grown in 14-mL round-bottom Falcon culture tubes for another 16-24 h at 22 °C with shaking at 250 rpm, after which they were collected by centrifugation at 20,817 × *g* for 1 min 4°C. Supernatant (1.5 mL) was removed and extracted with 4 × 0.5 mL ethyl acetate by vortexing for 15-30 s and separating the layers by centrifugation at 13,000 × *g* for 3 min. The combined organic layers were dried by Speed Vac SC110 (Savant), resuspended in 30 µL of methanol for analysis.

### Large scale production of 2-fluoro-2-desmethyl 6dEB analog

6dEB production medium (12 × 500 mL) containing appropriate antibiotics in a 12 × 2.5 L Ultrayield flask was inoculated with 5 mL of overnight culture of *E. coli* harboring pBP130_DEBS3(Mod6 AT^0^), pBP144, and pFmal(MadLM) in the same medium. Cultures were grown to OD_600_ = 0.4-0.6 at 37 °C with shaking at 200 rpm. Cells were collected by centrifugation at 8,000 × *g* for 5 min at 4 °C and resuspended in 20 mL of fresh media containing 1 mM IPTG, 0.2% (*w*/*v*) arabinose, 5 mM fluoromalonate, 20 mM propionate, and appropriate antibiotics. The high-density cell suspension was transferred to 250 mL baffled flask and incubated for another 16-24 h at 22 °C with shaking at 250 rpm. Supernatant was collected for extraction.

### Purification of 2-fluoro-2-desmethyl 6dEB analog from *E. coli* culture medium

Supernatant (40 mL) was extracted with 4 × 10 mL ethyl acetate by vortexing for 15-30 s and separating the layers by centrifugation at 8,000 × *g* for 5 min. The combined organic layers were dried by rotary evaporation. The extract was purified on an Eclipse XDB-C18 (5 µm, 9.4 mm × 250 mm, Agilent) on an Agilent 1200 HPLC using a linear gradient from 15 to 45% acetonitrile over 42 min with water as the aqueous mobile phase after an initial increase of 0 to 15% acetonitrile over 1.5 min at flow rate of 4 mL/min. Fractions (2 mL) were collected and screened using LC-QQQ. Fractions were analyzed on Poroshell 120 EC-C18 (2.7 µm, 2.1 mm × 50 mm, Agilent) on Agilent 1290 UPLC using a linear gradient from 25 to 60% acetonitrile over 2 min with 0.1% (*v/v*) formic acid as the aqueous mobile phase after an initial increase of 0 to 25% acetonitrile over 12 s at flow rate of 0.7 mL/min. Fractions containing 2-fluoro-2-desmethyl 6dEB shown by transition 373.2→275.1 were combined and lyophilized to dryness.

### ^19^F-NMR of fluorinated 6dEB analog isolated from *E. coli* culture medium

Lyophilized culture extract containing 2-fluoro-2-desmethyl 6dEB was resuspended in 500 µL of 50:50 MeOH:D_2_O mixture for ^19^F-NMR analysis. 5-fluorouracil (1 μM, 100 μL) in D_2_O was added to the coaxial insert as internal standard. ^19^F-NMR spectrum was collected at 298 K on a Bruker AV-600 spectrometer at the College of Chemistry NMR Facility. Tune and shim were optimized. Spectrum was collected using the 19f.cocnmr.av600 pulse program with following parameters: o1p = -200, sw = 200, td =131072, d1 = 1 s, ds = 20, ns = 3200. Resulting spectrum was processed by MestReNova by backward linear prediction from 0 to 128 using Toeplitz method and phased manually. Spectrum was referenced by a set of doublet corresponding to 5-fluorouracil at -171 ppm.^19^F-NMR (600 MHz, MeOH:D_2_O) δ -195.6 (dd, *J*_1_ = 54 Hz, *J*_2_ = 12 Hz).

## Notes

### Competing Interest Statement

The authors have declared no competing interest.

