## Supplemental Information File for "Engineering site-selective incorporation of fluorine into natural product analogs"

Sasilada Sirirungruang, Omer Ad, Thomas M. Privalsky, Swetha Ramesh, Joel L. Sax, Hongjun Dong, Edward E. K. Baidoo, Bashar Amer, Chaitan Khosla, Michelle C. Y. Chang*

#### Supplementary Methods

*Commercial materials* S2
*Gene and plasmid construction* S2
*Expression of His-tagged proteins* S6
*Purification of His-tagged proteins* S7
*Synthesis of N-acetylcysteamine thioester of (2S,3R)-2-methyl-3-hydroxypentanic acid (NDK-SNAC)* S9
*Synthesis of fluoromalonate* S10
*Synthesis of fluoromalonyl-CoA* S10

#### Extended Data Figures

*Extended Data Fig. 1. Engineering a fluoromalonyl-CoA specific trans-AT* S11
*Extended Data Fig. 2. Triketide lactone formation assay* S12
*Extended Data Fig. 3. trans-AT library screening* S13
*Extended Data Fig. 4. Characterization of DszAT F190V* S15
*Extended Data Fig. 5. SDS-PAGE of purified proteins used for enzymatic generation of fluorodesmethyl 6dEB analogs.* S17
*Extended Data Fig. 6. Generation of 2-fluoro-2-desmethyl-6dEB analog through in vitro reconstitution of Mod6 $AT^0$ DEBS* S18
*Extended Data Fig. 7. Fragmentation reactions observed of 6dEB and analogs* S20
*Extended Data Fig. 8 Characterization of 2-desmethyl-6dEB analog through in vitro reconstitution of Mod6 $AT^0$ DEBS.* S21
*Extended Data Fig. 9. Generation of 4-fluoro-4-desmethyl-6dEB analog through in vitro reconstitution of Mod5 $AT^0$ DEBS* S22
*Extended Data Fig. 10. Characterization of E. coli FabD and its role in polyketide biosynthesis* S25
*Extended Data Fig. 11. Influence of extender unit availability on in vivo selectivity of chain elongation by single-modular DEBS constructs in engineered E. coli* S28
*Extended Data Fig. 12. In vivo production of desmethyl 6dEB analogs by engineered E. coli* S29
*Extended Data Fig. 13. In vivo production of monofluorinated desmethyl 6dEB analogs by engineered E. coli* S31

#### Supplementary Tables

*Table S1. Strains, plasmids, oligonucleotides, and sequences.* S33

#### References

S38

#### Supplementary Methods

**Commercial materials.** Luria-Bertani (LB) Broth Miller, LB Agar Miller, Terrific Broth (TB), magnesium sulfate anhydrous, and glycerol were purchased from EMD Biosciences (Damstadt, Germany). Carbenicillin (Cb), glucose, isopropyl-β-D-thiogalactopyranoside (IPTG), sodium chloride, calcium chloride, 4-(2-hydroxyethyl)-1-piperazineethanesulfonic acid (HEPES), magnesium chloride hexahydrate, kanamycin (Km), acetonitrile, dichloromethane, ethyl acetate, ethylene diamine tetraacetic acid disodium dihydrate (EDTA), sodium phosphate monobasic, potassium phosphate monobasic, ammonium chloride, and arabinose were purchased from Fisher Scientific (Pittsburgh, PA). Coenzyme A trilithium sodium salt (CoA), malonyl-CoA, methylmalonyl-CoA, malonic acid, methylmalonic acid, tris(2-carboxyethyl)phosphine (TCEP) hydrochloride, phosphoenolpyruvate (PEP), adenosine trisodium phosphate sodium salt (ATP), adenylate kinase (myokinase), pyruvate kinase, lactate dehydrogenase, poly(ethyleneimine) (PEI), β-mercaptoethanol (BME), thiamine pyrophosphate (TPP), α-ketoglutaric acid, sodium phosphate dibasic heptahydrate, cysteamine, 4-hydroxy-6-methyl-2-pyrone acetic anhydride, 1-ethyl-3-(-dimethylaminopropyl)carbodiimide hydrochloride (EDC), 4-dimethylaminopyridine (DMAP), reduced β-nicotinamide adenine dinucleotide 2′-phosphate (NADPH), acetonitrile, dimethyl sulfoxide (DMSO), biotin, thiamine hydrochloride, sodium propionate, ammonium formate, chloramphenicol, β-nicotinamide adenine dinucleotide (NAD+), reduced β-nicotinamide adenine dinucleotide (NADH), β-nicotinamide adenine dinucleotide 2′-phosphate (NADP+), α-ketoglutarate dehydrogenase, and bovine serum albumin were purchased from Sigma-Aldrich (St. Louis, MO). Diethylfluoromalonate was purchased from Sigma-Aldrich (St. Louise, MO) or from Matrix Scientific (West Columbia, SC). Spectinomycin (Sp) was purchased from Chem-Impex International (Wood Dale, IL). Formic acid, dithiothreitol (DTT), and perchloric acid were purchased from Fluka (Charlotte, NC). 2-methyl-3-oxopentanoic acid ethyl ester was purchased from Fragmenta (Monmouth Junction, NJ). Mini-PROTEAN TGX 8-16%, and Bio-Rad Protein Assay Dye Reagent concentrate was purchased from Bio-Rad Laboratories (Hercules, CA). Restriction enzymes, T4 DNA ligase, and Phusion DNA polymerase were purchased from New England Biolabs (Ipswich, MA). GoTaq Green Master Mix was purchased from Promega (Madison, WI). Deoxynucleotides (dNTPs) were purchased from Invitrogen (Carlsbad, CA). PageRuler™ Plus prestained protein ladder was purchased from Fermentas (Glen Burnie, Maryland). Oligonucleotides were purchased from Integrated DNA Technologies (Coralville, IA), resuspended at a stock concentration of 100 μM in water. DNA purification kits and Ni-NTA agarose were purchased from Qiagen (Valencia, CA). PD-10 columns Sephadex G-25M was purchased from GE (Chicago, IL). Complete EDTA-free protease inhibitor was purchased from Roche Applied Science (Penzberg, Germany). Zymoclean Large Fragment DNA Recovery Kit was purchased from Zymoresearch (Tustin, CA). Amico Ultra 3,000 MWCO, 10,000 MWCO, 30,000 MWCO, and 100,000 MWCO centrifugal concentrators were purchased from EMD Millipore (Billerica, MA). Chloroform-d and $D_2O$ were purchased from Cambridge Isotope Laboratories (Andover, MA). $^{19}$F-NMR spectra were collected at 25 ºC on Bruker AV-600 at the College of Chemistry NMR Facility at the University of California, Berkeley. High-resolution mass spectral analyses were carried out on an Agilent 6530 Quadrupole-Time-of-Flight (QTOF) Accurate Mass spectrometer and an Agilent 6460 Triple Quadrupole (QQQ).

**Gene and plasmid construction.** Standard molecular biology techniques were used to carry out plasmid construction. All PCR amplifications were carried out with Phusion High Fidelity DNA polymerase or with Taq DNA polymerase using GoTaqGreen master mix. For amplification of GC-rich sequences, PCR reactions carried out with Phusion High Fidelity DNA polymerase were performed in the GC buffer supplemented with 10 % (*v/v*) DMSO and 1 M betaine with primer annealing temperatures 4-8 °C below the $T_m$. DNA assembly was performed using the isothermal Gibson assembly protocol [*1*]. For analysis and isolation of large DNA fragments from agarose gel, 0.3%-0.6 % gel and Zymoclean large fragment DNA recovery kit (Zymoresearch) were used. All constructs were verified by sequencing (UC Berkeley DNA Sequencing Facility, Berkeley, CA; and MGH CCIB DNA Core, Center for Computational and Integrative Biology, Massachusetts General Hospital, Cambridge, MA). pFW3 [*2*], pBL12 , pBL13, pBL36, pFW98, and pFW100 [*3*] were gifted by the laboratory of Professor Chaitan Khosla (Stanford University). pBP130 and pBP144 [*4*] were gifts from the laboratory of Professor Blaine Pfeifer at the University at Buffalo.

*DszAT mutant libraries.* DszAT mutant libraries were generated by site-specific mutagenesis at selected catalytic residues within the active site of the enzyme. The libraries constructed full saturation at the amino acids at the F190 position (according to DszD numbering), a library at the S86 catalytic residue (S86C, S86D, S86E, S86A) and a double mutant S86; H191 library for the same mutants (S86C; H191A, S86D; H191A, and S86E; H191A) was also constructed. Finally, a double mutant library for the F190; L87 residues was generated for select F190 mutants from the single mutant screens a secondary screen for enhanced fluorine substrate selectivity (F190G; L87A, F190G; L87V, F190S; L87A, F190S; L87V, F190T; L87A, F190T; L87V, F190V; L87A, F190V; L87V, F190P; L87A, F190I; L87A, and F190I; L87V). The mutant library was constructed by amplification from pFW3.

The F190 mutations were introduced by amplifying pFW3 with two sets of primers. One fragment was amplified with DszAT F190 F1 and various F190 R1 primers (depending on mutation). The second fragment was amplified using various F190 F2 primers (depending on mutation) and DszAT F190 R2. The two fragments contained a 60 bp overlap. After treatment with DpnI, the two PCR fragments were inserted into the XbaI-HindIII sites of pFW3 using the Gibson Protocol. The L87 mutations were introduced by amplifying pFW3 with two sets of primers. One fragment was amplified with DszAT L87 F1 and various L87 R1 primers (depending on mutation). The second fragment was amplified using various L87 F2 primers (depending on mutation) and DszAT L87 R2. The two fragments contained a 60 bp overlap. After treatment with DpnI, the two PCR fragments were inserted into the XbaI-HindIII sites of pFW3 using Gibson assembly.

The H191A mutation was introduced by amplifying pFW3. One fragment was amplified with DszAT H191A F1/R1 and the second fragment was amplified DszAT H191A F2/R2. The two fragments contained a 60 bp overlap. After treatment with DpnI, the two PCR fragments were inserted into the NdeI-EcoRI sites of pFW3 using Gibson assembly.

The S86 mutations were introduced by amplifying pET21c-DszAT H191A-His6 or pFW3 with two sets of primers. One fragment was amplified with DszAT S86 F1 and various S86 R1 primers (depending on mutation). The second fragment was amplified using various S86 F2 primers (depending on mutation) and DszAT S86 R2. The two fragments contained a 60 bp overlap. After treatment with DpnI, the two PCR fragments were inserted into the XbaI-HindIII sits of pFW3 or pET21c-DszAT-H191A-His6 using Gibson assembly.

All double mutants were designed as described above, with the exception of PCR template containing desired mutations from the initial round of screening.

*pET16b-His$_{10}$Pres-ACP$_{DEBSMod6}$.* The ACP domain of Mod6$_{DEBS}$ was amplified from pAYC-138 [*2*] using primers resACP6_Fwd3 and PresACP6_Rev. The PCR product was assembled into the NdeI-BamHI site of a modified pET16b with the Factor Xa cleavage site replaced by the PreScission cleavage site via Gibson assembly.

*pFW3_F190V.* pFW3_F190V was constructed by amplification from pFW3 with two pairs of primers: DszAT F190 F1/DszAT F190V R1 and DszAT F190V F2/DszAT F190 R2. Primers DszAT F190V R1 and DszAT F190V F2 contain the F190V mutation to be introduced to the plasmid. The two PCR products were inserted into the XbaI-HindIII site of pFW3 via Gibson assembly.

*pFW98_DEBS2(Mod3 AT$^0$).* pFW98_DEBS2(Mod3 AT$^0$) encodes for DEBS2 with S653A mutation. pBP130_DEBS2(Mod3 AT$^0$) and pFW98 were digested with MauBI and Pfl23II. The 2 kB band from pBP130_DEBS2(Mod3 AT$^{0)}$ and the 14 kB band from pFW98 were gel-purified and ligated with T4 ligase. The resulting plasmids were screened via restriction mapping with KpnI. Plasmids with the correct restriction patterns were confirmed by complete plasmid sequencing at MGH CCIB DNA Core, Center for Computational and Integrative Biology, Massachusetts General Hospital (Cambridge, MA).

*pFW100_DEBS3(Mod5 AT$^0$).* pFW100_DEBS3(Mod5 AT$^0$) encodes for DEBS3 with S642A mutation. The plasmid was cloned by first replacing the section to be mutated from the parent plasmid with a Cm$^R$ marker and then removing it upon insertion of the piece bearing the desired mutation. This strategy allows for identification of mutants using antibiotic selection. Briefly, the Cm$^R$ cassette was first amplified from pACYC184 with the primers pACYC184_CmOperon_pFW100_M5_Fwd_PacI/pACYC184_CmOperon_pFW100_M5_Rev_SpeI and inserted into the SfiI-BsiWI site of pFW100 via Gibson assembly. *E. coli* cells transformed with the assembly mixture were selected on culture medium containing carbenicillin and chloramphenicol. Plasmids containing both resistance markers were further screened by restriction analysis with SacI. The pFW100 plasmid intermediates with the correct restriction pattern were confirmed by diagnostic sequencing. The target segments were amplified from pFW100 by two primer pairs to introduce the AT$^0$ mutation into module 5 of DEBS: pFW100_M5AT0_SfiI_F/pFW100_M5AT0_SfiI_R and pFW100_M5AT0_BsiWI_F/pFW100_M5AT0_BsiWI_R. The PCR products were then inserted via Gibson assembly into the pFW100 intermediate, which was digested with PacI and SpeI to remove the Cm$^R$ cassette. Resulting plasmids were selected for the loss of the Cm$^R$ marker and screened by SacI restriction analysis. Plasmids with the correct restriction patterns were confirmed by complete plasmid sequencing at MGH CCIB DNA Core, Center for Computational and Integrative Biology, Massachusetts General Hospital (Cambridge, MA).

*pFW100_DEBS3(Mod6 AT$^0$).* pFW100_DEBS3(Mod6 AT$^0$) encodes for DEBS3 with S2107A mutation. The plasmid was constructed using a similar strategy. The Cm$^R$ cassette was amplified from pACYC184 with primers pACYC184_CmOperon_pBP130_M6_Fwd_PacI/pACYC184_CmOperon_pBP130_M6_Rev_SpeI and inserted into the BbvCI-AjuI site of pFW100 via Gibson assembly. *E. coli* transformed with the assembly mixture were selected on culture medium containing carbenicllin and chloramphenicol. Plasmids containing both resistant genes were further screened by restriction analysis with SacI. The pFW100 plasmid intermediates with the correct restriction pattern were confirmed by diagnostic sequencing. Segments of module 6 of DEBS were amplified by two primers pairs to introduce the $AT^0$ mutation into $DEBS_{Mod6}$. The M6TE-SA-M6-RP/M6TE-SA-M6-FP primer pair was used to amplify a segment of module 6 containing the inactivating mutation to the AT domain from pAYC138, whereas the M6TE-SA-130-FP/M6TE-SA-130-RP primer pair was used to amplify another contiguous segment from pBP130 [*4*]. These PCR products were inserted via Gibson assembly into the pFW100 intermediate, which was digested with PacI and SpeI to remove the $Cm^R$ cassette. Resulting plasmids were selected for the loss of the $Cm^R$ marker and screened by SacI restriction analysis. Plasmids with the correct restriction patterns were confirmed by complete plasmid sequencing at MGH CCIB DNA Core, Center for Computational and Integrative Biology, Massachusetts General Hospital (Cambridge, MA).

*p15A-DszAT.* The gene encoding DszAT was amplified from pFW3 with the DszATF1/ T7TerminatorR1 primer pair. The lac operon was amplified from the same template with the LacCasetteLacI/LacCasetteLacO primer pair. The PCR products were inserted by Gibson assembly into the ClaI-PacI site of pJA4MCS2, which is pACYC184 [*2*] modified by replacing TcR cassette with a HindIII-XhoI-KpnI-XmaI-PacI-AflII-SFcI-PstI-AscI-AvrII-HincII linker.

*p15A-DszATS86A.* The S86A mutation was introduced to p15A-DszAT by amplifying p15A-DszAT with two sets of primers DszATF1/S86AMutationR and S86AMutationF/ DszATMutant_CTerm. S86AMutationR and S86AMutationF contain S86A mutation. The two PCR fragments were inserted by Gibson assembly into the NdeI-HindIII site of p15A-DszAT.

*p15A-DszATF190V.* F190V mutation was introduced to p15A-DszAT by amplifying p15A-DszAT with two sets of primers: DszATF1/F190VMutationR and F190VMutationF/ DszATMutant_CTerm. F190VMutationR and F190VMutationF contain S86A mutation. The two PCR fragments were inserted by Gibson assembly into the NdeI-HindIII site of p15A-DszAT.

*pET21c-FabD.* FabD gene was amplified from *E. coli* BL21(DE)-T1R using the MalACP_F1/ MalACP_R1 primer set. The PCR product was inserted into the NdeI-EcoRI site of pET21c by Gibson assembly.

*pBP144_DEBS1(Mod1 $AT^0$).* pBP144_DEBS1(Mod1 $AT^0$) encodes for DEBS1 with S1181A mutation. The plasmid was cloned by first replacing the section to be mutated from the parent plasmid with a CmR marker and then removing it upon insertion of the piece bearing the desired mutation. This strategy allows for identification of mutants using antibiotic selection. The $Cm^R$ cassette was amplified with primers Cm-pT7-PhaEC-F/Cm-pT7-PhaEC-R from pT7-CapPhaEC [*5*] and inserted into the BsiWI-BstBI site of BP144 via Gibson assembly. *E. coli* containing resulting plasmids were selected on culture medium containing carbenicillin and chloramphenicol. Plasmids containing both resistance markers were further screened by restriction analysis by XhoI. Plasmids with the correct restriction pattern were confirmed by diagnostic Sanger sequencing. Segments of DEBS Module 1 were then amplified from pBP144 by two primer sets, DEBS1-1/DEBS1-2, and DEBS1-3/DEBS1-4. DEBS1-2 and DEBS1-3 contained the desired S1181A mutation to be introduced to the active site of the AT domain of Module 1. The $Cm^R$-containing pBP144 intermediate was digested with BsiWI and BstBI to remove the $Cm^R$ insert, which was replaced with the mutated Module 1 segments by Gibson assembly to generate a scarless construct with the $AT^0$ mutation in Module 1. The resulting transformants were screened for Cm sensitivity followed by restriction analysis of the isolated plasmids by XhoI. Plasmids with the correct restriction patterns were confirmed by complete plasmid squencing to ensure that no other changes had been introduced.

*pBP130_DEBS2(Mod3 $AT^0$).* pBP130_DEBS2(Mod3 $AT^0$) encodes for DEBS2 with S653A mutation and DEBS3. The $Cm^R$ cassette was amplified with primers pACYC184_CmOperon_ pBP130_M3_Fwd_PacI/pACYC184_CmOperon_pBP130_M3_Rev_SpeI from pACYC184 and inserted into the MauBI-BaeI site of pBP130 using Gibson assembly. *E. coli* colonies containing resulting plasmids were selected on culture medium containing carbenicillin and chloramphenicol. A segment of DEBS Module 3 was amplified with primers pET21-M3-RP/pET21-M3-FP2 from pAYC136 to introduce the desired S653A into the AT domain of DEBS Module 3. The $Cm^R$-containing pBP130 intermediate was digested with PacI and SpeI to remove the $Cm^R$ cassette, which was replaced with the mutated Module 3 segments by Gibson assembly to generate a scarless construct with the $AT^0$ mutation in Module 3. The resulting transformants were screened for Cm sensitivity followed by restriction analysis of the isolated plasmids by XhoI. Plasmids with the correct restriction patterns were confirmed by complete plasmid sequencing to ensure that no other changes had been introduced.

*pBP130_DEBS3(Mod 5 $AT^0$).* pBP130_DEBS3(Mod5 $AT^0$) encodes for DEBS2 and DEBS3 with S642A mutation. The segment of DEBS Module 5 containing the desired S642A mutation was amplified with primers M5_BbvCI/ M5_NsiI from pFW100_DEBS3(Mod5 $AT^0$) and gel purified (3.5 kB). pBP130 was digested with BbvCI and NsiI and the resulting 22 kB band was gel purified to provide the backbone. The backbone and insert were then ligated with T4 ligase. The plasmid candidates were screened by restriction analysis with NotI. Plasmids with the correct restriction patterns were confirmed by complete plasmid sequencing to ensure that no other changes had been introduced.

*pBP130_DEBS3(Mod 6 $AT^0$).* pBP130_DEBS3(Mod6 $AT^0$) encodes for DEBS2 and DEBS3 with S642A mutation. pFW100_DEBS3(Mod6 $AT^0$) was digested with AsiSI and BbvCI and the resulting 6 kB fragment containing the desired S2107A mutation was gel purified. pBP130 was digested with AsiSI and BbvCI and the resulting 20 kB band was gel purified to provide the backbone. The backbone and insert were then ligated with T4 ligase. The plasmid candidates were screened by restriction analysis with XhoI. Plasmids with the correct restriction patterns were confirmed by complete plasmid sequencing to ensure that no other changes had been introduced.

**Expression of His-tagged proteins.** For $His_6$-MatB, DszAT-$His_6$, DszAT F190V-$His_6$, $His_{10}$-$ACP_{DEBSMod6}$, $Mod3_{DEBS}$+TE($AT^0$)-$His_6$, $Mod6_{DEBS}$+TE($AT^0$)-$His_6$, plasmids encoding the proteins of interest were transformed into *E. coli* BL21(DE3)-$T1^R$ ($His_6$-MatB, DszAT-$His_6$, DszAT F190V-$His_6$) or *E. coli* BAP1-$T1^R$ for proteins that require phosphopantetheine modification of ACP domains ($His_{10}$-$ACP_{DEBSMod6}$, $Mod3_{DEBS}$+TE($AT^0$)-$His_6$, $Mod6_{DEBS}$+TE($AT^0$)-$His_6$). TB culture medium (1 L) with appropriate antibiotic (carbenicillin, kanamycin, chloramphenicol: 50 µg/mL; spectinomycin: 100 µg/mL) in a 2.5 L ultra-yield flask was inoculated with overnight culture of freshly transformed *E. coli* cells. Cells were grown at 37 $^oC$ with shaking at 200 rpm to $OD_{600}$ = 0.6-0.8, at which time, they were cold-shocked on ice for 20-40 min. Expression was induced by addition of IPTG ($His_6$-MatB, $His_{10}$-$ACP_{DEBSMod6}$, DszAT-$His_6$, DszATF190V-$His_6$: 1 mM; $Mod3_{DEBS}$+TE($AT^0$)-$His_6$, $Mod6_{DEBS}$+TE($AT^0$)-$His_6$: 0.2 mM). Cells were then grown at 16 $^oC$ with shaking at 200 rpm overnight and harvested by centrifugation at 8,000 × *g* for 5 min at 4 ºC. Cell pellets were flash-frozen in liquid $N_2$ and stored at -80 ºC until purification.

For $His_6$-PrpE, $His_6$-Epi, $His_6$-$LDD_{DEBS}$, $Mod1_{DEBS}$-$His_6$, $Mod2_{DEBS}$-$His_6$, DEBS2-$His_6$, DEBS2(Mod3 $AT^0$)-$His_6$, DEBS3-$His_6$, DEBS3(Mod5 $AT^0$)-$His_6$, and DEBS3(Mod6 $AT^0$)-$His_6$, expression plasmids were introduced into *E. coli* BL21(DE3) ($His_6$-PrpE and $His_6$-Epi) or *E. coli* BAP1 cells to allow phosphopantetheinyl modification of ACP domains ($His_6$-$LDD_{DEBS}$, $Mod1_{DEBS}$-$His_6$, $Mod2_{DEBS}$-$His_6$, DEBS2-$His_6$, DEBS2(Mod3 $AT^0$)-$His_6$, DEBS3-$His_6$, DEBS3(Mod5 $AT^0$)-$His_6$, and DEBS3(Mod6 $AT^0$)-$His_6$). Overnight seed cultures were used to inoculate 0.5 L of LB medium with 2% glucose containing the appropriate antibiotic (carbenicillin, kanamycin: 50 µg/mL) in 2.5 L ultra-yield shake flask. Cells were grown at 37 ºC with shaking at 200 rpm to an approximate $OD_{600}$ of 0.3, at which point the temperature was slowly lowered. When cells reached $OD_{600}$ of 0.6, at which point cells were at approximately 20 ºC, protein expression was induced with IPTG ($His_6$-PrpE, $His_6$-Epi, $His_6$-$LDD_{DEBS}$, $Mod1_{DEBS}$-$His_6$, $Mod2_{DEBS}$-$His_6$, DEBS2: 0.1 mM; DEBS2-$His_6$, DEBS2(Mod3 $AT^0$)-$His_6$, DEBS3-$His_6$, DEBS3(Mod5 $AT^0$)-$His_6$, and DEBS3(Mod6 $AT^0$)-$His_6$: 0.5 mM). Cells were then grown at 18 ºC with shaking at 200 rpm overnight and harvested by centrifugation at 8,000 × *g* for 5 min at 4 ºC. Cell pellets were flash-frozen in liquid $N_2$ and stored at -80 ºC until purification.

**Purification of His-tagged proteins.** Cell pellets were resuspended in Lysis Buffer A (200 mM sodium phosphate, 200 mM sodium chloride, 30% (*v/v*) glycerol, 2.5 mM DTT, pH 7.5) at 5 mL/g of cell pellet. Lysozyme (0.5 mg/mL) and cOmplete EDTA-free protease inhibitor cocktail (Roche, 1 eq) were then added before homogenization and sonication. Cleared cell lysates were then obtained after centrifugation at 8,000 × *g* for 20 min at 4 ºC. To precipitate DNA, poly(ethyleneimine) was added dropwise to the soluble fraction at a final concentration of 0.015% (*w/v*). Precipitated DNA was removed by centrifugation at 8,000 × *g* for 20 min at 4 ºC. Protein concentration was determined using $A_{280}$ and protein extinction coefficients calculated from the sequence using the ExPASY ProtParam program.

*$His_6$-MatB, DszAT-$His_6$, DszAT F190V-$His_6$, and FadD-$His_6$* [*6*]. The lysate was diluted 3-fold with Wash Buffer B (50 mM sodium phosphate, 300 mM sodium chloride, 20% (*v/v*) glycerol, 20 mM BME, pH 7.5) with 10 mM imidazole and incubated with Ni-NTA agarose resin (2-4 mL) for 45-60 min at 4 ºC to bind the His-tagged protein before loading onto the column by gravity flow. The column was washed with Wash Buffer B containing 10 mM imidazole until the eluate was negative for protein content when tested by Bradford protein assay reagent (Bio-Rad). Column was further washed by wash buffer B containing 25mM imidazole until eluate was negative for protein content again. Protein was eluted from the column with Elution Buffer C (50 mM sodium phosphate, 50 mM sodium chloride, 20 mM BME, 20% (*v/v*) glycerol) containing 250 mM imidazole. Eluted protein was concentrated using a 10 kDa MWCO Amicon Ultra spin concentrator (Millipore) before exchanging into Storage Buffer D (50 mM HEPES, 2.5 mM EDTA, 100 mM sodium chloride, 2.5mM DTT, 20% (*v/v*) glycerol) using a PD-10 desalting column containing Sephadex G-25 resin (GE Healthcare). Purified protein was flash-frozen in liquid $N_2$ before storing at -80 ºC.

*$His_6$-$ACP_{DEBSMod6}$.* The protein was purified as described above up to protein elution. After elution from the Ni-NTA column, the concentration of the protein solution was determined using Bradford protein assay reagent (Bio-Rad) against BSA as a standard. PreScission protease was then added to the protein solution at a ratio of 1 mg PreScission:25 mg protein substrate. The proteolysis reaction was dialyzed 2× 1:200 overnight at 4 ºC into Storage Buffer D without DTT.

The mixture was then diluted 3-fold with Wash Buffer B without BME containing 10 mM imidazole. The solution was passed three times through Ni-NTA resin (2 mL) to remove PreScission protease and the cleaved $His_6$ tag before concentrating the eluate using a 3 kDa MWCO Amicon Ultra spin concentrator (Millipore). The purified protein was exchanged into Storage Buffer D without DTT using PD-10 desalting column with Sephadex G-25 resin (GE Healthcare). Thiol-containing reducing agents such as BME and DTT were excluded from buffers starting at the dialysis step as they interfere with transacylation assay. Purified protein was flash-frozen in liquid $N_2$ before storing at -80 ºC. Presence of the phosphopantethiene modification was checked by LC-TOF using positive ionization mode on an Agilent 6224 TOF MS. Samples were buffer exchanged into 10 mM sodium phosphate by 0.5 mL 7K Zeba Spin desalting column and analyzed on Proswift RP-4H HPLC column (1mm × 50 mm, room temperature) using a linear gradient from 10 to 100% acetonitrile over 3.5 min with 0.1% (*v/v*) formic acid as the aqueous mobile phase at flow rate of 0.3 mL/min. Resulting mass chromatograms were deconvoluted with Agilent MassHunter BioConfirm software.

*$Mod3_{DEBS}$+TE($AT^0$)-$His_6$, $Mod6_{DEBS}$+TE($AT^0$)-$His_6$* [6]. The cleared lysate was incubated with Ni-NTA resin (2-4 mL) for 45-60 min at 4 ºC to bind His-tagged protein before loading onto the column by gravity flow. The column was washed with Wash Buffer B containing 10 mM imidazole until eluate was negative for protein content when tested by Bradford protein assay reagent (Bio-Rad). Protein was then eluted from the column with Elution Buffer C containing 100 mM imidazole until eluate was negative for protein content. Eluted protein was concentrated using a 30 kDa MWCO Amicon Ultra spin concentrator and diluted 3-fold in Anion Exchange Buffer E (50 mM HEPES, 2.5 mM EDTA, 2.5 mM DTT, 20% (*v/v*) glycerol, pH 7.5) without sodium chloride. The diluted protein solution was loaded onto a 5 mL Hi-Trap Q HP column (GE Healthcare) using an NGC Medium-Pressure Liquid Chromatography System (Bio-Rad). The protein was eluted with a linear gradient from 0 to 1 M sodium chloride in Anion Exchange Buffer E over 30 col vol (150 mL). Fractions containing the target protein eluted ~350 mM sodium chloride were pooled and concentrated with a 30 kD MWCO Amicon Ultra spin concentrator. Purified protein was flash-frozen in liquid $N_2$ before storing at -80 ºC.

*DEBS2-$His_6$, DEBS2(Mod3 $AT^0$)-$His_6$, DEBS3-$His_6$, DEBS3(Mod5 $AT^0$)-$His_6$, and DEBS3(Mod6 $AT^0$)-$His_6$* [*3*]. Cell pellets were resuspended in Lysis Buffer F (50 mM sodium phosphate, 10 mM imidazole, 450 mM NaCl, 20% (*v/v*) glycerol, pH 7.6) at 5 mL/g of cell pellet. Lysozyme (0.5 mg/mL) and cOmplete EDTA-free protease inhibitor cocktail (Roche, 1 eq) were then added and incubated at 30 ºC for 30 min before sonication. Cleared cell lysates were obtained after centrifugation at 8,000 × *g* for 20 min at 4 ºC and loaded onto column containing Ni-NTA (10 mL/ 3 L cell culture) by gravity flow. The column was washed with Lysis Buffer F until eluate was negative for protein content when tested by Bradford protein assay reagent (Bio-Rad) The column was further washed with Wash Buffer G (50 mM sodium phosphate, 25 mM imidazole, 300 mM sodium chloride, 10% (*v/v*) glycerol, pH 7.5) before protein was eluted with Elution Buffer H (75 mM sodium phosphate, 500 mM imidazole, 20 mM sodium chloride, 10% (*v/v*) glycerol, pH 7.5) until eluate was negative for protein content. Eluted protein was loaded onto a 5 mL Hi-Trap Q HP column connected to an AKTA FPLC system. A gradient of 0 to 1 M sodium chloride was applied over 30 col vol (150 mL) of Anion Exchange Buffer I (50 mM sodium phosphate, 10% (*v/v*) glycerol) at flow rate of 5 mL/ min. Fractions (2.5 mL) were collected and analyzed by SDS-PAGE. Fractions containing the target proteins (*DEBS2-$His_6$, DEBS2(Mod3 $AT^0$)-$His_6$* eluting at ~330-430 mM sodium chloride; *DEBS3-$His_6$, DEBS3(Mod5 $AT^0$)-$His_6$, and DEBS3(Mod6 $AT^0$)-$His_6$* eluting at ~370-500 mM sodium chloride) were pooled and concentrated sing a 100 kDa MWCO Amicon Ultra spin concentrator. Purified protein was flash-frozen in liquid $N_2$ before storing at -80 ºC.

*$His_6$-$LDD_{DEBS}$, $Mod1_{DEBS}$-$His_6$, $Mod2_{DEBS}$-$His_6$, $His_6$-PrpE, and $His_6$-Epi* [*3*]. Cell pellets were resuspended in 35 mL Lysis Buffer J (50 mM sodium phosphate, 10 mM imidazole, 450 mM NaCl, 10% (*v/v*) glycerol, pH 7.6) containing 1 EDTA-Free Protease Inhibitor tablet (Pierce) per L of cell culture. The cells were lysed by sonication and centrifuged at 20,000 × *g* for 60 min at 4 ºC. The cleared cell lysates were added to Ni-NTA agarose resin (1 mL/L cell culture) and incubated at 4 °C for 1 h to bind His-tagged protein. The resin was loaded into a Kimble-Kontes Flex column and washed with 20 col vol of Lysis Buffer J and 10 col vol of Wash Buffer G. Proteins were eluted with 8 col vol of Elution Buffer H. The eluate was further purified by anion-exchange on a Hi-Trap Q column using a gradient from 0 to 500 mM sodium chloride in Anion Exchange Buffer I on an AKTA FPLC system. Fractions were collected and analyzed by SDS-PAGE. Fractions containing the target protein were pooled and concentrated using a 100 kD MWCO Amicon Ultra spin concentrator. Purified protein was flash-frozen in liquid $N_2$ before storing at -80 ºC.

**Synthesis of N-acetylcysteamine thioester of (2*S*,3*R*)-2-methyl-3-hydroxypentanic acid (NDK-SNAC).** The preparation of NDK-SNAC was adapted from literature protocol [*7*,*8*]. Ethyl-2-methyl-3-oxopentanoate (1.5 mmol) was dissolved in 10 mL of water and saponified in aqueous sodium hydroxide (10 M, 180 µL; 1.8 mmol). The reaction was stirred overnight at room temperature before pH was adjusted to <1 and reaction was extracted with 2 × 35 mL of ethyl acetate. The organic layers were combined, dried over anhydrous magnesium sulfate, and evaporated to dryness by rotary evaporation.

The resulting 2-methyl-3-oxopentanoic acid (0.85 mmol) was dissolved in 10 mL of dichloromethane on ice. After 3 mins, N-acetylcysteamine (86 µL, 0.8 mmol, 0.95 eq) and DMAP (8 pellets) were added to the reaction. After 5 mins, ethyl diisopropyl carbodiimide (244 mg, 1.275 mmol; 1.5 eq) was added. The reaction was removed from ice and stirred at room temperature overnight. The reaction was extracted with 2 × 20 mL saturated aqueous ammonium chloride and 2 × 20 mL and saturated aqueous sodium chloride. The aqueous layers were combined and back extracted with 3 × 20 mL dichloromethane. All organic fractions were combined, dried over anhydrous magnesium sulfate, and evaporated to dryness by rotary evaporation. The resulting 2-methyl-3-oxopentanoyl-SNAC product was confirmed by $^1$H-NMR and stored in solid form at -80 ºC. $^1$H NMR (400 MHz, $CDCl_3$=7.26 ppm) δ 5.79 (s, N-H), 3.77-3.82 (m, 1H), 3.42-3.49 (m, 2H), 3.03-3.12 (m, 2H), 2.50-2.62 (m, 2H), 1.97 (s, 3H), 1.20 (d, J = 6.8 Hz, 3H,), 1.07 (t, J = 7.2 Hz, 3H).

2-Methyl-3-oxopentanoyl-SNAC was enzymatically reduced with the ketoreductase domain from the first module of the pikromycin synthase ($KR_{PIKSMod1}$) and NADPH. This reaction (3 mL) contained 50 mM 2-methyl-3-oxopentanoyl-SNAC (1 M stock in DMSO), 300 mM D-glucose, 100 µM $NADP^+$, 20 U/mL glucose-1-dehydrogenase, and 15 µM $Piks_{Mod1}$KR in 300 mM HEPES, 100 mM sodium chloride, 10% (*v/v*) glycerol, pH 7.5. After stirring at room temperature overnight, thin-layer chromatography analysis of crude reaction mixture in 100% ethyl acetate solvent system revealed a product spot at $R_f$ ~ 0.3. Saturated aqueous sodium chloride (3 mL) was then added to the reaction mixture, which was extracted with 3 × 9 mL of ethyl acetate. The organic layer was dried over anhydrous magnesium sulfate and evaporated to dryness by rotary evaporation. Product was confirmed by $^1$H-NMR and stored as 1 M solution in 25% (*v/v*) DMSO at -20 ºC. $^1$H NMR (400 MHz, $CDCl_3$=7.26 ppm) δ 3.81-3.84 (m, 1H), 3.43-3.47 (m, 2H), .99-3.05 (m, 2H), 2.69-2.75

(m, 1H), 2.62 (s, O-H), 1.97 (s, 3H), 1.43-1.57 (m, 2H), 1.23 (d, J = 7.2 Hz, 3H,), 0.98 (t, J = 7.2 Hz, 3H).

**Synthesis of fluoromalonate.** Methanolic sodium hydroxide (2 M, 10.5 mL; 21 mmol) was added to 9:1 (*v/v*) dichloromethane:methanol (96 mL). Diethylfluoromalonate (1.5 mL, 9.6 mmol) was added and saponified with stirring at room temperature for 1-3 h. Disodium fluoromalonate was isolated by filtration through a Büchner funnel fitted with a filter paper and left to air-dry.

**Synthesis of fluoromalonyl-CoA.** Fluoromalonyl-CoA was prepared enzymatically from fluoromalonate and coenzyme A using the malonyl-CoA synthetase from *Streptomyces coelicolor* (MatB) [*6*]. The reaction mixture (5 mL) contained 20 mM fluoromalonate, 4 mM coenzyme A, 5 mM ATP, 20 mM phosphoenol pyruvate, 5 mM TCEP, 10 mM magnesium chloride, 144 U/mL pyruvate kinase, 80 U/mL adenylate kinase, and 20 µM MatB in 200 mM sodium phosphate, pH 7.5. The reaction was incubated at 37 ºC for 1 h. Enzymes were removed by filtration through Nanosep with 10K Omega membrane spin column before reaction was lyophilized overnight. The residue from lyophilization was resuspended in water (2 mL) and stored frozen at -80 ºC until purification. The resuspended crude reaction residue (0.5 mL per injection) was purified on an Eclipse XDB-C18 column (5 µm, 9.4 × 250 mm, room temperature; Agilent) using a linear gradient from 0 to 10% acetonitrile over 30 min with 10 mM ammonium formate as the aqueous mobile phase at flow rate of 4 mL/min. Fractions (2 mL) were screened using an Agilent 1290 UPLC connected to a 6460 Triple Quadrupole (QQQ) mass spectrometer using multiple reaction monitoring (MRM) in positive ionization mode (transition, collision energy, fragmentation voltage: fluoromalonyl-coA (872 → 365, 135, 35)). Fractions were analyzed on a Poroshell HPH-C18 column (2.7 µm, 2.1 × 100 mm, room temperature; Agilent) with a linear gradient from 0 to 20 % acetonitrile over 6 min after an initial hold at 0 % acetonitrile for 30 s with 10 mM ammonium formate as the aqueous mobile phase at flow rate of 0.6 mL/min. Fractions containing pure fluoromalonyl-CoA were pooled and lyophilized overnight. The residue was re-dissolved in water and stored at -80 ºC. The concentration of fluoromalonyl-CoA was estimated using $A_{260}$ ($\varepsilon_{260}$ = 15,400 $M^{-1}$ $cm^{-1}$).

#### Extended Data Figures

**Extended Data Figure 1. Engineering a fluoromalonyl-CoA specific *trans*-AT.** (A) Active site architecture of DszAT. Investigation of the DszAT active site (PDB ID 6APG) reveals F190 and L87 residues near substrate binding pocket. The two residues have no direct role in binding or catalysis but may affect substrate binding and activation and were thus targeted for engineering [*9*]. (B) Mechanism of transacylation reaction. The first step in transacylation reaction involves nucleophilic addition by active site serine to carbonyl carbon, leading to the formation of tetrahedral intermediate, which collapses to form product. Both the nucleophilicity of substrate and the stability of intermediate are enhanced by the presence of fluorine atom at the α position as fluorine prefers to be bonded to an $sp^3$ carbon over an $sp^2$ carbon [*10*].

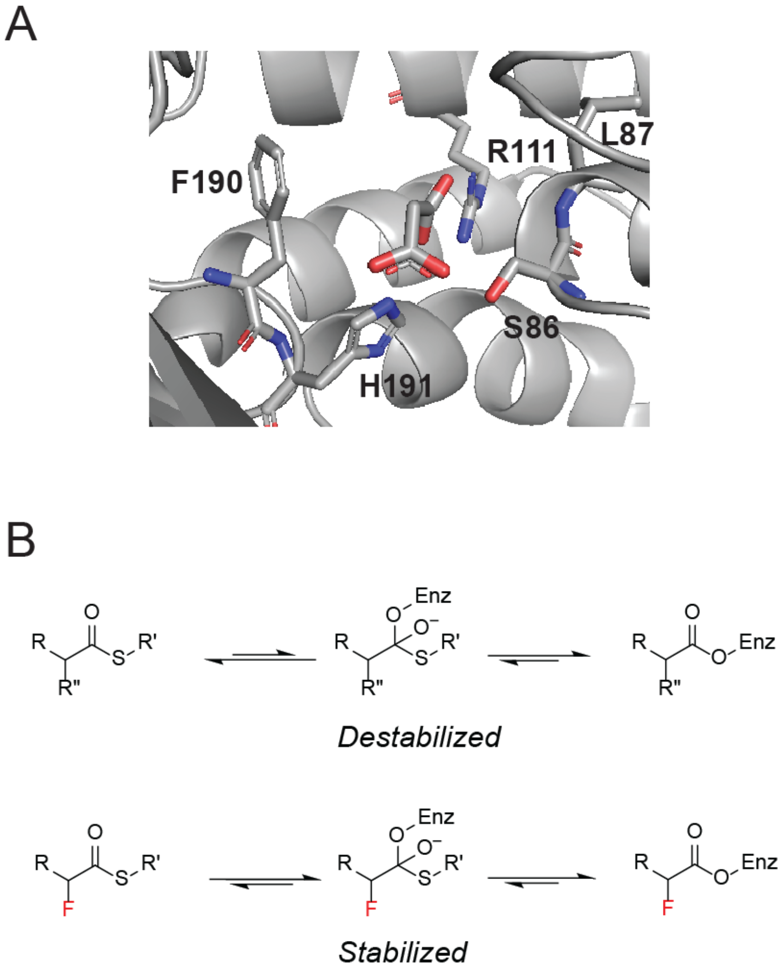

**Extended Data Figure 2. Triketide lactone formation assay.** Formation of triketide lactone (TKL) from the N-acetylcysteamine thioester of (2*S*,3*R*)-2-methyl-3-hydroxypentanic acid (NDK-SNAC) and carboxyacyl-CoA extender unit is initiated when NDK-SNAC self-acylates the active site of the KS domain, mimicking the inter-modular transfer process. The *trans*-AT protein selects the extender unit and transacylates the carboxyacyl group to the phosphopantetheinyl arm of the ACP domain. The KS domain then catalyzes carbon-carbon bond formation through decarboxylative Claisen condensation reaction to produce triketide product. The triketide product is cyclized by the TE domain and become triketide lactone. Alternatively, the triketide Claisen product can be back transferred to the active site of the KS domain to undergo another extension, forming tetraketide pyrone product (TTKP). Carboxyacyl-CoA may be provided as pure extender unit or generated *in situ* from a dicarboxylate precursor by malonyl-CoA synthetase MatB. AMP produced by MatB is regenerated to ATP through an enzyme cascade involving adenylate cyclase (AK) and pyruvate kinase (PK), which derives energy from two molecules of phosphoenolpyruvate (PEP) to produce one molecule of ATP from AMP.

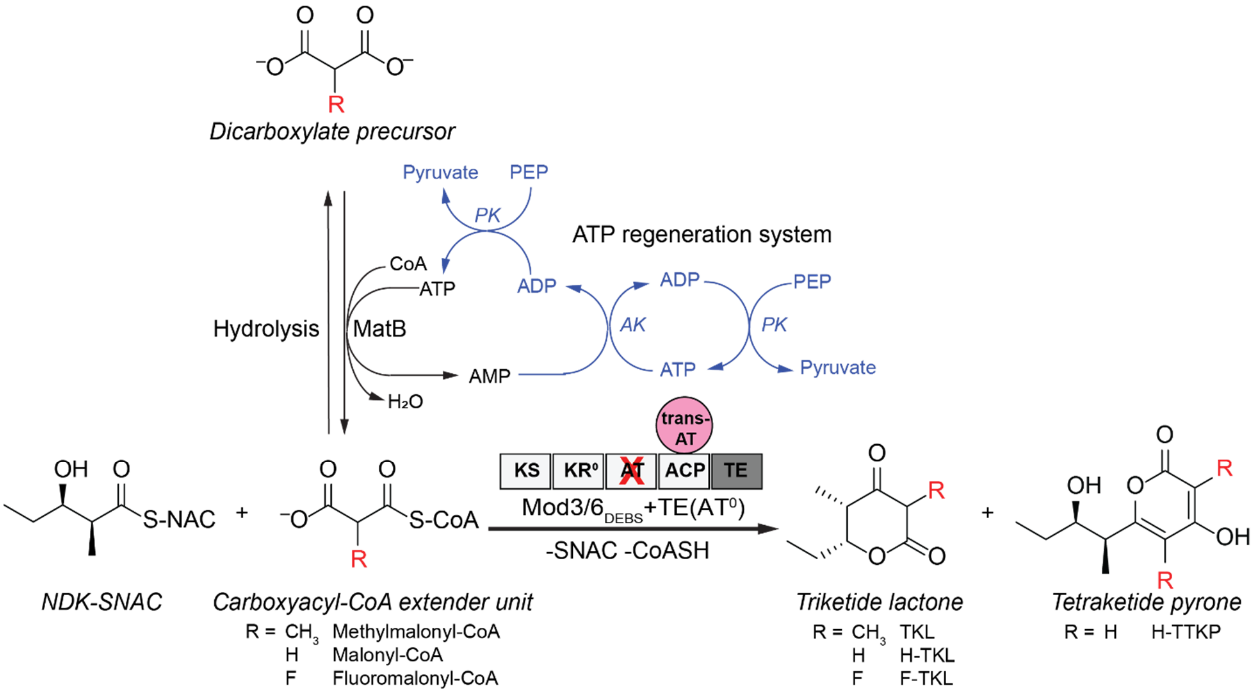

**Extended Data Figure 3. *trans*-AT library screening.** (A) Lysate of *E. coli* expressing following DszAT mutants were screened through the triketide lactone production assay with purified Mod3$_{DEBS}$+TE(AT$^0$) protein, malonate or fluoromalonate extender unit, malonate-coeznzyme A ligase MatB, and NDK-SNAC: **1**, F190A; **2**, F190C; **3**, F190D; **4**, F190E; **5**, F190G; **6**, F190H; **7**, F190I; **8**, F190K; **9**, F190L; **10**, F190M; **11**, F190N; **12**, F190P; **13**, F190Q; **14**, F190R; **15**, F190S; **16**, F190T; **17**, F190V; **18**, F190W; **19**, F190Y; **21**, F190G L87V; **22**, F190G L87A; **23**, F190I L87V; **24**, F190I L87A; **25**, F190P L87A; **26**, F190S L87V; **27**, F190S L87A; **28**, F190T L87V; **29**, F190T L87A; **30**, F190V L87V; **31**, F190V L87A. Product formation was monitored by LC-QQQ using negative ionization mode (transition): F-TKL (173→59), H-TKL (155→97), H-TTKP (197→95), TKL (169→111). The amounts of F-TKL (red), H-TKL (black), and H-TTKP (grey) products were determined by integrating extracted ion counts for the relevant species and are shown normalized by the amount of corresponding product produced by wild-type enzyme. H-TTKP arises from two chain extensions with malonyl-CoA. Mutants that showed potentially increased selectivity towards fluoromalonyl-CoA (marked with asterisks) compared to wild-type enzyme were selected for *in vitro* screening. (B) Selected DszAT mutants from lysate screen (F190G, F190T, F190V, F190P L87A, and F190I L87A) were overexpressed, purified, and assessed through *in vitro* triketide lactone experiment with purified Mod3$_{DEBS}$+TE(AT$^0$) protein, MatB, and NDK-SNAC for the production H-TKL and H-TTKP from malonyl-CoA (top panels), F-TKL from fluoromalonyl-CoA (bottom left), and TKL from methylmalonyl-CoA (bottom right). The amounts of products were determined as in (A). Data are mean of technical replicates (n = 2).

A

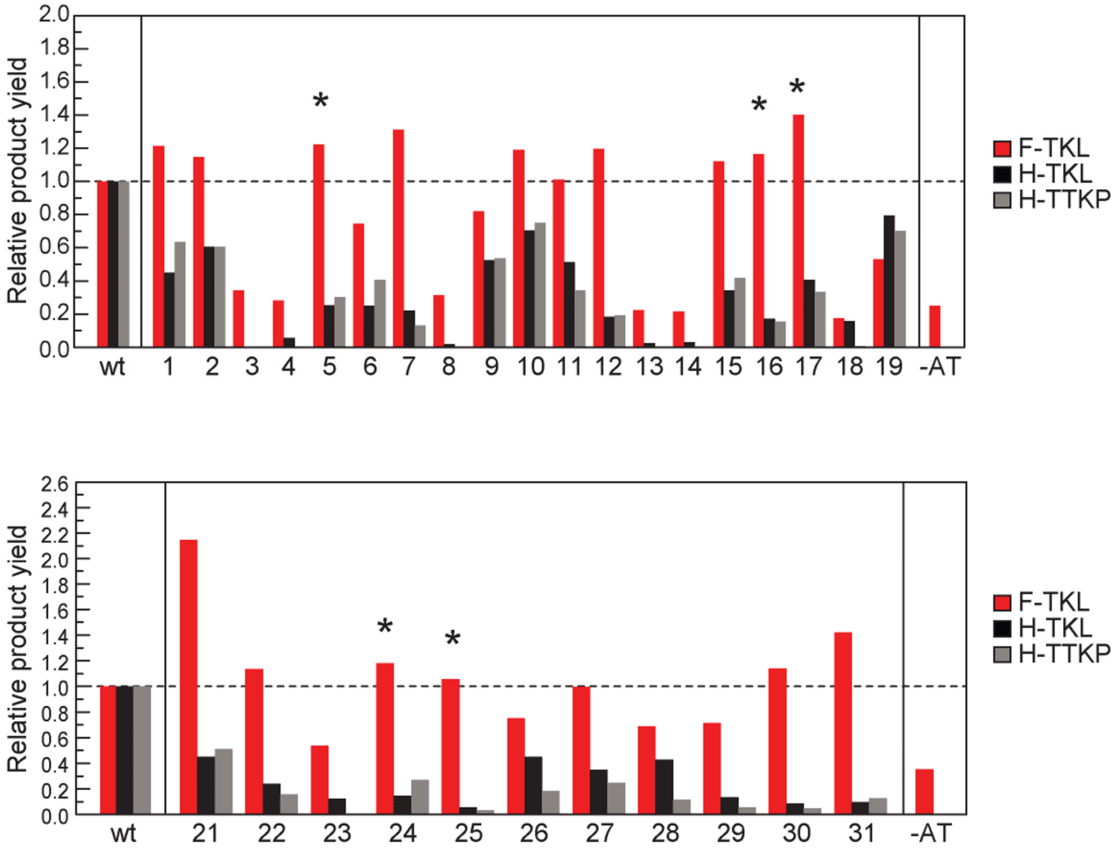

B

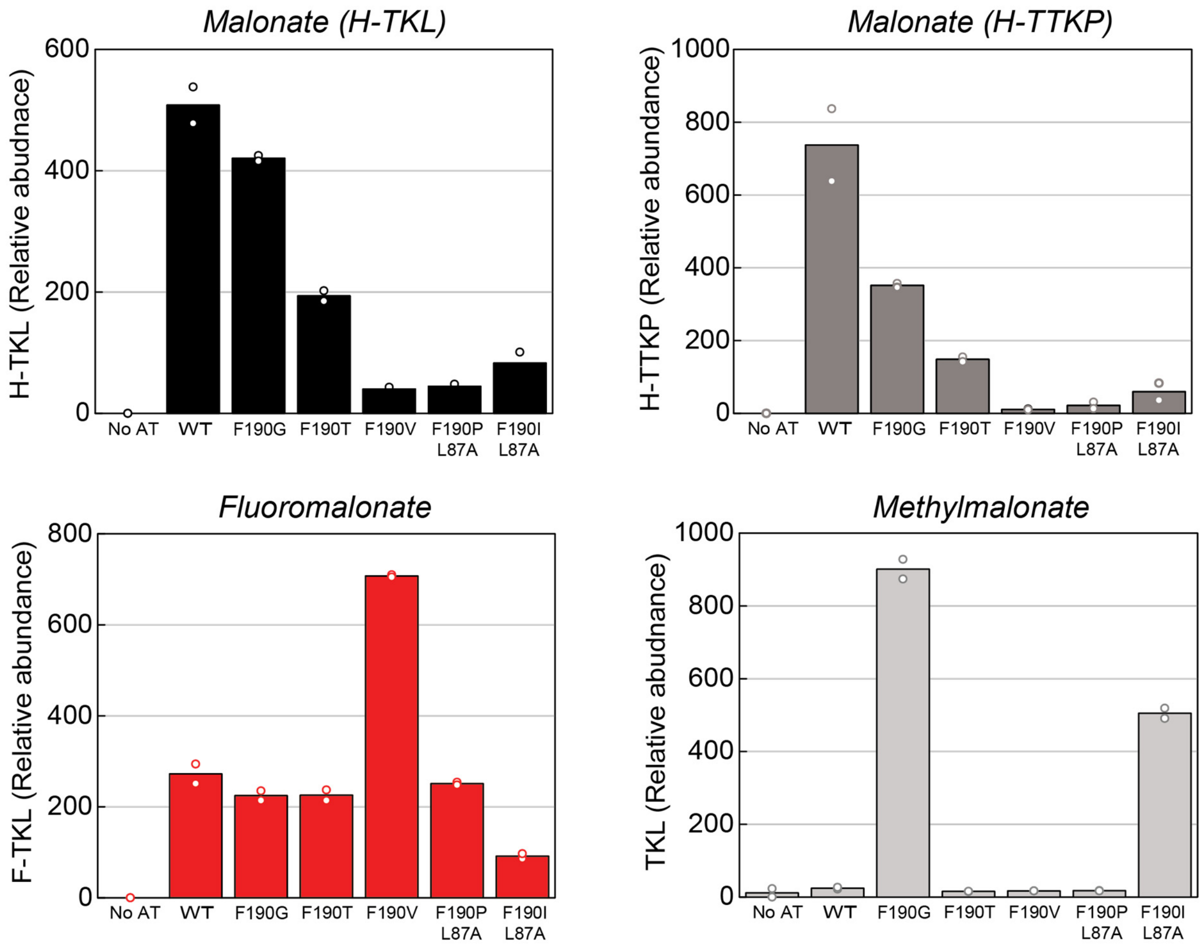
Malonate (H-TKL)
H-TKL (Relative abudnace)
600
400
200
0
No AT
WT
F190G
F190T
F190V
F190P L87A
F190I L87A
Malonate (H-TTKP)
H-TTKP (Relative abundance)
1000
800
600
400
200
0
No AT
WT
F190G
F190T
F190V
F190P L87A
F190I L87A
Fluoromalonate
F-TKL (Relative abundance)
800
600
400
200
0
No AT
WT
F190G
F190T
F190V
F190P L87A
F190I L87A
Methylmalonate
TKL (Relative abundance)
1000
800
600
400
200
0
No AT
WT
F190G
F190T
F190V
F190P L87A
F190I L87A

**Extended Data Figure 4. Characterization of DszAT F190V.** (A) SDS-PAGE of purified proteins used for kinetics and end-point analysis of wild-type and F190V DszAT. **1**, wild-type DszAT; **2**, F190V DszAT; **3**, ACP$_{DEBSMod6}$; **4**, Mod3$_{DEBS}$+TE(AT$^0$); **5**, Mod6$_{DEBS}$+TE(AT$^0$); **6**, MatB; **7,** Epi. (B) Steady-state kinetic analysis of transacylation reaction of malonyl-CoA (left), methylmalonyl-CoA (middle), and fluoromalonyl-CoA (right) by wild-type (black) and F190V (red) DszAT with 75 uM of ACP$_{DEBSMod6}$ as acyl accepter. Transacylation activity was measured by α-ketoglutarate dehydrogenase coupled assay monitoring CoA release. The dose-response curves for transacylation of methylmalonyl-CoA by WT and F190V DszAT were fit with Michaelis-Menten equation. The dose-response curves for WT and F190V DszAT transacylation with malonyl- and fluoromalonyl-CoA exhibited sigmoidal behavior with evidence of inhibition at high substrate concentration and were therefore fit with the Hill equation modified for substrate inhibition. Data shown are mean ± s.d. of technical replicates (n = 3). (C) Steady-state kinetic analysis of hydrolysis reaction of fluoromalonyl-CoA by wild-type and F190V DszAT. Hydrolytic activity was measured by DTNB assay monitoring CoA release. The dose-response curves were obtained from non-linear fitting of data to Michaelis-Menten equation. Data points are mean ± s.d. of technical replicates (n = 3).

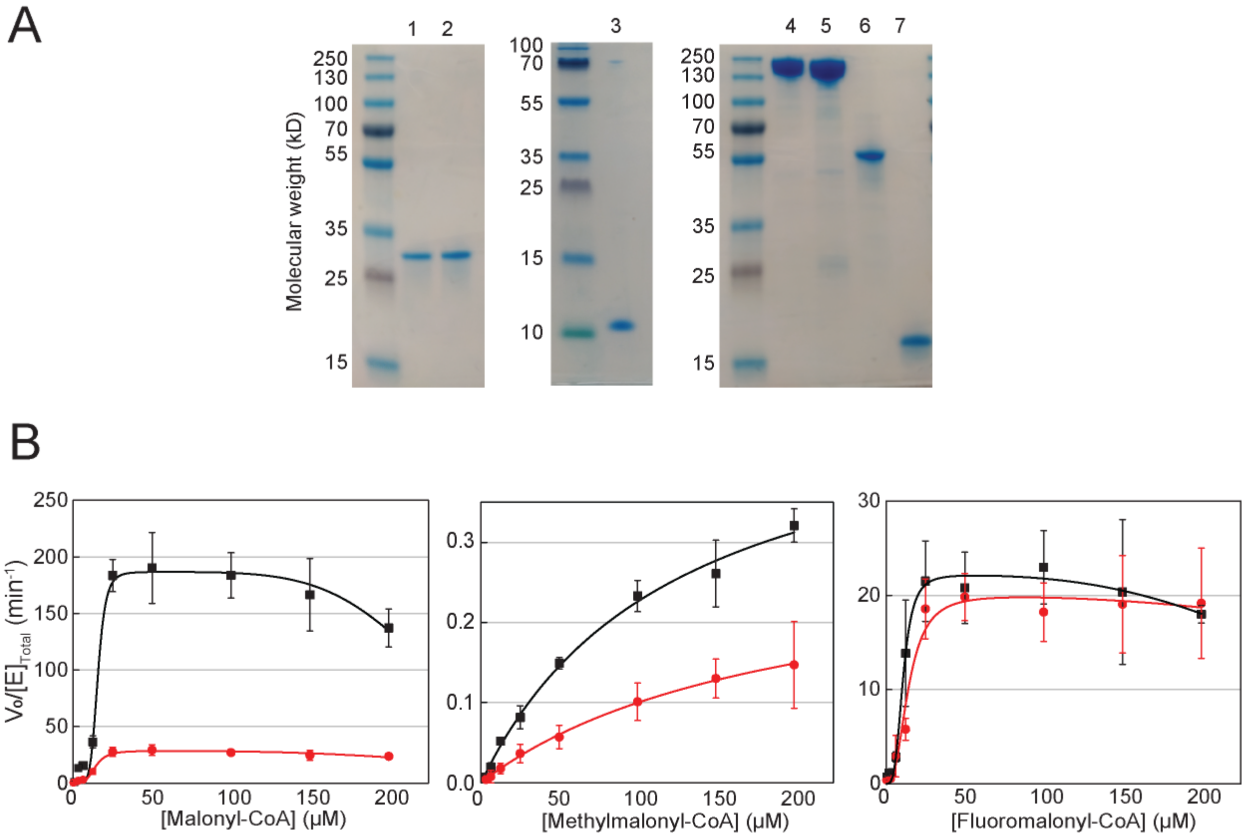

C

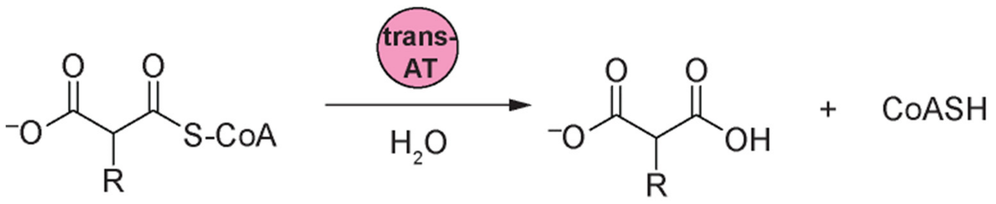


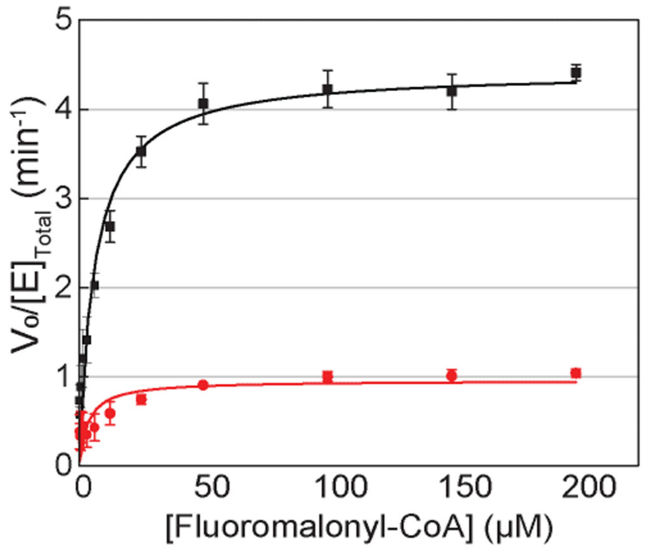


| DszAT | $k_{cat}$ ($min^{-1}$) | $K_M$ (μM) | $k_{cat}/K_M$ ($min^{-1}μM^{-1}$) |
|---|---|---|---|
| Wild-type | 4.4 ± 0.1 | 6.1 ± 0.7 | 0.72 ± 0.1 |
| F190V | 0.96 ± 0.06 | 3.5 ± 1.1 | 0.28 ± 0.11 |

**Extended Data Figure 5. SDS-PAGE of purified proteins used for enzymatic generation of fluorodesmethyl 6dEB analogs.** **1**, PrpE; **2**, Epi; **3**, MatB; **4**, wild-type DszAT; **5**, F190V DszAT; **6**, $LDD_{DBES}$; **7**, $Mod1_{DBES}$; **8**, $Mod2_{DBES}$; **9**, DEBS2; **10**, DEBS2(Mod3 $AT^0$); **11**, DEBS3; **12**, DEBS3(Mod5 $AT^0$); and **13**, DEBS3(Mod6 $AT^0$).

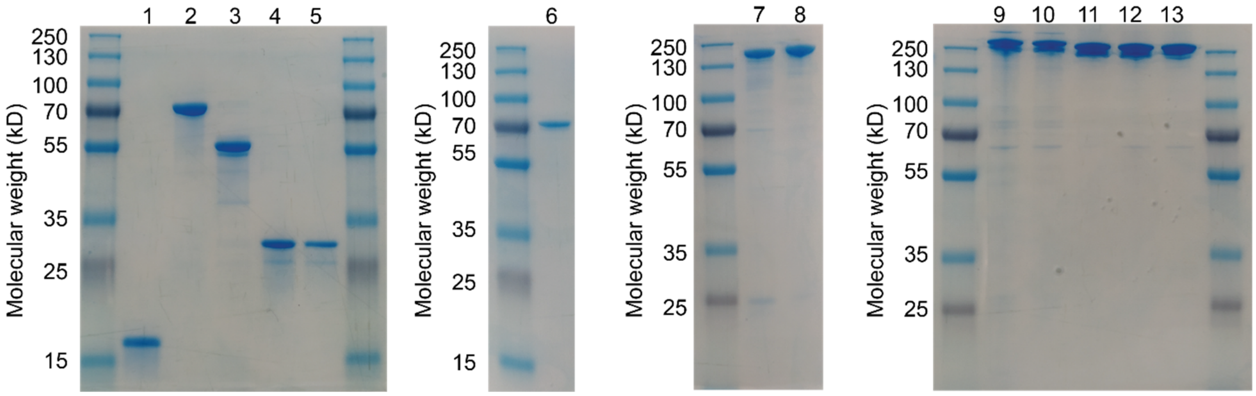

**Extended Data Figure 6. Generation of 2-fluoro-2-desmethyl-6dEB analog through *in vitro* reconstitution of Mod6 AT$^0$ DEBS.** (A) Extracted ion chromatograms showing exact mass of *in vitro* production of monofluorinated desmethyl 6dEB by Mod6 AT$^0$ DEBS complemented with WT DszAT (left) or F190V DszAT (right). Reactions containing 2 μM each of $LDD_{DEBS}$, $Mod1_{DEBS}$, $Mod2_{DEBS}$, DEBS2, and DEBS3(Mod6 AT$^0$), 6 μM WT or F190V DszAT, 2 mM methylmalonate, and when used 10 mM fluoromalonate (+Fmal, red) were analyzed by LC-QTOF. Chromatograms are representative of at least 3 technical replicates. (B) Fragmentation pattern of the monofluorinated desmethyl 6dEB analog produced by Mod6 AT$^0$ DEBS complemented by WT DszAT or F190V DszAT is consistent with 2-fluoro-2-desmethyl 6dEB, which is expected when fluoromalonyl-CoA is incorporated by module 6. Fragmentation pattern shows presence of ions in A, B, and D families (*Extended Data Fig. 7*) with mass shifts according to a substitution of $-CH_3$ with -F group on carbon 2, 4, 6, or 8 of 6dEB. Observed loss of HF from daughter ions expected to contain fluorine substitution further supports the presence of fluorine in the molecule. Notably, the most prominent fragment in 6dEB MS/MS spectrum, the C ion, was entirely missing in the MS/MS spectrum of the monofluorinated desmethyl 6dEB analog [*11*]. This lack of detectable fragment prevents narrowing down the definite location of the fluorine substitution. However, the suppression of ion C was similarly observed when malonate was provided to the system in place of fluoromalonate (*Extended Data Fig. 8*) and is consistent with the reactivity of 6dEB analog with a modification near the C3 position. The lack of methyl substituent at C2 as in 2-fluoro-2-methyl 6dEB would reduce the reactivity of A ion with respect to the hydride shift from C3 to C9 necessary to form C ion. Spectra are representative of at least 3 replicates (nd, not detected). (C) Relative amounts of 2-fluoro-2-desmethyl 6dEB and 6dEB by complementation of Mod6 AT$^0$ DEBS by WT or F190V DszAT. The amounts of products were determined by integrating extracted ion counts for the relevant species monitored by LC-QQQ (transition): 6dEB (369.2→239.1) and 2-fluoro-2-desmethyl 6dEB (373.2 → 275.1). Although the absolute amounts of the two observed products vary among replicates; F190V DszAT reactions consistently produce higher ratio of fluorinated to non-fluorinated product than WT DszAT.

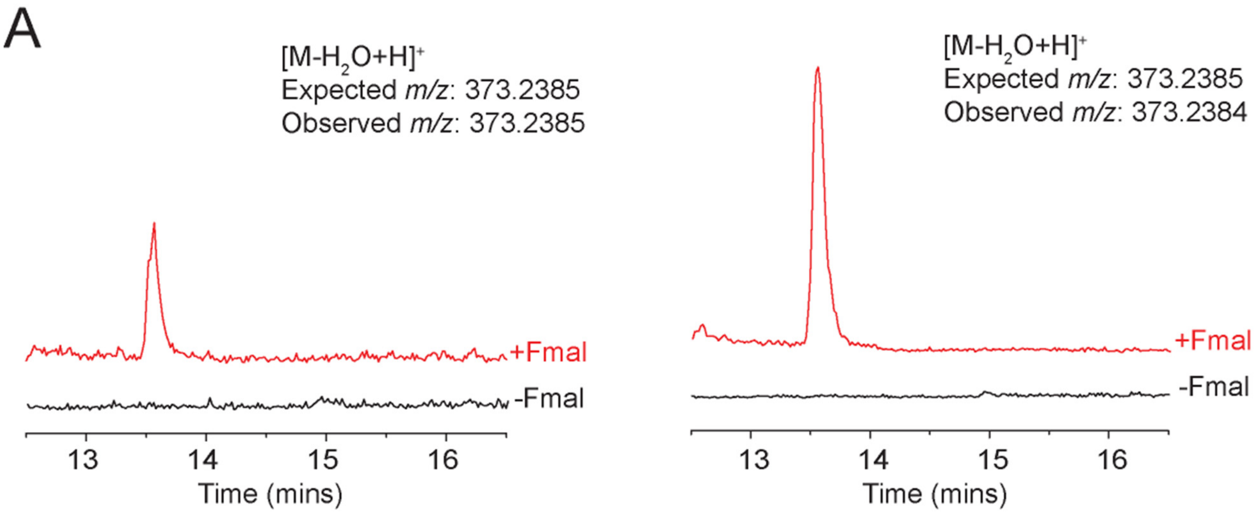

B

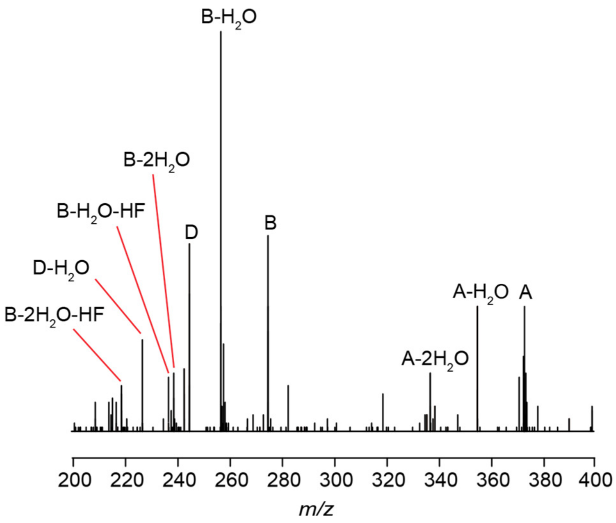


| Ions | Mol. Formula | *m/z* (Exp) | *m/z* (Obs) |
|---|---|---|---|
| A | $C_{20}H_{34}FO_5^+$ | 373.2385 | 373.2372 |
| A-$H_2O$ | $C_{20}H_{32}FO_4^+$ | 355.2279 | 355.2305 |
| A-2$H_2O$ | $C_{20}H_{30}FO_3^+$ | 337.2173 | 337.2124 |
| A-2$H_2O$-HF | $C_{20}H_{29}O_3^+$ | 317.2111 | nd |
| B | $C_{14}H_{24}FO_4^+$ | 275.1653 | 275.1675 |
| B-$H_2O$ | $C_{14}H_{22}FO_3^+$ | 257.1547 | 257.1568 |
| B-2$H_2O$ | $C_{14}H_{20}FO_2^+$ | 239.1442 | 239.2423 |
| B-$H_2O$-HF | $C_{14}H_{21}O_3^+$ | 237.1485 | 237.1471 |
| B-2$H_2O$-HF | $C_{14}H_{19}O_2^+$ | 219.1380 | 219.1356 |
| B-3$H_2O$-HF | $C_{14}H_{17}O^+$ | 201.1274 | nd |
| C | $C_{15}H_{27}O_2^+$ | 239.2006 | nd |
| C-$H_2O$ | $C_{15}H_{25}O^+$ | 221.1900 | nd |
| D | $C_{12}H_{18}FO_4^+$ | 245.1184 | 245.1186 |
| D-$H_2O$ | $C_{12}H_{16}FO_3^+$ | 227.1078 | 227.1082 |
| D-2$H_2O$ | $C_{12}H_{14}FO_2^+$ | 209.0972 | nd |
| D-$H_2O$-HF | $C_{12}H_{15}O_2^+$ | 207.1016 | nd |

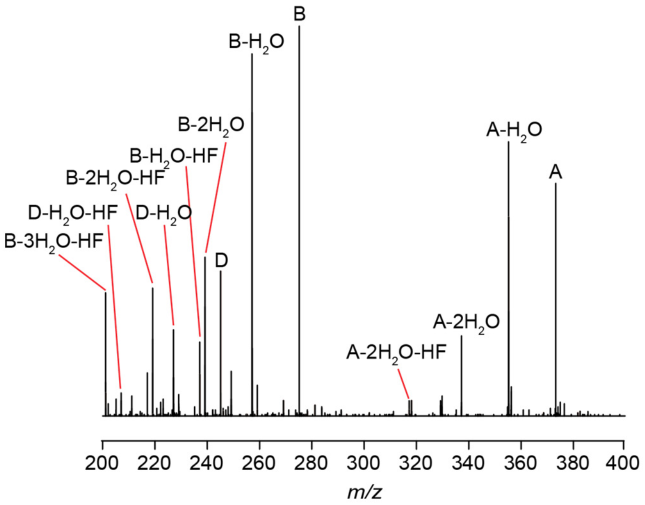


| Ions | Mol. Formula | *m/z* (Exp) | *m/z* (Obs) |
|---|---|---|---|
| A | $C_{20}H_{34}FO_5^+$ | 373.2385 | 373.2373 |
| A-$H_2O$ | $C_{20}H_{32}FO_4^+$ | 355.2279 | 355.2258 |
| A-2$H_2O$ | $C_{20}H_{30}FO_3^+$ | 337.2173 | 337.2156 |
| A-2$H_2O$-HF | $C_{20}H_{29}O_3^+$ | 317.2111 | 317.2136 |
| B | $C_{14}H_{24}FO_4^+$ | 275.1653 | 275.1661 |
| B-$H_2O$ | $C_{14}H_{22}FO_3^+$ | 257.1547 | 257.1548 |
| B-2$H_2O$ | $C_{14}H_{20}FO_2^+$ | 239.1442 | 239.1466 |
| B-$H_2O$-HF | $C_{14}H_{21}O_3^+$ | 237.1485 | 237.1471 |
| B-2$H_2O$-HF | $C_{14}H_{19}O_2^+$ | 219.1380 | 219.1352 |
| B-3$H_2O$-HF | $C_{14}H_{17}O^+$ | 201.1274 | 201.1281 |
| C | $C_{15}H_{27}O_2^+$ | 239.2006 | nd |
| C-$H_2O$ | $C_{15}H_{25}O^+$ | 221.1900 | nd |
| D | $C_{12}H_{18}FO_4^+$ | 245.1184 | 245.1179 |
| D-$H_2O$ | $C_{12}H_{16}FO_3^+$ | 227.1078 | 227.1083 |
| D-2$H_2O$ | $C_{12}H_{14}FO_2^+$ | 209.0972 | nd |
| D-$H_2O$-HF | $C_{12}H_{15}O_2^+$ | 207.1016 | nd |

C

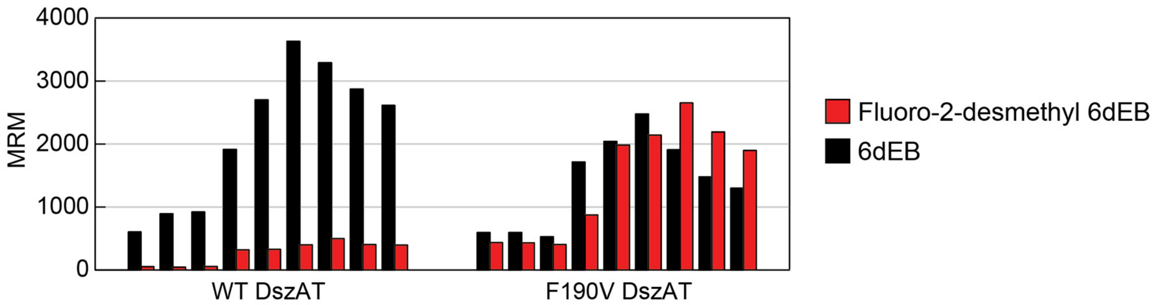

**Extended Data Figure 7. Fragmentation reactions observed of 6dEB and analogs.** 6dEB fragments into 4 daughter fragments shown as A, B, C, and D [*11*]. Each daughter ion is derived from different parts of the carbon skeleton of the parent 6dEB. Module numbering (M1-M6, red) indicates the DEBS module responsible for installing the specified substituent. Carbon numbering (C2-C12, blue) starts at the carboxylic acid with Module 6 being responsible for the substituent at C2. In addition, successive losses of water due to hydroxy groups are observed from all major daughter ions. Exact masses are noted underneath each ion in italic.

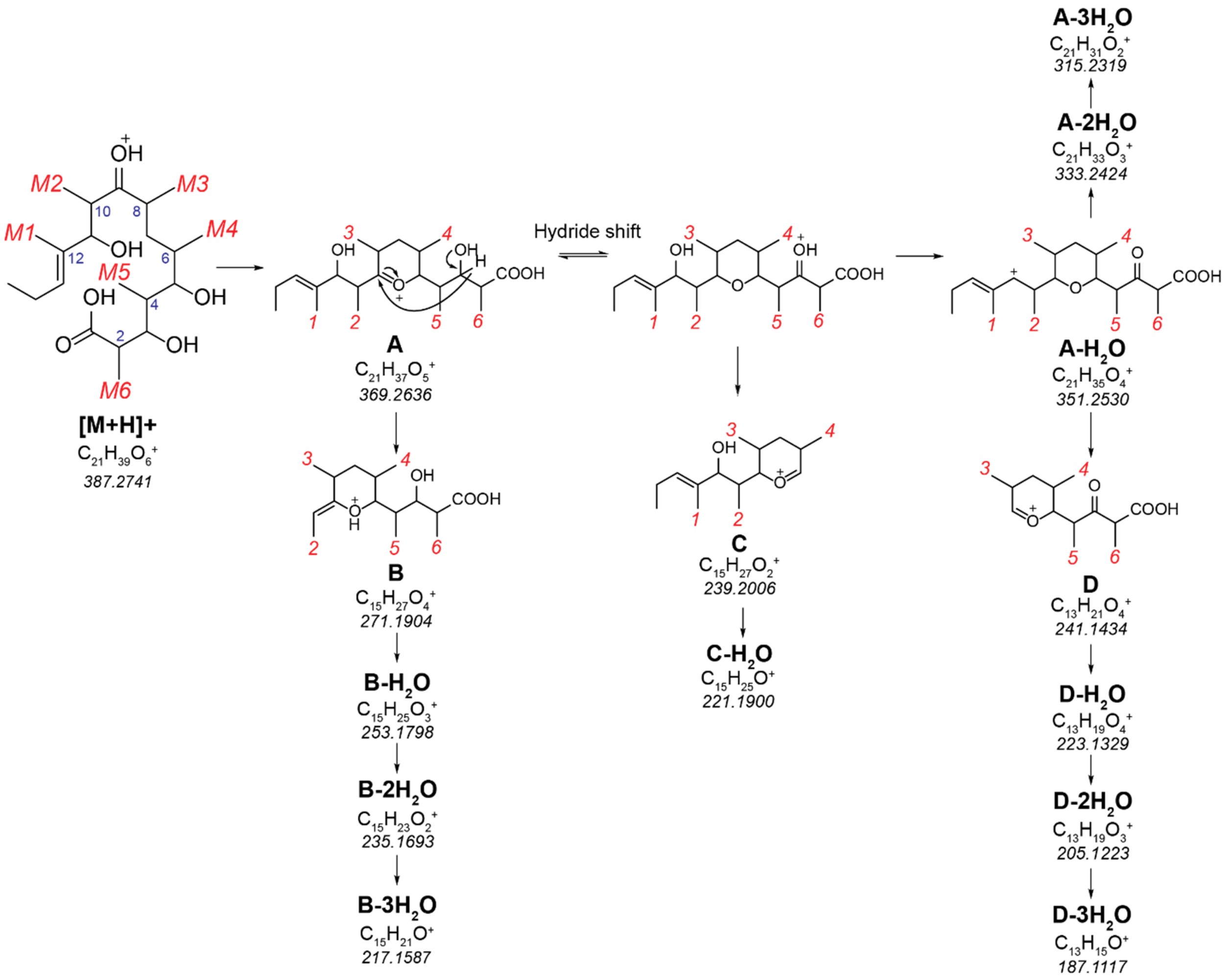

**Extended Data Figure 8. Characterization of 2-desmethyl-6dEB analog through *in vitro* reconstitution of Mod6 AT⁰ DEBS.** (A) Structure of expected product 2-desmethyl 6dEB. (B) Extracted ion chromatograms showing exact mass of *in vitro* production of desmethyl 6dEB by Mod6 $AT^0$ DEBS complemented with WT DszAT. Reactions containing 2 μM each of $LDD_{DEBS}$, $Mod1_{DEBS}$, $Mod2_{DEBS}$, DEBS2, and DEBS3(Mod6 $AT^0$), 6 μM WT DszAT, 2 mM methylmalonate, and when used 10 mM malonate (+Mal, red) were analyzed by LC-QTOF. Chromatograms are representative of at least 3 technical replicates. (C) Fragmentation pattern of the desmethyl 6dEB analog produced by Mod6 $AT^0$ DEBS complemented by WT DszAT is consistent with 2-desmethyl 6dEB. Fragmentation pattern shows presence of ions in A, B, and D families with mass shifts according to a substitution of $-CH_3$ with -H group on carbon 2, 4, 6, or 8 of 6dEB. The C ion, which is the most prominent fragment in 6dEB MS/MS spectrum [*11*] (*Extended Data Fig. 7*) is not observed in the MS/MS spectrum of this desmethyl 6dEB analog, similarly to that observed for the monofluorinated desmethyl 6dEB analog produced by the same enzyme system. Spectrum is representative of at least 3 replicates (nd, not detected).

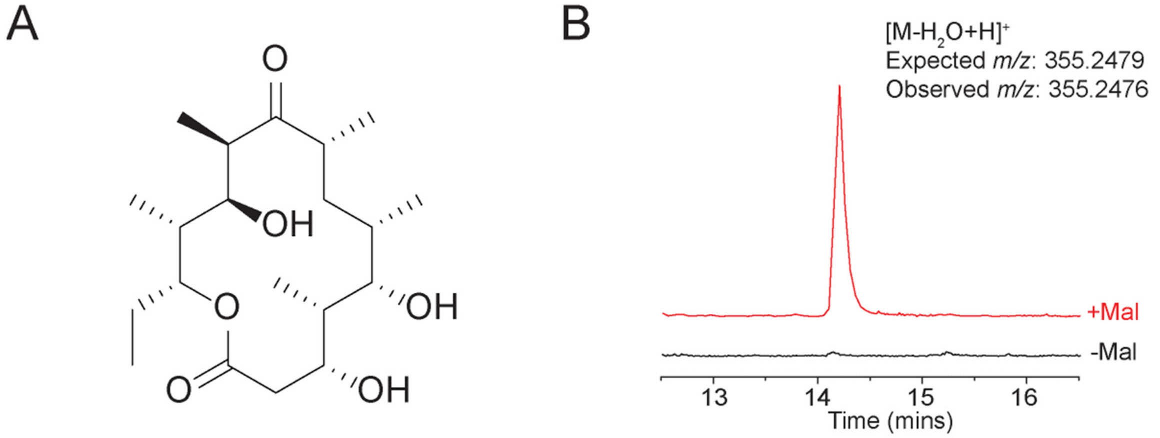


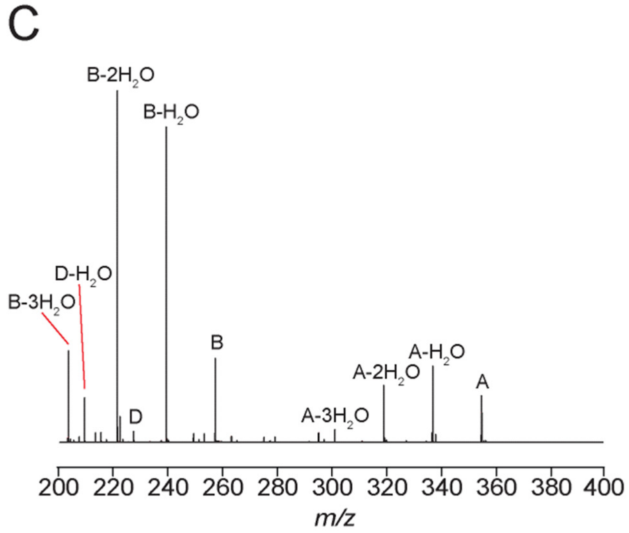


| Ions | Mol. Formula | *m/z* (Exp) | *m/z* (Obs) |
|---|---|---|---|
| A | $C_{20}H_{35}O_5^+$ | 355.2479 | 355.2424 |
| A-$H_2O$ | $C_{20}H_{33}O_4^+$ | 337.2373 | 337.2385 |
| A-2$H_2O$ | $C_{20}H_{31}O_3^+$ | 319.2268 | 319.2294 |
| A-3$H_2O$ | $C_{20}H_{29}O_2^+$ | 301.2162 | 301.2153 |
| B | $C_{14}H_{25}O_4^+$ | 257.1747 | 257.1741 |
| B-$H_2O$ | $C_{14}H_{23}O_3^+$ | 239.1642 | 239.1628 |
| B-2$H_2O$ | $C_{14}H_{21}O_2^+$ | 221.1536 | 221.1518 |
| B-3$H_2O$ | $C_{14}H_{19}O^+$ | 203.1430 | 203.1428 |
| C | $C_{15}H_{27}O_2^+$ | 239.2006 | nd |
| C-$H_2O$ | $C_{15}H_{25}O^+$ | 221.1900 | nd |
| D | $C_{12}H_{19}O_4^+$ | 227.1278 | 227.1314 |
| D-$H_2O$ | $C_{12}H_{17}O_3^+$ | 209.1172 | 209.1163 |
| D-2$H_2O$ | $C_{12}H_{15}O_2^+$ | 191.1067 | nd |
| D-3$H_2O$ | $C_{12}H_{13}O^+$ | 173.0961 | nd |

**Extended Data Figure 9. Generation of 4-fluoro-4-desmethyl-6dEB analog through *in vitro* reconstitution of Mod5 AT$^0$ DEBS.** (A) Extracted ion chromatograms showing exact mass of *in vitro* production of monofluorinated desmethyl 6dEB by Mod5 AT$^0$ DEBS complemented with WT DszAT (left) or F190V DszAT (right). Reactions containing 2 μM each of LDD$_{DEBS}$, Mod1$_{DEBS}$, Mod2$_{DEBS}$, DEBS2, and DEBS3(Mod5 AT$^0$), 6 μM WT or F190V DszAT, 2 mM methylmalonate, and when used 10 mM fluoromalonate (+Fmal, red) were analyzed by LC-QTOF. Chromatograms are representative of at least 3 technical replicates. (B) Fragmentation pattern of the monofluorinated desmethyl 6dEB analog produced by Mod5 AT$^0$ DEBS complemented by WT DszAT or F190V DszAT is consistent with 4-fluoro-4-desmethyl 6dEB, which is expected when fluoromalonyl-CoA is incorporated by Mod5. The observed daughter ion masses are necessarily similar to those of 2-fluoro-2-desmethyl 6dEB observed in Mod6 AT$^0$ DEBS system, due to fluorine substitution in similar positions. In short, mass shifts according to a substitution of $-CH_3$ with -F group on carbon 2, 4, 6, or 8 of 6dEB, loss of HF from daughter ions, and absence of C ion are observed. Spectra are representative of at least 3 replicates (nd, not detected). (C) Fragmentation pattern of the desmethyl 6dEB analog produced by Mod5 AT$^0$ DEBS complemented by WT DszAT in the presence of 10 mM malonate in place of fluoromalonate is consistent with 4-desmethyl 6dEB. Fragmentation pattern is similar to that of 2-desmethyl 6dEB observed in Mod6 AT$^0$ DEBS system due to lack of methyl substitution in similar positions. Spectrum is representative of at least 3 replicates. (D) Relative amounts of 4-fluoro-4-desmethyl 6dEB and 6dEB by complementation of Mod5 AT$^0$ DEBS by WT or F190V DszAT. The amounts of products were determined by integrating extracted ion counts for the relevant species monitored by LC-QQQ (transition): 6dEB (369.2 → 239.1) and 2-fluoro-2-desmethyl 6dEB (373.2 → 275.1).

A

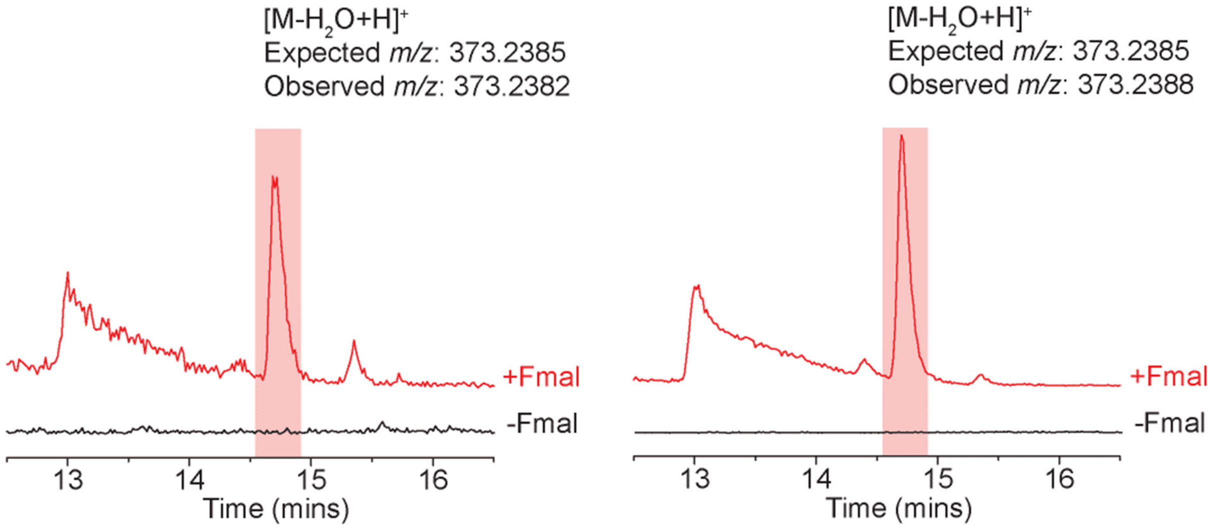

**B**

**Wild-type DszAT**

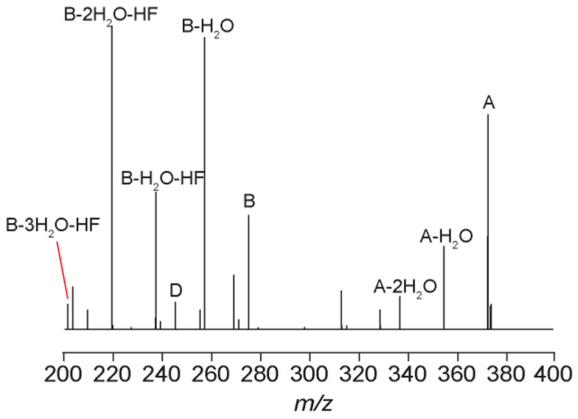


| Ions | Mol. Formula | *m/z* (Exp) | *m/z* (Obs) |
|---|---|---|---|
| A | $C_{20}H_{34}FO_5^+$ | 373.2385 | 373.2372 |
| A-$H_2O$ | $C_{20}H_{32}FO_4^+$ | 355.2279 | 355.2259 |
| A-2$H_2O$ | $C_{20}H_{30}FO_3^+$ | 337.2173 | 337.1691 |
| A-2$H_2O$-HF | $C_{20}H_{29}O_3^+$ | 317.2111 | nd |
| B | $C_{14}H_{24}FO_4^+$ | 275.1653 | 275.1638 |
| B-$H_2O$ | $C_{14}H_{22}FO_3^+$ | 257.1547 | 257.1515 |
| B-2$H_2O$ | $C_{14}H_{20}FO_2^+$ | 239.1442 | nd |
| B-$H_2O$-HF | $C_{14}H_{21}O_3^+$ | 237.1485 | 237.1503 |
| B-2$H_2O$-HF | $C_{14}H_{19}O_2^+$ | 219.1380 | 219.1399 |
| B-3$H_2O$-HF | $C_{14}H_{17}O^+$ | 201.1274 | 201.1129 |
| C | $C_{15}H_{27}O_2^+$ | 239.2006 | nd |
| C-$H_2O$ | $C_{15}H_{25}O^+$ | 221.1900 | nd |
| D | $C_{12}H_{18}FO_4^+$ | 245.1184 | 245.1190 |
| D-$H_2O$ | $C_{12}H_{16}FO_3^+$ | 227.1078 | nd |
| D-2$H_2O$ | $C_{12}H_{14}FO_2^+$ | 209.0972 | nd |
| D-$H_2O$-HF | $C_{12}H_{15}O_2^+$ | 207.1016 | nd |

**F190V DszAT**

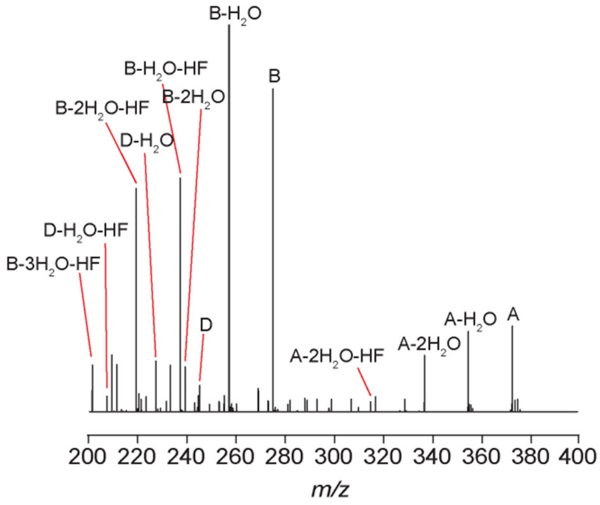


| Ions | Mol. Formula | *m/z* (Exp) | *m/z* (Obs) |
|---|---|---|---|
| A | $C_{20}H_{34}FO_5^+$ | 373.2385 | 373.2383 |
| A-$H_2O$ | $C_{20}H_{32}FO_4^+$ | 355.2279 | 355.2250 |
| A-2$H_2O$ | $C_{20}H_{30}FO_3^+$ | 337.2173 | 337.2116 |
| A-2$H_2O$-HF | $C_{20}H_{29}O_3^+$ | 317.2111 | 317.2102 |
| B | $C_{14}H_{24}FO_4^+$ | 275.1653 | 275.1641 |
| B-$H_2O$ | $C_{14}H_{22}FO_3^+$ | 257.1547 | 257.1548 |
| B-2$H_2O$ | $C_{14}H_{20}FO_2^+$ | 239.1442 | 239.1445 |
| B-$H_2O$-HF | $C_{14}H_{21}O_3^+$ | 237.1485 | 237.1481 |
| B-2$H_2O$-HF | $C_{14}H_{19}O_2^+$ | 219.1380 | 219.1375 |
| B-3$H_2O$-HF | $C_{14}H_{17}O^+$ | 201.1274 | 201.1296 |
| C | $C_{15}H_{27}O_2^+$ | 239.2006 | nd |
| C-$H_2O$ | $C_{15}H_{25}O^+$ | 221.1900 | nd |
| D | $C_{12}H_{18}FO_4^+$ | 245.1184 | 245.1143 |
| D-$H_2O$ | $C_{12}H_{16}FO_3^+$ | 227.1078 | 227.1119 |
| D-2$H_2O$ | $C_{12}H_{14}FO_2^+$ | 209.0972 | nd |
| D-$H_2O$-HF | $C_{12}H_{15}O_2^+$ | 207.1016 | 207.0978 |

**C**

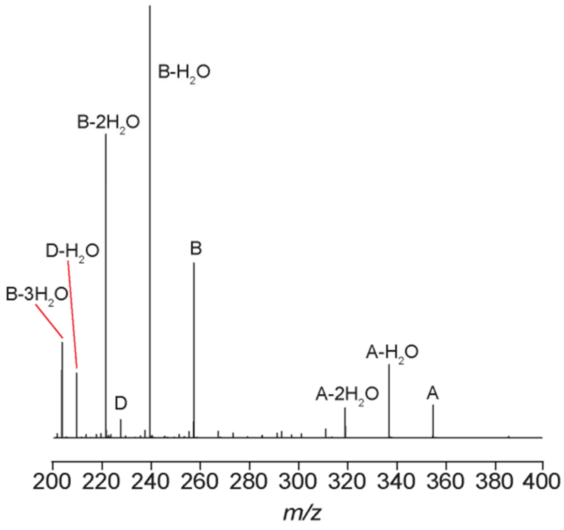


| Ions | Mol. Formula | *m/z* (Exp) | *m/z* (Obs) |
|---|---|---|---|
| A | $C_{20}H_{35}O_5^+$ | 355.2479 | 355.2479 |
| A-$H_2O$ | $C_{20}H_{33}O_4^+$ | 337.2373 | 337.2372 |
| A-2$H_2O$ | $C_{20}H_{31}O_3^+$ | 319.2268 | 319.2271 |
| A-3$H_2O$ | $C_{20}H_{29}O_2^+$ | 301.2162 | nd |
| B | $C_{14}H_{25}O_4^+$ | 257.1747 | 257.1742 |
| B-$H_2O$ | $C_{14}H_{23}O_3^+$ | 239.1642 | 239.1641 |
| B-2$H_2O$ | $C_{14}H_{21}O_2^+$ | 221.1536 | 221.1534 |
| B-3$H_2O$ | $C_{14}H_{19}O^+$ | 203.1430 | 203.1424 |
| C | $C_{15}H_{27}O_2^+$ | 239.2006 | nd |
| C-$H_2O$ | $C_{15}H_{25}O^+$ | 221.1900 | nd |
| D | $C_{12}H_{19}O_4^+$ | 227.1278 | 227.1265 |
| D-$H_2O$ | $C_{12}H_{17}O_3^+$ | 209.1172 | 209.1162 |
| D-2$H_2O$ | $C_{12}H_{15}O_2^+$ | 191.1067 | 191.1073 |
| D-3$H_2O$ | $C_{12}H_{13}O^+$ | 173.0961 | nd |

D

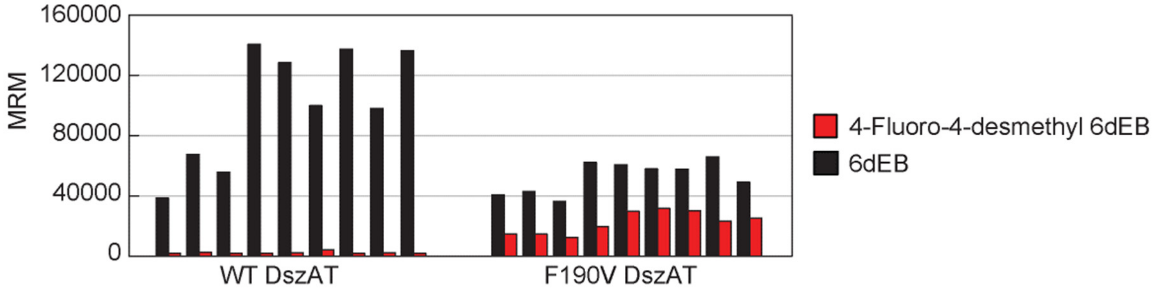

160000
120000
80000
40000
0
MRM
WT DszAT
F190V DszAT
4-Fluoro-4-desmethyl 6dEB
6dEB

**Extended Data Figure 10. Characterization of *E. coli* FabD and its role in polyketide biosynthesis.** (A) Sequence comparison between FabD and DszAT. Conserved functional residues are shown in red; specificity-determining residue are shown in blue; and ACP-interacting residues are shown in green. (B) Structural comparison between Fab (blue) and DszAT (grey). The two proteins have the same overall structure (left, rmsd = 1.7 Å), active site architecture (middle), and ACP-interacting surface (right). Both proteins contain two subdomains with the active site in between. The substrate-binding pocket is largely polar. In both structures, malonate is bound by salt bridge interactions between its β-carboxy group and the side chain amine groups of R117/R111, positioning the carbonyl carbon for nucleophilic attack near the catalytic serine S84/S92, which is found in nearly identical position in both proteins. Notably, in DszAT, F190 is found positioned near the Cα of malonate that may prevent binding of substrate with bulkier substituent such as a methyl group at the α-position. In FabD, this residue is replaced with S200, which may provide more flexibility in terms of substrate binding. Interactions between AT and ACP domains has previously been shown to rely on a small number of charged amino acid residues located near the active sites of the two protein domains[*2, 12*]. R278 and R279 of DszAT interact with E70 of its native protein partner, DSZS ACP1. Similarly, K180 of DszAT interacts with D45 of DSZS ACP1. Structural alignment shows that these positively charged interface residues of DszAT, R278, R279, and K180, correspond to K276, R277, and R190 of FabD, maintaining the charge at the interface for interaction with an ACP partner. Active site, specificity-determining, and interface residues are shown in sticks. (C) Steady-state kinetic characterization of transacylation reaction of malonyl-CoA (left), methylmalonyl-CoA (middle), and fluoromalonyl-CoA (right) catalyzed by FabD with 75 μM of $ACP_{DEBSMod6}$ as acyl accepter. Activity was measured by α-ketoglutarate dehydrogenase coupled assay, monitoring coenzyme A release by NADH fluorescence. Curves were obtained from non-linear curve fitting of data to Hill equation modified for substrate inhibition model. Table contains $k_{cat}$, $K_{0.5}$, $k_{cat}/K_{0.5}$, $K_i$, and $n$ values. Data shown are mean ± s.d. (n=3). Error in $k_{cat}/K_M$ is obtained by propagation from individual kinetic terms**.** (D) *In vitro* triketide lactone assay for FabD complementation of $Mod3_{DEBS}$+TE($AT^0$). Normalized representation of the amount of F-TKL (left), H-TKL (middle), and TKL (right) products are shown for reactions containing 10 μM $Mod3_{DEBS}$+TE($AT^0$)), 30 μM FabD or WT DszAT, 1 mM NDK-SNAC, 1 mM fluoromalonate (left), malonate (middle), or methylmalonate (right), and 1 mM coenzyme A under regenerative condition. Reactions were quenched and analyzed by LC-QQQ after 24 h. The amounts of products were determined by integrating extracted ion counts for the relevant species (transition): F-TKL (175 → 157), H-TKL (157 → 139), and TKL (171 → 153). Data are mean of technical replicates (n = 2).

A

```
FabD   1    MTQFAFVFPGQGSQTVGMLADMAASYPIVEETFAEASAALGYDLWALTQQGPAEELNKTW  60
DszAT  1    --MKAYMFPGQGSQAKGMGRALFDAFPALT---ARADGVLGYSIRALCQDDPDQRLSQTQ  55
                *::*******: **   :  ::* :    * *...***.:.** *:.* : *.:*.

FabD   61   QTQPALLTASVALYRVWQQQGGKAPAMMAGHSLGEYSALVCAGVIDFADAVRLVEMRGKF  120
DszAT  56   FTQPALYVVNALSY-LKRREEEAPPDFLAGHSLGEFSALFAAGVFDFETGLALVKKRGEL  114
             ***** ....  * :...:   .* ::*******:***..***:**  .: **: **::

FabD   121  MQEAVPEGTGAMAAIIGLDDASIAKACEEAAEGQVVSPVNFNSPGQVVIAGHKEAVERAG  180
DszAT  115  MGDA---RGGGMAAVIGLDEERVRELLDQNG-ATAVDIANLNSPSQVVISGAKDEIARLQ  170
            * :*     *.***:****:  : :  :: . . .*. .*:***.****:* *: : *

FabD   181  AACKAAGAKRALPLPVSVPSHCALMKPAADKLAVELAKITFNAPTVPVVNNVDVK-CETN  239
DszAT  171  VPFEAAGAKKYTVLRVSAAFHSRFMRPAMVEFGRFLEGYDFAPPKIPVISNVTARPCKAD  230
            .. :*****.   * **.. *. :*.**  ::.  *    * .*.:**:.** .. *:::

FabD   240  GDAIRDALVRQLYNPVQWTKSVEYMAAQGVEHLYEVGPGKVLTGLTKRIVDTLTASALNE  299
DszAT  231  G--IRAALSEQIASPVRWCESIRYLMGRGVEEFVECGHGIVLTGLYAQI--------RRD  280
            *  ** **  *: .**.* :*: *: ..*** : * * *****  .*         .:

FabD   300  PSAMAAALEL                                                    309
DszAT  281  AQPLVVA---                                                    287
            ...:..*
```

B

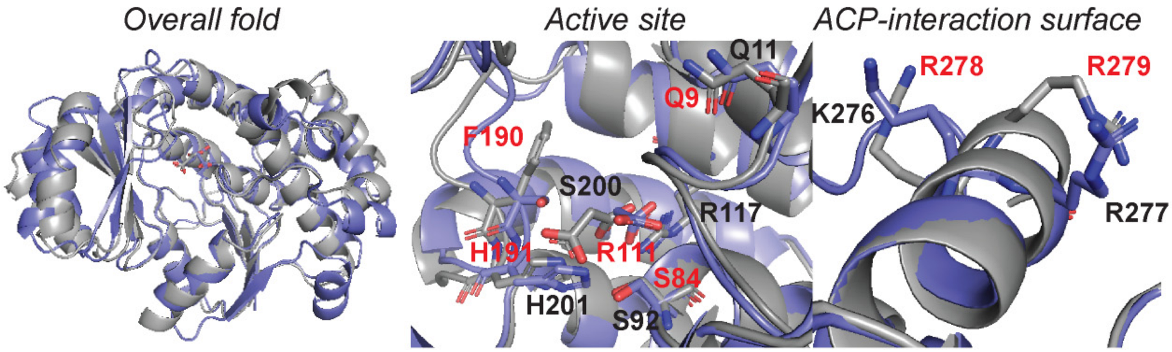

C

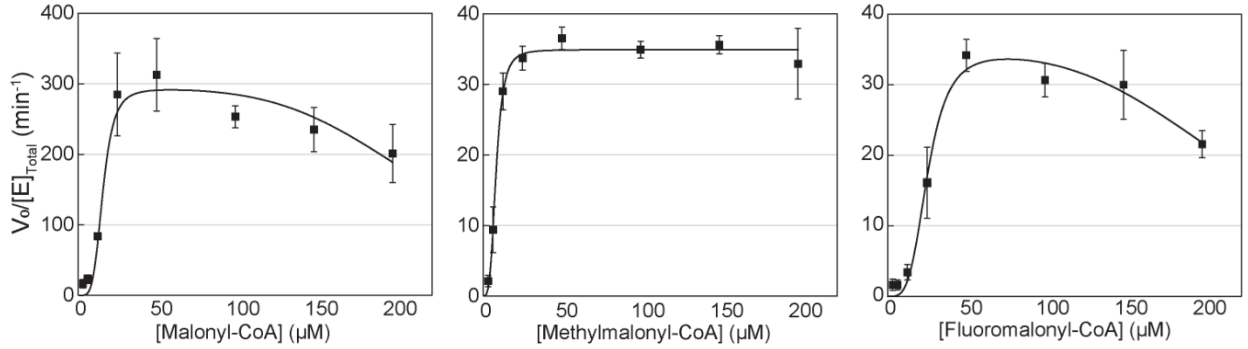


| Substrate | $k_{cat}$ ($min^{-1}$) | $K_{0.5}$ (μM) | $k_{cat}/K_{0.5}$ ($min^{-1}$ $μM^{-1}$) | $K_i$ (μM) | n |
|---|---|---|---|---|---|
| Malonyl-CoA | 293 ± 20 | 15 ± 1.5 | 20 ± 3.3 | 230 ± 19 | 4.3 ± 1.5 |
| Methylmalonyl-CoA | 36 ± 0.9 | 8.3 ± 0.4 | 4.2 ± 0.3 | 429 ± 92 | 3.4 ± 0.5 |
| Fluoromalonyl-CoA | 35 ± 3 | 25 ± 2 | 1.4 ± 0.2 | 230 ± 15 | 3.8 ± 1.2 |

D

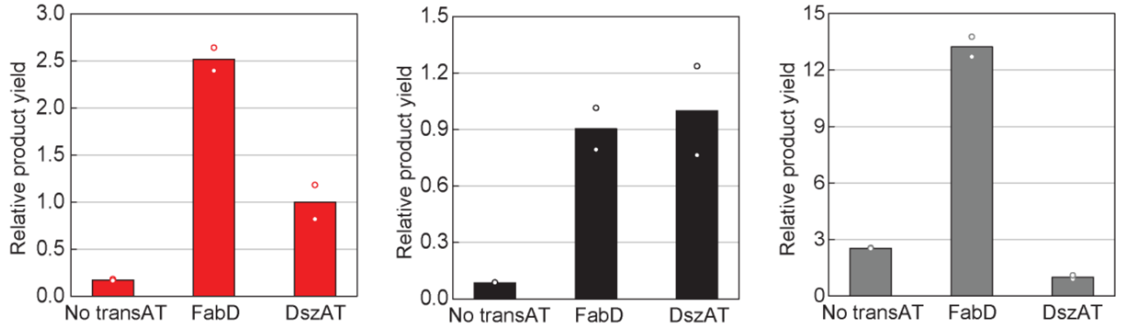

**Extended Data Figure 11. Influence of extender unit availability on the *in vivo* selectivity of chain elongation by single-modular DEBS constructs in engineered *E. coli*.** Concentrated cell suspension of *E. coli* BAP1 expressing Mod3$_{DEBS}$+TE(AT$^0$) (left) or Mod6$_{DEBS}$+TE(AT$^0$) (right), MatB, and MadLM were provided with 1 mM NDK-SNAC and 5 mM fluoromalonate, malonate, and/or methylmalonate and analyzed by LC-QQQ after 24 h. The concentrations of products were determined by integrating extracted ion counts for the relevant species (transition): F-TKL (175 → 157), H-TKL (157 → 139), and TKL (171 → 153), and comparing to external standard curves generated using synthetic standards of the molecules. Data revealed that the product profile follows the corresponding precursor profile, suggesting that the outcome is mainly governed by the provided precursors. Data are mean ± s.d. of biological replicates (n = 3).

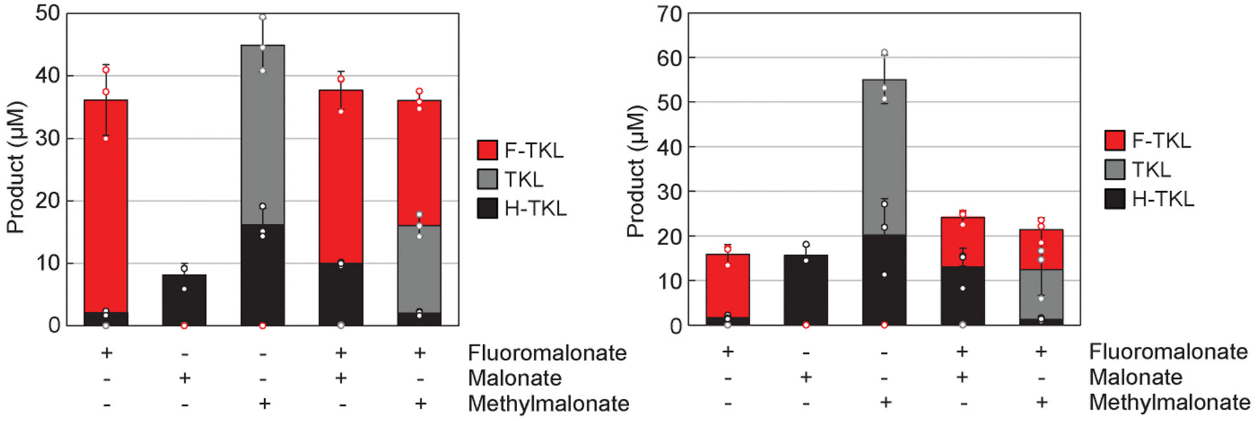

**Extended Data Figure 12. *In vivo* production of desmethyl 6dEB analogs by engineered *E. coli*.** (A) Extracted ion chromatograms of culture media of *E. coli* expressing DEBS, Mod5 $AT^0$ DEBS, or Mod6 $AT^0$ DEBS, showing production of 6dEB. *E. coli* BAP1 harboring variants of pBP130 and pBP144 were grown to $OD_{600}$ = 0.4-0.6 at 37 ºC with shaking at 200 rpm, at which point protein expression was induced with 1 mM IPTG and 0.2% arabinose and cultures provided with 20 mM sodium propionate. Cultures were then incubated with substrates at 22 ºC with shaking at 250 rpm for 1 day, after which, culture media were collected, extracted, and analyzed. 6dEB product was monitored by LC-QQQ using transition 369.1→239.1. Chromatograms are representative of at least 3 biological replicates. (B) Extracted ion chromatograms of culture media of *E. coli* expressing DEBS, Mod5 $AT^0$ DEBS, or Mod6 $AT^0$ DEBS, showing production of desmethyl-6dEB analogs. Samples were prepared as in (A). Desmethyl-6dEB products were monitored by LC-QQQ using transition: 355.2→225.1. Chromatograms are representative of at least 3 biological replicates. (C) Fragmentation patterns of the desmethyl 6dEB analogs produced by *E. coli* harboring Mod5 $AT^0$ DEBS or Mod6 $AT^0$ DEBS. The observed patterns are consistent with those of the desmethyl-6dEB analogs produced by *in vitro* reconstitution of corresponding enzyme systems. Spectra are representative of at least 3 replicates (nd, not detected).

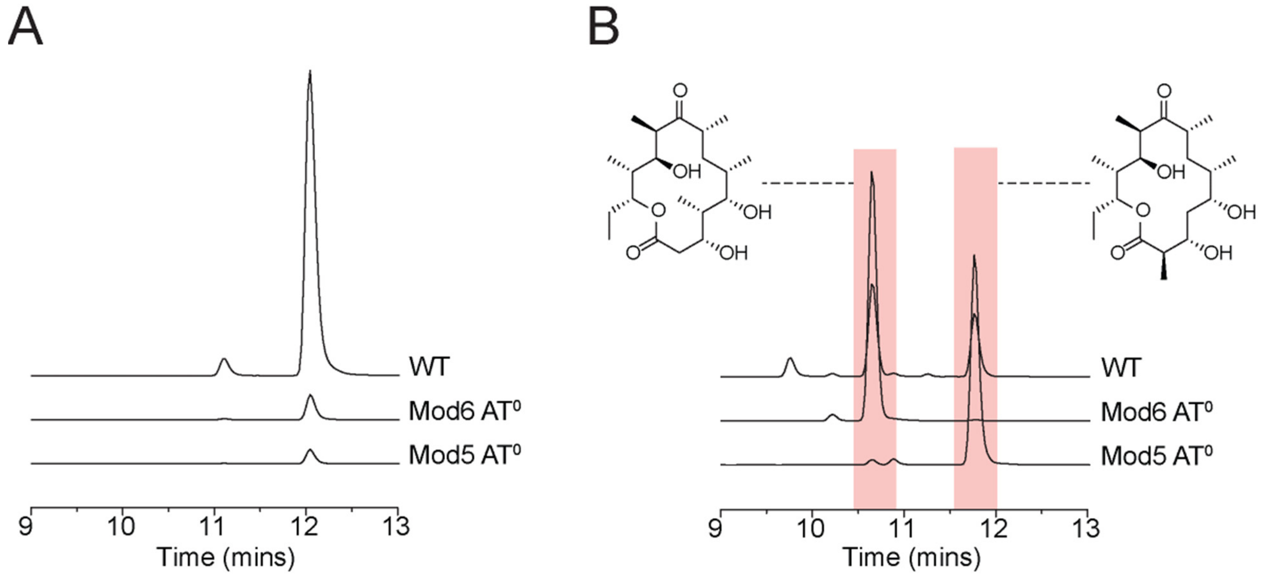

C

**Mod5 AT⁰ DEBS**

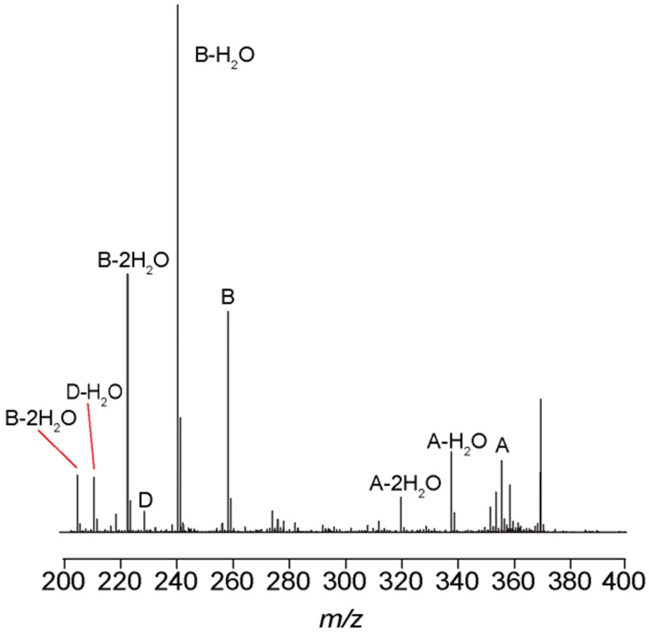


| Ions | Mol. Formula | *m/z* (Exp) | *m/z* (Obs) |
|---|---|---|---|
| A | $C_{20}H_{35}O_5^+$ | 355.2479 | 355.2484 |
| A-$H_2$O | $C_{20}H_{33}O_4^+$ | 337.2373 | 337.2367 |
| A-2$H_2$O | $C_{20}H_{31}O_3^+$ | 319.2268 | 319.2261 |
| A-3$H_2$O | $C_{20}H_{29}O_2^+$ | 301.2162 | nd |
| B | $C_{14}H_{25}O_4^+$ | 257.1747 | 257.1745 |
| B-$H_2$O | $C_{14}H_{23}O_3^+$ | 239.1642 | 239.1640 |
| B-2$H_2$O | $C_{14}H_{21}O_2^+$ | 221.1536 | 221.1533 |
| B-3$H_2$O | $C_{14}H_{19}O^+$ | 203.1430 | 203.1434 |
| C | $C_{15}H_{27}O_2^+$ | 239.2006 | nd |
| C-$H_2$O | $C_{15}H_{25}O^+$ | 221.1900 | nd |
| D | $C_{12}H_{19}O_4^+$ | 227.1278 | 227.1269 |
| D-$H_2$O | $C_{12}H_{17}O_3^+$ | 209.1172 | 209.1170 |
| D-2$H_2$O | $C_{12}H_{15}O_2^+$ | 191.1067 | 191.1069 |
| D-3$H_2$O | $C_{12}H_{13}O^+$ | 173.0961 | 173.0806 |

**Mod6 AT⁰ DEBS**

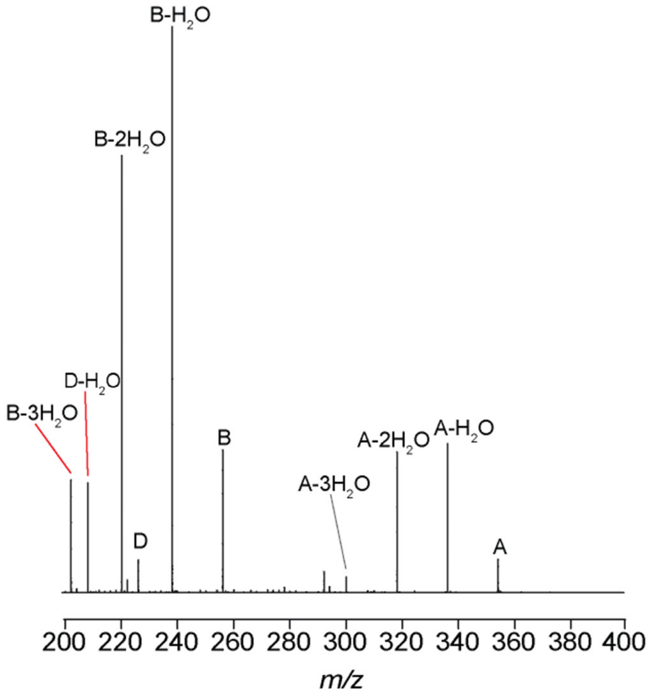


| Ions | Mol. Formula | *m/z* (Exp) | *m/z* (Obs) |
|---|---|---|---|
| A | $C_{20}H_{35}O_5^+$ | 355.2479 | 355.2472 |
| A-$H_2$O | $C_{20}H_{33}O_4^+$ | 337.2373 | 337.2366 |
| A-2$H_2$O | $C_{20}H_{31}O_3^+$ | 319.2268 | 319.2256 |
| A-3$H_2$O | $C_{20}H_{29}O_2^+$ | 301.2162 | 301.2158 |
| B | $C_{14}H_{25}O_4^+$ | 257.1747 | 257.1738 |
| B-$H_2$O | $C_{14}H_{23}O_3^+$ | 239.1642 | 239.1632 |
| B-2$H_2$O | $C_{14}H_{21}O_2^+$ | 221.1536 | 221.1528 |
| B-3$H_2$O | $C_{14}H_{19}O^+$ | 203.1430 | 203.1424 |
| C | $C_{15}H_{27}O_2^+$ | 239.2006 | nd |
| C-$H_2$O | $C_{15}H_{25}O^+$ | 221.1900 | nd |
| D | $C_{12}H_{19}O_4^+$ | 227.1278 | 227.1272 |
| D-$H_2$O | $C_{12}H_{17}O_3^+$ | 209.1172 | 209.1163 |
| D-2$H_2$O | $C_{12}H_{15}O_2^+$ | 191.1067 | 191.1056 |
| D-3$H_2$O | $C_{12}H_{13}O^+$ | 173.0961 | nd |

**Extended Data Fig. 13. *In vivo* production of monofluorinated desmethyl 6dEB analogs by engineered *E. coli*.** (A) Fragmentation pattern and tabulated masses of daughter ions of monofluorinated desmethyl 6dEB analog produced by *E. coli* expressing Mod5 AT$^0$ DEBS or Mod5 AT$^0$ DEBS. The observed patterns are consistent with those of the monofluorinated desmethyl-6dEB analogs produced by *in vitro* reconstitution of corresponding enzyme systems. Spectra are representative of at least 3 biological replicates. (B) $^{19}$F-NMR spectrum of 2-fluoro-2-desmethyl 6dEB isolated from culture media extract of *E. coli* expressing Mod6 AT$^0$ DEBS, MatB, and MadLM. Concentrated cell suspension was provided with 5 mM fluoromalonate and 20 mM propionate and incubated at 22 ºC with shaking at 250 rpm. After 24 h incubation, culture medium was extracted with ethyl acetate, dried via rotary evaporation. 2-fluoro-2-desmethyl 6dEB was then purified from ethyl acetate extract of culture medium through HPLC and fractions were screened using LC-QQQ. Fractions showing presence of 2-fluoro-2-desmethyl 6dEB with transition 373.2 → 275.1 were combined, lyophilized, and resuspended in 50:50 MeOH:$D_2O$ mixture for $^{19}$F-NMR analysis. Spectrum was collected on Bruker AV600 with following parameters: o1p = -200, sw = 200, d1 = 1s, d0 -20, n = 3200. 5-fluorouracil (1 mM) was used as internal standard. Spectrum reveals a set of signals between -195 and -196 ppm displaying a doublet of doublet splitting pattern ($J$ = 54, 12 Hz) consistent with the β-hydroxy-α-fluoro-carbonyl motif expected of 2-fluoro-2-desmethyl 6dEB. The observed chemical shift value is similar to that observed of other compounds with similar α-fluoro-β-hydroxy ester motif [*11*]. Based on vicinal coupling constant ($^3J_{F\text{-}H}$) and known (S)-orientation of hydroxy group on carbon 3, the observed molecule is assigned as 2-(*R*)-fluoro-2-desmethyl 6dEB.

A

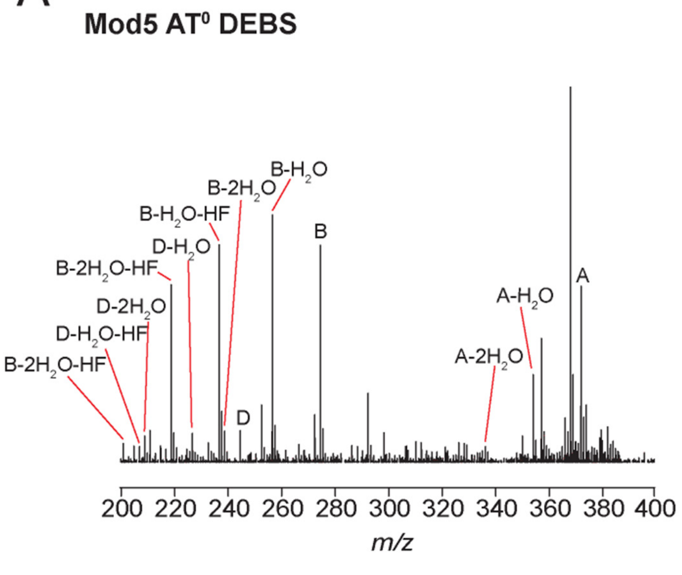


| Ions | Mol. Formula | *m/z* (Exp) | *m/z* (Obs) |
|---|---|---|---|
| A | $C_{20}H_{34}FO_5^+$ | 373.2385 | 373.2386 |
| A-$H_2O$ | $C_{20}H_{32}FO_4^+$ | 355.2279 | 355.2284 |
| A-2$H_2O$ | $C_{20}H_{30}FO_3^+$ | 337.2173 | 337.2164 |
| A-2$H_2O$-HF | $C_{20}H_{29}O_3^+$ | 317.2111 | nd |
| B | $C_{14}H_{24}FO_4^+$ | 275.1653 | 275.1653 |
| B-$H_2O$ | $C_{14}H_{22}FO_3^+$ | 257.1547 | 257.1547 |
| B-2$H_2O$ | $C_{14}H_{20}FO_2^+$ | 239.1442 | 239.1451 |
| B-$H_2O$-HF | $C_{14}H_{21}O_3^+$ | 237.1485 | 237.1487 |
| B-2$H_2O$-HF | $C_{14}H_{19}O_2^+$ | 219.1380 | 219.1377 |
| B-3$H_2O$-HF | $C_{14}H_{17}O^+$ | 201.1274 | 201.1289 |
| C | $C_{15}H_{27}O_2^+$ | 239.2006 | nd |
| C-$H_2O$ | $C_{15}H_{25}O^+$ | 221.1900 | nd |
| D | $C_{12}H_{18}FO_4^+$ | 245.1184 | 245.1180 |
| D-$H_2O$ | $C_{12}H_{16}FO_3^+$ | 227.1078 | 227.1074 |
| D-2$H_2O$ | $C_{12}H_{14}FO_2^+$ | 209.0972 | 209.1535 |
| D-$H_2O$-HF | $C_{12}H_{15}O_2^+$ | 207.1016 | 207.1020 |

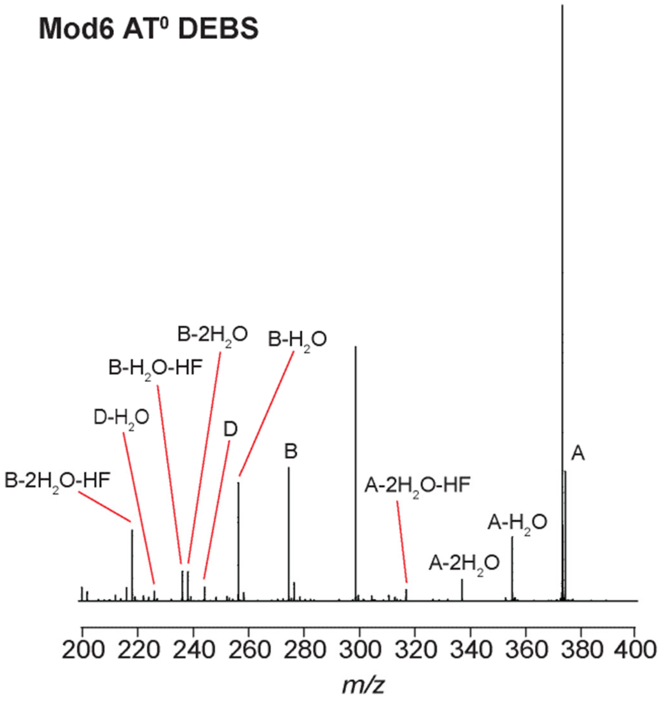


| Ions | Mol. Formula | *m/z* (Exp) | *m/z* (Obs) |
|---|---|---|---|
| A | $C_{20}H_{34}FO_5^+$ | 373.2385 | 373.2366 |
| A-$H_2O$ | $C_{20}H_{32}FO_4^+$ | 355.2279 | 355.2268 |
| A-2$H_2O$ | $C_{20}H_{30}FO_3^+$ | 337.2173 | 337.2170 |
| A-2$H_2O$-HF | $C_{20}H_{29}O_3^+$ | 317.2111 | 317.2129 |
| B | $C_{14}H_{24}FO_4^+$ | 275.1653 | 275.1651 |
| B-$H_2O$ | $C_{14}H_{22}FO_3^+$ | 257.1547 | 257.1537 |
| B-2$H_2O$ | $C_{14}H_{20}FO_2^+$ | 239.1442 | 239.1430 |
| B-$H_2O$-HF | $C_{14}H_{21}O_3^+$ | 237.1485 | 237.1480 |
| B-2$H_2O$-HF | $C_{14}H_{19}O_2^+$ | 219.1380 | 219.1373 |
| B-3$H_2O$-HF | $C_{14}H_{17}O^+$ | 201.1274 | 201.1259 |
| C | $C_{15}H_{27}O_2^+$ | 239.2006 | nd |
| C-$H_2O$ | $C_{15}H_{25}O^+$ | 221.1900 | nd |
| D | $C_{12}H_{18}FO_4^+$ | 245.1184 | 245.1157 |
| D-$H_2O$ | $C_{12}H_{16}FO_3^+$ | 227.1078 | 227.1070 |
| D-2$H_2O$ | $C_{12}H_{14}FO_2^+$ | 209.0972 | nd |
| D-$H_2O$-HF | $C_{12}H_{15}O_2^+$ | 207.1016 | nd |

B

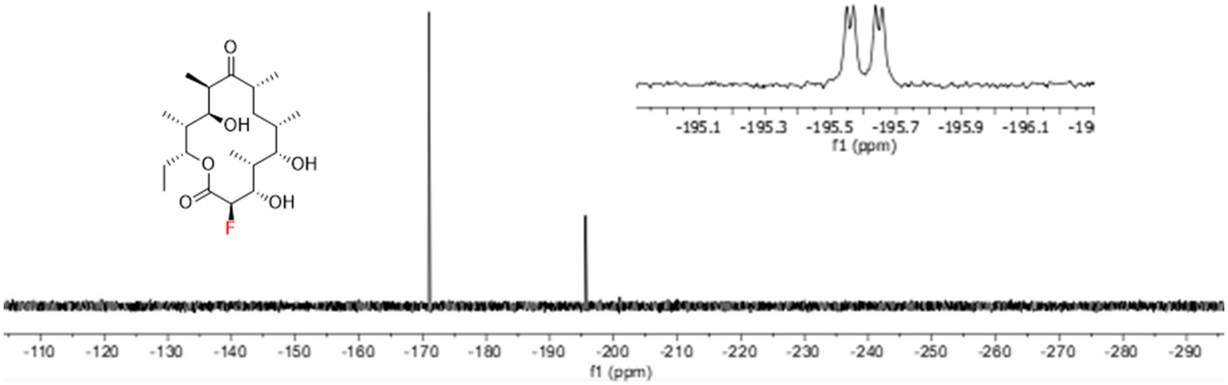

### Supplementary Tables

**Supplementary Table 1. Strains, plasmids, oligonucleotides, and sequences.** (A) Strains (B) Plasmids (C) Oligonucleotides

#### A. Strains

| Strains | Description | Source |
|---|---|---|
| *E. coli* BAP1 | F- ompT gal dcm lon hsdSB(rB- mB-) λ(DE3 [lacI lacUV5-T7 gene1 ind1 sam7 nin5) prpRBCDE (sfp (T7), prpE (T7))] | Ref. 4 |
| *E. coli* BAP1-T1$^R$ | Derived from BAP1, ΔfhuA::FRT, T1, and T5 Phage-resistant | Ref. 5 |
| *E. coli* DH10B-T1$^R$ | F- endA1 recA1 galE15 galK16 nupG rpsL ΔlacX74 Φ80lacZΔM15 araD139 Δ(ara,leu)7697 mcrA Δ(mrr-hsdRMS-mcrBC) λ-, T1 and T5 Phage-resistant | Invitrogen |
| *E. coli* BL21 (DE3)-T1$^R$ | F– ompT gal dcm lon hsdSB(rB- mB-) λ(DE3 [lacI lacUV5-T7 gene1ind1 sam7 nin5]), T1 and T5 Phage-resistant | Novagen |

#### B. Plasmids

| Plasmids | Description | Source |
|---|---|---|
| pFW3 | Cb$^R$; DszAT | Ref. 2 |
| pFW3_F190V | Cb$^R$; F190V DszAT | This study |
| pFW3_F190L | Cb$^R$; F190L DszAT | This study |
| pFW3_F190S | Cb$^R$; F190S DszAT | This study |
| pFW3_F190I | Cb$^R$; F190I DszAT | This study |
| pFW3_F190P | Cb$^R$; F190P DszAT | This study |
| pFW3_F190T | Cb$^R$; F190T DszAT | This study |
| pFW3_F190A | Cb$^R$; F190A DszAT | This study |
| pFW3_F190Y | Cb$^R$; F190Y DszAT | This study |
| pFW3_F190H | Cb$^R$; F190H DszAT | This study |
| pFW3_F190Q | Cb$^R$; F190Q DszAT | This study |
| pFW3_F190N | Cb$^R$; F190N DszAT | This study |
| pFW3_F190K | Cb$^R$; F190K DszAT | This study |
| pFW3_F190D | Cb$^R$; F190D DszAT | This study |
| pFW3_F190E | Cb$^R$; F190E DszAT | This study |
| pFW3_F190C | Cb$^R$; F190C DszAT | This study |
| pFW3_F190W | Cb$^R$; F190W DszAT | This study |
| pFW3_F190M | Cb$^R$; F190M DszAT | This study |
| pFW3_F190G | Cb$^R$; F190G DszAT | This study |
| pFW3_F190R | Cb$^R$; F190G DszAT | This study |
| pFW3_H191A | Cb$^R$; H191A DszAT | This study |
| pFW3_S86C | Cb$^R$; S86C DszAT | This study |
| pFW3_S86D | Cb$^R$; S86D DszAT | This study |
| pFW3_S86E | Cb$^R$; S86E DszAT | This study |
| pFW3_S86D H191A | Cb$^R$; S86D H191A DszAT | This study |
| pFW3_S86E H191A | Cb$^R$; S86E H191A DszAT | This study |
| pFW3_F190G L87V | Cb$^R$; F190G L87V DszAT | This study |
| pFW3_F190G L87A | Cb$^R$; F190G L87A DszAT | This study |

| | | |
|---|---|---|
| pFW3_F190I L87V | $Cb^R$; F190I L87V DszAT | This study |
| pFW3_F190I L87A | $Cb^R$; F190I L87A DszAT | This study |
| pFW3_F190P L87A | $Cb^R$; F190P L87A DszAT | This study |
| pFW3_F190S L87V | $Cb^R$; F190S L87V DszAT | This study |
| pFW3_F190S L87A | $Cb^R$; F190S L87A DszAT | This study |
| pFW3_F190T L87V | $Cb^R$; F190T L87V DszAT | This study |
| pFW3_F190T L87A | $Cb^R$; F190T L87A DszAT | This study |
| pFW3_F190V L87V | $Cb^R$; F190V L87V DszAT | This study |
| pBL12 | $Km^R$; LDD | Ref. 3 |
| pBL13 | $Cb^R$; $Mod1_{DEBS}$ | Ref. 3 |
| pBL36 | $Cb^R$; $Mod2_{DEBS}$ | Ref. 3 |
| pFW98 | $Cb^R$; DEBS2 | Ref. 3 |
| pFW100 | $Cb^R$; DEBS3 | Ref. 3 |
| pET16b-$His_{10}$Pres-$ACP_{DEBSMod6}$ | $Cb^R$; $ACP_{DEBSMod6}$ with N-terminal Prescission-cleavable His tag | This study |
| pAYC136 | $Cb^R$; $Mod3_{DEBS}$+TE($AT^0$) | Ref. 2 |
| pAYC138 | CbR; $Mod6_{DEBS}$+TE($AT^0$) | Ref. 2 |
| pFW98_DEBS2 (Mod3 $AT^0$) | $Cb^R$; DEBS2 (Mod6 $AT^0$; S653A) | This study |
| pFW100_DEBS3 (Mod5 $AT^0$) | $Cb^R$; DEBS3 (Mod5 $AT^0$; S642) | This study |
| pFW100_DEBS3 (Mod6 $AT^0$) | $Cb^R$; DEBS3 (Mod6 $AT^0$; S2107A) | This study |
| p15A-DszAT | $Cm^R$; lacI; pT7 lacO DszAT T7-term; p15A ori | This study |
| p15A-DszATS86A | $Cm^R$; lacI; pT7 lacO S86A DszAT T7-term; p15A ori | This study |
| p15A-DszATF190V | $Cm^R$; lacI; pT7 lacO F190V DszAT T7-term; p15A ori | This study |
| pET21c-FabD | $Cb^R$; FabD from E. coli | This study |
| pET-$Mod3_{DEBS}$ $AT^0ACP^0$ | $Cb^R$; $Mod3_{DEBS}$+TE($AT^0ACP^0$) | Ref. 14 |
| pFmal(MdcF) | $Sp^R$; lacI; AraC; pBAD33 mdcF rrnB T1 T2-term; pT7 lacO matB T7-term; CloDF13 ori | Ref. 15 |
| pFmal(MatC) | $Sp^R$; lacI; AraC; pBAD33 matC rrnB T1 T2-term; pT7 lacO matB T7-term; CloDF13 ori | Ref. 15 |
| pFmal(MadLM) | $Sp^R$; lacI; AraC; pBAD33 madLM rrnB T1 T2-term; pT7 lacO matB T7-term; CloDF13 ori | Ref. 15 |
| pFmal((MadLM, without MatB) | $Sp^R$; lacI; araC; pBAD33 madLM rrnB T1 T2-term; CloDF13 ori. | Ref. 15 |
| pBP130 | $Cb^R$; DEBS2 DEBS3; ColE1 ori | Ref. 4 |
| pBP144 | $Km^R$; DEBS1 pccAB; ColE1 ori | Ref. 4 |
| pBP144_DEBS1 (Mod1 $AT^0$) | $Km^R$; DEBS1 S1181A pccAB; ColE1 ori | This study |
| pBP130_DEBS2 (Mod3 $AT^0$) | $Cb^R$; DEBS2 S653A DEBS3; ColE1 ori | This study |
| pBP130_DEBS3 (Mod5 $AT^0$) | $Cb^R$; DEBS2 DEBS3 S642; ColE1 ori | This study |
| pBP130_DEBS3 (Mod6 $AT^0$) | $Cb^R$; DEBS2 DEBS3 S2107A; ColE1 ori | This study |

#### C. Oligonucleotides

| Name | Sequence |
|---|---|
| DszAT F190 F1 | gccggtgatgccggccacgatgcgtccggcgtagaggatcgagatctcgatcccgcgaaa |
| DszAT F190 R2 | gccaactcagcttcctttcgggctttgttagcagccggatctcagtggtggtggtggtgg |
| DszAT F190L R1 | cggtcgcatgaagcgggaatggagagcggcgctcacgcgcaggactgtgtacttcttcgc |
| DszAT F190I R1 | cggtcgcatgaagcgggaatgaatagcggcgctcacgcgcaggactgtgtacttcttcgc |
| DszAT F190V R1 | cggtcgcatgaagcgggaatgaacagcggcgctcacgcgcaggactgtgtacttcttcgc |
| DszAT F190S R1 | cggtcgcatgaagcgggaatgagaagcggcgctcacgcgcaggactgtgtacttcttcgc |
| DszAT F190P R1 | cggtcgcatgaagcgggaatgaggagcggcgctcacgcgcaggactgtgtacttcttcgc |
| DszAT F190T R1 | cggtcgcatgaagcgggaatgagtagcggcgctcacgcgcaggactgtgtacttcttcgc |
| DszAT F190A R1 | cggtcgcatgaagcgggaatgagcagcggcgctcacgcgcaggactgtgtacttcttcgc |
| DszAT F190Y R1 | cggtcgcatgaagcgggaatgataagcggcgctcacgcgcaggactgtgtacttcttcgc |
| DszAT F190H R1 | cggtcgcatgaagcgggaatgatgagcggcgctcacgcgcaggactgtgtacttcttcgc |
| DszAT F190L R1 | cggtcgcatgaagcgggaatggagagcggcgctcacgcgcaggactgtgtacttcttcgc |
| DszAT F190Q R1 | cggtcgcatgaagcgggaatgttgagcggcgctcacgcgcaggactgtgtacttcttcgc |
| DszAT F190N R1 | cggtcgcatgaagcgggaatgattagcggcgctcacgcgcaggactgtgtacttcttcgc |
| DszAT F190K R1 | cggtcgcatgaagcgggaatgtttagcggcgctcacgcgcaggactgtgtacttcttcgc |
| DszAT F190D R1 | cggtcgcatgaagcgggaatgatcagcggcgctcacgcgcaggactgtgtacttcttcgc |
| DszAT F190E R1 | cggtcgcatgaagcgggaatgttcagcggcgctcacgcgcaggactgtgtacttcttcgc |
| DszAT F190C R1 | cggtcgcatgaagcgggaatgacaagcggcgctcacgcgcaggactgtgtacttcttcgc |
| DszAT F190W R1 | cggtcgcatgaagcgggaatgccaagcggcgctcacgcgcaggactgtgtacttcttcgc |
| DszAT F190M R1 | cggtcgcatgaagcgggaatgtagagcggcgctcacgcgcaggactgtgtacttcttcgc |
| DszAT F190G R1 | cggtcgcatgaagcgggaatgaccagcggcgctcacgcgcaggactgtgtacttcttcgc |
| DszAT F190L F2 | gcgaagaagtacacagtcctgcgcgtgagcgccgctctccattcccgcttcatgcgaccg |
| DszAT F190I F2 | gcgaagaagtacacagtcctgcgcgtgagcgccgctattcattcccgcttcatgcgaccg |
| DszAT F190V F2 | gcgaagaagtacacagtcctgcgcgtgagcgccgctgttcattcccgcttcatgcgaccg |
| DszAT F190S F2 | gcgaagaagtacacagtcctgcgcgtgagcgccgcttctcattcccgcttcatgcgaccg |
| DszAT F190P F2 | gcgaagaagtacacagtcctgcgcgtgagcgccgctcctcattcccgcttcatgcgaccg |
| DszAT F190T F2 | gcgaagaagtacacagtcctgcgcgtgagcgccgctactcattcccgcttcatgcgaccg |
| DszAT F190A F2 | gcgaagaagtacacagtcctgcgcgtgagcgccgctgctcattcccgcttcatgcgaccg |
| DszAT F190Y F2 | gcgaagaagtacacagtcctgcgcgtgagcgccgcttatcattcccgcttcatgcgaccg |
| DszAT F190H F2 | gcgaagaagtacacagtcctgcgcgtgagcgccgctcatcattcccgcttcatgcgaccg |
| DszAT F190Q F2 | gcgaagaagtacacagtcctgcgcgtgagcgccgctcaacattcccgcttcatgcgaccg |
| DszAT F190N F2 | gcgaagaagtacacagtcctgcgcgtgagcgccgctaatcattcccgcttcatgcgaccg |
| DszAT F190K F2 | gcgaagaagtacacagtcctgcgcgtgagcgccgctaaacattcccgcttcatgcgaccg |
| DszAT F190D F2 | gcgaagaagtacacagtcctgcgcgtgagcgccgctgatcattcccgcttcatgcgaccg |
| DszAT F190E F2 | gcgaagaagtacacagtcctgcgcgtgagcgccgctgaacattcccgcttcatgcgaccg |
| DszAT F190C F2 | gcgaagaagtacacagtcctgcgcgtgagcgccgcttgtcattcccgcttcatgcgaccg |
| DszAT F190W F2 | gcgaagaagtacacagtcctgcgcgtgagcgccgcttggcattcccgcttcatgcgaccg |
| DszAT F190M F2 | gcgaagaagtacacagtcctgcgcgtgagcgccgctatgcattcccgcttcatgcgaccg |
| DszAT F190G F2 | gcgaagaagtacacagtcctgcgcgtgagcgccgctggtcattcccgcttcatgcgaccg |
| DszAT F190R F1 | atgcgtccggcgtagaggatcgagatctcgatcccgcgaaattaatacgactcactataggggaattgtgagc ggataacaattcccctc |
| DszAT F190R R1 | cggtcgcatgaagcgggaatggcgagcggcgctcacgcgcaggactgtgtacttcttcgc |
| DszAT F190R F2 | gcgaagaagtacacagtcctgcgcgtgagcgccgctcgccattcccgcttcatgcgaccg |
| DszAT F190R R2 | gccaactcagcttcctttcgggctttgttagcagccggatctcagtggtggtggtggtgg |
| DszAT S86 F1 | ccacggggcctgccaccatacccacgccgaaacaagcgctcatgagcccgaagtggcgag |
| DszAT S86 R2 | gccaactcagcttcctttcgggctttgttagcagccggatctcagtggtggtggtggtgg |

| | |
|---|---|
| DszAT S86C R1 | aacagggcgctgaactcgcccagacagtggccggccaggaaatcgg |
| DszAT S86A R1 | aacagggcgctgaactcgcccagacagtggccggccaggaaatcgg |
| DszAT S86D R1 | aacagggcgctgaactcgcccagatcgtggccggccaggaaatcgg |
| DszAT S86E R1 | aacagggcgctgaactcgcccagttcgtggccggccaggaaatcgg |
| DszAT S86C F2 | ccgatttcctggccggccactgtctgggcgagttcagcgccctgtt |
| DszAT S86A F2 | ccgatttcctggccggccacgctctgggcgagttcagcgccctgtt |
| DszAT S86D F2 | ccgatttcctggccggccacgatctgggcgagttcagcgccctgtt |
| DszAT S86E F2 | ccgatttcctggccggccacgaactgggcgagttcagcgccctgtt |
| DszAT L87 F1 | ccacggggcctgccaccatacccacgccgaaacaagcgctcatgagcccgaagtggcgag |
| DszAT L87 R2 | gccaactcagcttcctttcgggctttgttagcagccggatctcagtggtggtggtggtgg |
| DszAT L87V R1 | aacagggcgctgaactcgccaaccgagtggccggccaggaaatcgg |
| DszAT L87A R1 | aacagggcgctgaactcgccagccgagtggccggccaggaaatcgg |
| DszAT L87V F2 | ccgatttcctggccggccactcggttggcgagttcagcgccctgtt |
| DszAT L87A F2 | ccgatttcctggccggccactcggctggcgagttcagcgccctgtt |
| DszAT H191A R1 | gaccatcgccggtcgcatgaagcgggaagcgaaagcggcgctcacgcgcaggactgtgta |
| DszAT H191A F2 | tacacagtcctgcgcgtgagcgccgctttcgcttcccgcttcatgcgaccggcgatggtc |
| PresACP6_Fwd3 | catctagaagtgctttttcagggcccgcatatggcggccccggcgcgggagatgacgtcgcaggagt |
| PresACP6_Rev | tcctttcgggctttgttagcagccggatcctcagagctgctgtcctatgtggt |
| DszAT F190 F1 | gccggtgatgccggccacgatgcgtccggcgtagaggatcgagatctcgatcccgcgaaa |
| DszAT F190V R1 | cggtcgcatgaagcgggaatgaacagcggcgctcacgcgcaggactgtgtacttcttcgc |
| DszAT F190V F2 | gcgaagaagtacacagtcctgcgcgtgagcgccgctgttcattcccgcttcatgcgaccg |
| DszAT F190 R2 | gccaactcagcttcctttcgggctttgttagcagccggatctcagtggtggtggtggtgg |
| pACYC184_CmOperon_pFW100_M5_Fwd_PacI | cggccgggtcgcctactgcctgggcctgga ttaattaa ttacgccccgccctgccact |
| pACYC184_CmOperon_pFW100_M5_Rev_SpeI | agtcgacctccacgccctgcgcgtac actagt aacaggagggacagctgatagaaacaga |
| pFW100_M5AT0_SfiI_F | tcgcctactgcctgggcctgga |
| pFW100_M5AT0_SfiI_R | atctcgccctg cgc gtggccga |
| pFW100_M5AT0_BsiWI_F | tcggccac gcg cagggcgagat |
| pFW100_M5AT0_BsiWI_R | acctccacgccctgcgcgtac |
| pACYC184_CmOperon_pBP130_M6_Fwd_PacI | cggcgcgagccagtaccgct ttaattaa ttacgccccgccctgccact |
| pACYC184_CmOperon_pBP130_M6_Rev_SpeI | acccggcgcgtccgagctga actagt aacaggagggacagctgatagaaacaga |
| M6TE-SA-M6-RP | tcggacacctccggcgcgag |
| M6TE-SA-M6-FP | ggtggaggcgtcggcgttgc |
| M6TE-SA-130-FP | aacccagcagcaccgggccc |
| M6TE-SA-130-RP | ctggtgctgctgggcaggcg |
| DszATF1 | attttgtttaactttaagaaggagatatacatatgaaagcatacatgtttcccgggcaag |
| T7TerminatorR1 | ccctgcagcttaagttaattaacccggggcaaaaaacccctcaagacccgtttagag |
| LacCasetteLacI | attctcatgtttgacagcttatcatcgatactgcccgctttccagtcggg |
| LacCasetteLacO | cccttgcccgggaaacatgtatgctttcatatgtatatctccttcttaaagttaaacaaa |
| S86AMutationR | gaactcgcccagagcgtggccggccag |
| S86AMutationF | ctggccggccacgctctgggcgagttc |
| F190VMutationR | gaagcgggaatgaacagcggcgctcac |
| F190VMutationF | gtgagcgccgctgttcattcccgcttc |
| DszATMutant_CTerm | tggtggtggtggtgctcgagt |
| MalACP-F1 | ttttgtttaactttaagaaggagatatacatatgacgcaatttgcatttgtgttccctgg |

| | |
|---|---|
| MalACP-R1 | ggtggtggtgctcgagtgcggccgcaagcttgtcgacggagctcgaattaagctcgagcgccgctgccatcg ctgaagg |
| PresACP6_Fwd3 | catctagaagtgctttttcagggcccgcatatggcggccccggcgcgggagatgacgtcgcaggagt |
| PresACP6_Rev | tcctttcgggctttgttagcagccggatcctcagagctgctgtcctatgtggt |
| Cm-pT7-PhaEC-F | gggcgctgttcgaggcgtacg aacaggagggacagctgatagaaacaga |
| Cm-pT7-PhaEC-R | cggcgttgcggacgcgttcgaa ttacgccccgccctgccact |
| DEBS1-1 | gggcgctgttcgaggcgtacgggcgcgaccgcgagc |
| DEBS1-2 | ggcgatctcgccctgcgcgtgcccgatgac |
| DEBS1-3 | gtcatcgggcacgcgcagggcgagatcgcc |
| DEBS1-4 | cggcgttgcggacgcgttcgaacggccccgtcgcgacg |
| pACYC184_CmOperon_pBP130_M3_Fwd_PacI | tgcacgagcggccccgcgcg ttaattaa ttacgccccgccctgccact |
| pACYC184_CmOperon_pBP130_M3_Rev_SpeI | tggcgcggctgcaccacccc actagt aacaggagggacagctgatagaaacaga |
| pET21-M3-RP | caggcgaccgaggtgcacga |
| pET21-M3-FP2 | ccgaccggggctgggacctg |
| M5_BbvCI | tcagctcctccctcagctcg |
| M5_NsiI | ctgagcatgcatgggtctagaaataattttgtttaactttaagaaggagatatacatatgagcggtgacaacgg cat |
